# Comprehensive Lineage Tracing Maps the Landscape of Cell Fate Decisions in Mouse Embryogenesis

**DOI:** 10.64898/2026.05.07.722278

**Authors:** William N. Colgan, Luke W. Koblan, JoAnne Villagrana, Tien-Chi Jason Hou, Minming Wang, Gokul Gowri, Whitney Chandler, Leonardo A. Sepúlveda, Didar Ciftci, Karina Smolyar, Kathryn E. Yost, Alicia Young, Lars Wittler, Styliani Markoulaki, Kyle M. Loh, Xiaowei Zhuang, Nir Yosef, Zachary D. Smith, Jonathan S. Weissman

## Abstract

A comprehensive cell fate map of mammalian embryogenesis has remained out of reach given the scale, cellular diversity, and non-deterministic nature of development *in utero*. Here, we use PEtracer to continuously install heritable genetic marks as development progresses, reconstructing lineage trees that resolve ∼75% of cell divisions across >1.4 million cells from 16 replicate embryos. We pair these trees with deep transcriptional profiling to resolve cell fate biases, restriction timing, progenitor pool sizes, and lineage relationships throughout embryogenesis. Using this quantitative reference, we uncover strikingly reproducible lineage architecture between replicate embryos and chart the lineage dynamics driving fate specification across diverse tissues. Specific biological insights include the progressive restriction of neural crest fate, the relative contributions of distinct mesodermal origins to endothelium, and the dynamics of axial elongation. This work provides a foundation for a quantitative and predictive understanding of mammalian development.

## INTRODUCTION

Mammalian development is a non-deterministic process that reliably transforms a single fertilized cell into hundreds of distinct cell types, which are collectively organized into a conserved body plan^1^. A comprehensive cell fate map–the complete record of every progenitor-progeny relationship through which cellular diversity and morphogenetic complexity arise–would transform our understanding of this process^2^. Such a record would reveal how robust developmental outcomes emerge from probabilistic, context-dependent differentiation trajectories, define how cellular form and function are specified across successive divisions, and provide a quantitative blueprint for tissue assembly and patterning^3^.

The complete lineage map of *C. elegans*, reconstructed through direct observation of every cell division from zygote to adult, represents a landmark achievement in developmental biology^4^. Similar live cell tracking approaches in zebrafish and other transparent organisms have reconstructed lineage relationships and cell movements during embryogenesis^5–8^. However, these approaches have been limited in mammalian development by its scale, regulative complexity, and obligate *in utero* nature^8,9^. Instead, classical fate-mapping studies used vital dyes to label cells and track their progeny throughout development^10,11^. Subsequent advances, including recombinase-based genetic labeling and static DNA barcoding, expanded the repertoire of heritable markers and enabled cell type-specific labeling^2,12–17^, providing insights into how progenitor populations contribute to tissue patterning, as well as general descriptions of lineage potentials. In parallel, scalable single-cell profiling has revealed the diversity of cell types and states across embryogenesis^18–26^, but pairing this detailed molecular view with quantitative fate restriction dynamics has remained challenging.

Reconstructing complete cell fate restriction trajectories in mouse development requires continuous lineage tracing, sufficient recording capacity to resolve millions of divisions, and paired cell state and lineage readout from individual cells. Recently described prime editing-based systems build on earlier engineered lineage-recording approaches^27–41^ to enable high-content, evolving lineage recording while minimizing the formation of cytotoxic Cas9-induced double-stranded DNA breaks^42,43^. In particular, we developed PEtracer, which continuously installs predefined, heritable lineage marks (LMs) at genomically integrated lineage tracing cassettes (LTCs)^43–46^. PEtracer editing rates are fast enough to resolve individual cell divisions in rapidly dividing embryonic cells and can be tuned to capture lineage dynamics over longer timescales. Further, this system is compatible with both dissociative single-cell sequencing and imaging-based spatial readouts^47^, enabling scalable, joint measurement of lineage and cell state across modalities.

Here, we use PEtracer to chart cell state and lineage dynamics in early mouse embryogenesis. We generated high-grade chimeric embryos by introducing engineered mouse embryonic stem cells (mESCs) carrying the PEtracer system, including >100 editable sites, into host morulae. By performing single-cell RNA-seq (scRNA-seq) profiling of entire embryos at half-day intervals from E7.5 to E10.0, we determined each cell’s current state together with its lineage history. In total, we recovered >1.7 million cells across 16 embryos, with triplicate sampling at all but the final timepoint. By sampling approximately half of all cells and resolving ∼75% of divisions along recovered branches, we generated high-confidence trees that reveal the landscape of cellular fate decisions across the developing embryo. This comprehensive sampling approach allowed us to capture late cell divisions and fate commitment events that are systematically missed in subsampled datasets.

Building on more than a century of fate-mapping and lineage-tracing studies that established qualitative relationships between embryonic progenitors, we provide a quantitative view of cell fate dynamics across mouse gastrulation and early organogenesis. We measure progenitor populations, fate restriction timing, and clonal outputs, and show that these features are strikingly reproducible across replicate embryos. Analyses across specific cell types and tissues–including notochord specification, heart field separation, neural crest fate restriction, endothelial origins, and axial progenitors–illustrate how cell fates are jointly instructed by lineage, spatial position, and local signaling. Together, these data provide a lineage-resolved reference for interpreting and contextualizing focused studies of developmental cell fate specification. An interactive website (<u>mela.wi.mit.edu</u>) enables exploration of the lineage trees, gene expression patterns, and dynamics of fate restriction.

## RESULTS

### Enabling PEtracer studies of mouse development

To enable joint profiling of embryonic cell state and lineage history, we engineered the three PEtracer components into karyotypically normal mESCs: 1) a homozygous *Rosa26* knock-in of a Cre-activatable PEmax editor; 2) 35 unique genomically integrated LTCs, each containing a unique integration barcode (intBC) and three editable sites (ESs); and 3) pegRNA arrays (pegArrays) encoding 24 pegRNAs that install distinct 5-nucleotide LMs (8 per ES) (**Figures 1A, S1A, and S1B; Methods**). As development proceeds, PEmax continuously installs these predefined LMs at unedited ESs, generating heritable combinations of edits that encode each cell’s division history. *In silico* simulations indicate that the 105 editable sites in our mESCs are sufficient to reconstruct high-resolution lineage trees for cell populations >10^9^ cells, with appropriately tuned tracing kinetics^43^. We therefore created lines with pegArrays tuned for slow, intermediate, and fast editing using engineered protospacer mismatches, and confirmed expected tracing kinetics *in vitro* (**Figure 1B**).

**Figure 1.**
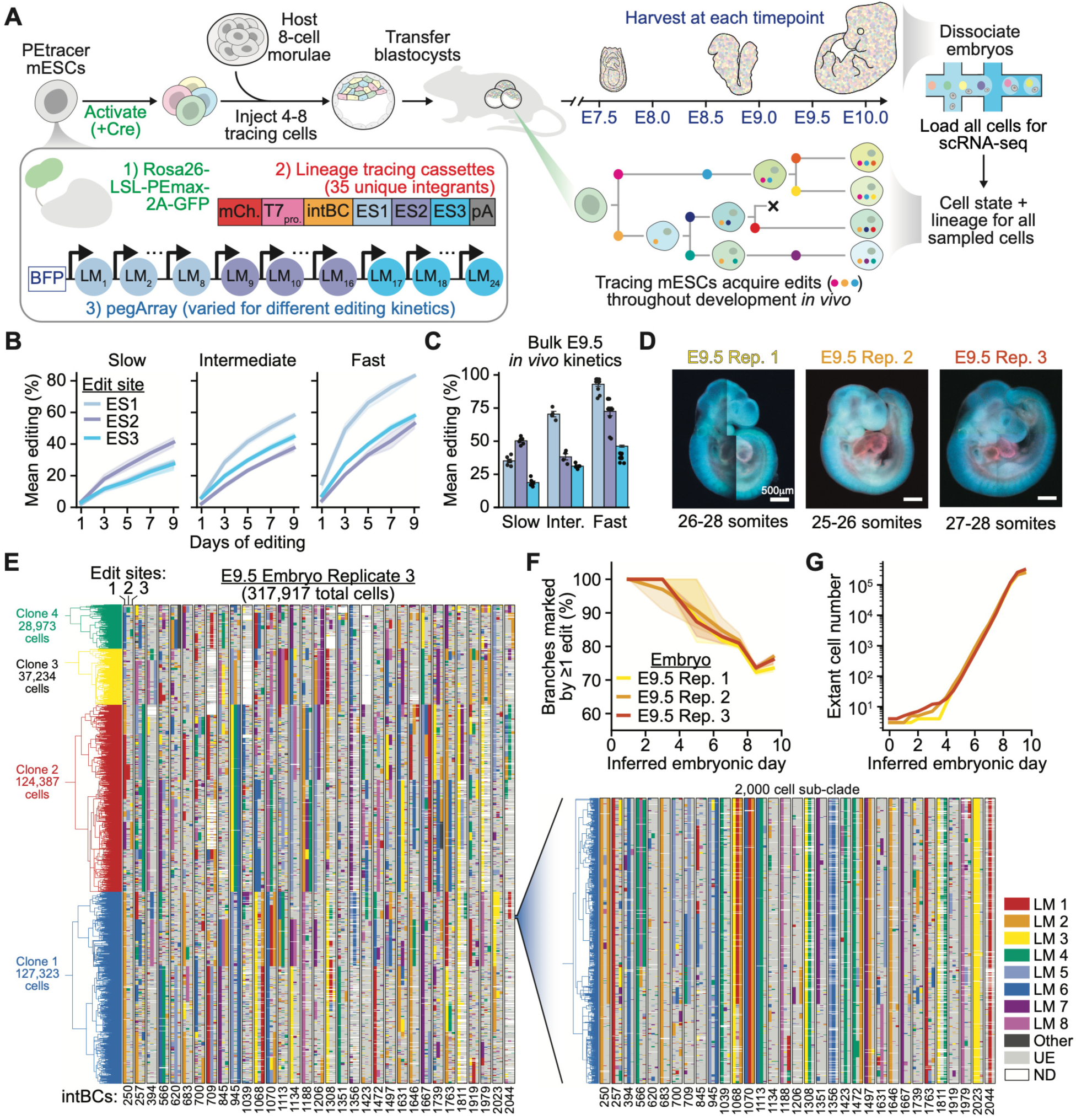
Embryonic-scale cell fate mapping of mouse development with PEtracer. (**A**) mESCs engineered with PEtracer components: Rosa26 knock-in of Cre-activatable PEmax-T2A-GFP, lineage tracing cassettes (LTCs) containing unique integration barcodes (intBCs) and three edit sites (ESs), and pegRNA arrays (pegArrays) encoding 24 lineage marks (LMs; eight per ES). Following PEmax activation, 4-8 donor mESCs were introduced into host 8-cell morulae and transferred into pseudopregnant recipients. LMs accumulate over time and are read out by scRNA-seq to reconstruct lineage trees. (**B,C**) Editing kinetics for slow, intermediate, and fast pegArray mESCs at ES 1-3 in (**B**) cultured mESCs and (**C**) in chimeric E9.5 embryos. Bars represent the mean of ≥4 biological replicates ± standard error of the mean (SEM). (**D**) E9.5 embryos before dissociation and scRNA-seq (green: PEmax-T2A-GFP, red: mCherry:LTC, blue: BFP-linked pegArray). Rep. = Replicate. (**E**) Lineage tree and LMs for 317,917 cells across four mESC clones comprising E9.5 Rep. 3, with a 2,000 cell sub-clade expanded. Columns are intBCs colored by LM identity for each ES; UE = unedited, ND = not detected. (**F**) Mean fraction of branches marked by ≥1 edit across three E9.5 embryo trees over inferred embryonic day; ribbons indicate 95% confidence intervals. (**G**) Estimated number of extant cells over inferred embryonic day based on branch length estimates for E9.5 replicate embryos.

We then generated chimeric embryos by introducing multiple Cre-recombinase-activated mESCs into host embryos at the 4-to-8-cell stage (∼E2-E2.5) (**Figure 1A**)^48^. Because all donor cells derive from a common engineered population, this approach yields highly reproducible tracing across embryos, while multiple founder clones provide independent lineage trajectories within each embryo and host-derived wild-type cells serve as an internal control for developmental competence. Chimeric embryos generated from all three pegArray lines were morphologically normal at E9.5, with high chimerism, *in vivo* tracing kinetics closely matching *in vitro* rates, and well-balanced LM installation frequencies across edit sites (**Figures 1C and S1C-G**).

We selected the intermediate kinetics line for all subsequent work in this study, as it provided substantial information content without saturation during the E7.5–E10.0 collection window. To validate this selection, we performed transcriptional profiling of two intermediate-kinetics E9.5 embryos, confirming expected contributions of tracing cells to inner-cell-mass-derived cell types and minimal bias between mESC clones composing these embryos (**Figures S1E and S1H-J**). We further validated our system by injecting activated PEtracer mESCs into tetraploid host morulae, which confirmed no obvious adverse effects of lineage tracing on embryogenesis both during our collection window and at later timepoints (**Figures S1K-M**).

### Comprehensive cell state and lineage profiling of early mouse embryogenesis

To chart cell state and lineage dynamics across early mouse development, we profiled 16 embryos at half-day intervals from E7.5 to E10.0–with triplicate sampling at all but the final timepoint (**Figures 1D and S2)**. This window spans late gastrulation, axis elongation, and early organogenesis, as the embryo grows from a few thousand cells to approximately one million^22,49–51^. To resolve late fate restriction events that subsampled lineage trees systematically miss, we dissociated each embryo in its entirety and loaded all cells for scRNA-seq profiling, achieving a mean cell recovery of 44% (1,342 to 382,254 cells/embryo; **Table S1**).

Guided by extensive *in silico* benchmarking, we reconstructed lineage trees using a scalable heuristic neighbor-joining approach (**Figures S3A-I**; Methods). These trees revealed high-quality lineage tracing, with a 94% LM detection rate, 98% of cells carrying a unique edit profile, and ∼75% of branches marked by ≥1 edit (**Figures 1E**, **1F, S3J, and S3K**). By inferring division timing from the number of edits accumulated along each branch–where more edits indicate longer intervals between divisions–we obtained highly reproducible estimates of developmental time (**Figures 1G and S3L**). The inferred number of extant cells tracked both published estimates and measured cell counts in this study, with plateaus at the end of each growth curve consistent with ∼50% cell loss during dissociation and capture^50,52^. PEmax expression was stable across trees (Moran’s *I* = 0.14) and showed minimal variation by cell type; LM edit fraction likewise showed no systematic cell-type dependence and was unrelated to PEmax expression or cell cycle phase (**Figures S3M–P**). These results do not preclude increased edit rate variability in more differentiated cell populations at later timepoints^53^.

Across 16 embryos, we recovered 1,792,520 high-quality cells at a mean depth of 16,847 UMIs per cell (**Figure 2A; Table S1**). Transcriptional states were highly consistent across embryos collected at the same timepoint, with minimal batch effects, and integration with existing atlases confirmed expected cell type distributions and enabled precise transcriptional staging of each embryo alongside our own morphological assessment, revealing expected variation of up to a few hours within each timepoint (**Figures S2 and S4A-H**)^21,22,54^. We resolved 99 transcriptionally distinct cell types and 142 subtypes spanning embryonic and extraembryonic tissues, including 24 neuron subtypes and rare populations such as primordial germ cells (**Figures 2A-C, S5, and S6; Table S2**).

**Figure 2.**
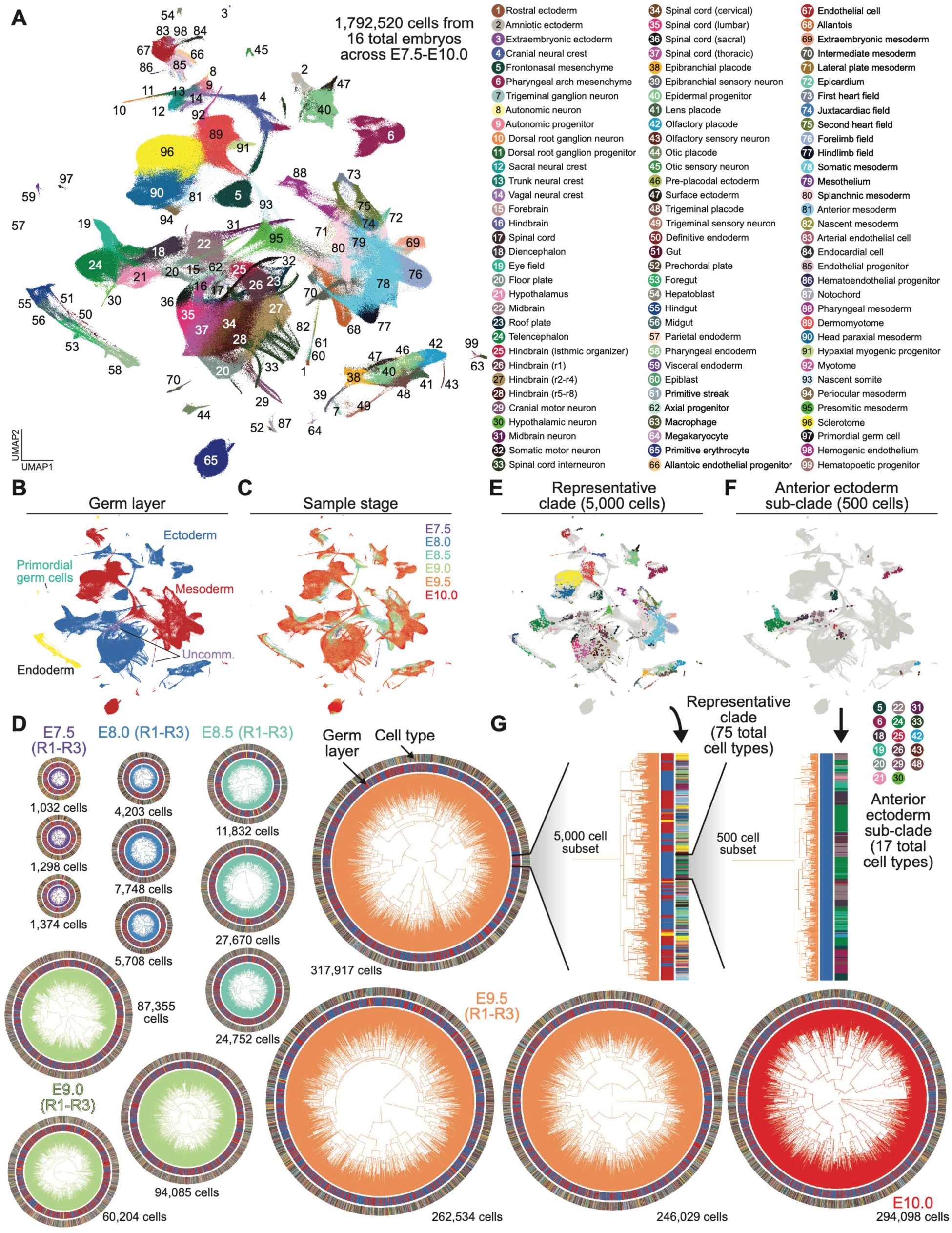
Whole-embryo joint cell state and lineage atlas of early mouse embryogenesis. UMAP of 1,792,520 cells from 16 embryos spanning E7.5–E10.0, colored by (**A**) cell type, (**B**) germ layer assignment (ectoderm, mesoderm, endoderm, primordial germ cells, and uncommitted), and (**C**) embryonic stage. (**D**) Lineage trees colored by sample stage. Rings indicate cell type and germ layer assignment; cell number is indicated for each tree. R=Replicate. (**E,F**) UMAPs colored by cell type for a representative 5,000-cell clade from E9.5 replicate 3 and a 500-cell anterior ectoderm restricted sub-clade. (**G**) Corresponding lineages of the clades in (E) and (F).

Chimerism rates ranged from 52-99% across embryos, with host and donor cells reproducibly populating near-identical transcriptional states in each embryo, suggesting that PEtracer does not perturb normal embryonic cell fate specification (**Figures S4I and S4J; Table S1**). As expected, donor PEtracer mESCs showed minimal contribution to extraembryonic endoderm and ectoderm (**Figures S4K and S4L**)^55,56^. In total, we reconstructed trees for 1,447,839 donor cells, capturing both pre- and post-gastrulation dynamics, with broad germ layer structure evident at whole-embryo scale and progressively finer sub-clades resolving to increasingly lineage-restricted groups (**Figures 2D-G**).

### Quantifying germ layer cell fate restriction dynamics

By pairing high-resolution lineages with transcriptional state across >1.4 million cells, we sought to dissect the dynamics of cell fate restriction. Here, we define cell fate operationally as the observed distribution of descendant cell types, rather than as an inference of developmental potential^3,26,57^. Examining ancestral nodes within our reconstructed trees, we asked when fates became restricted, whether biases emerged prior to full specification, and how the number and output of fate-restricted clades changed over time.

Beginning with germ layer specification, we calculated a fate distribution of each internal node based on the germ layer composition of its descendants (**Figure 3A**). Ancestral fate distributions were initially consistent with pluripotency, with nodes producing all three germ layers, before becoming increasingly biased during the window associated with gastrulation (∼E6.5 to ∼E7.5) (**Figure 3B**). To quantify these dynamics, we defined a fate bias index based on the normalized entropy of each node’s fate distribution (**Methods**), which revealed highly reproducible specification trajectories across timepoints and embryos (**Figure 3C**). Weak, yet statistically significant, germ layer fate bias first appeared at ∼E4.0–around the time of implantation but before the onset of gastrulation–suggesting that spatial position and signaling cues in the epiblast subtly influence fate prior to differentiation (**Figures 3C and S7A**).

**Figure 3.**
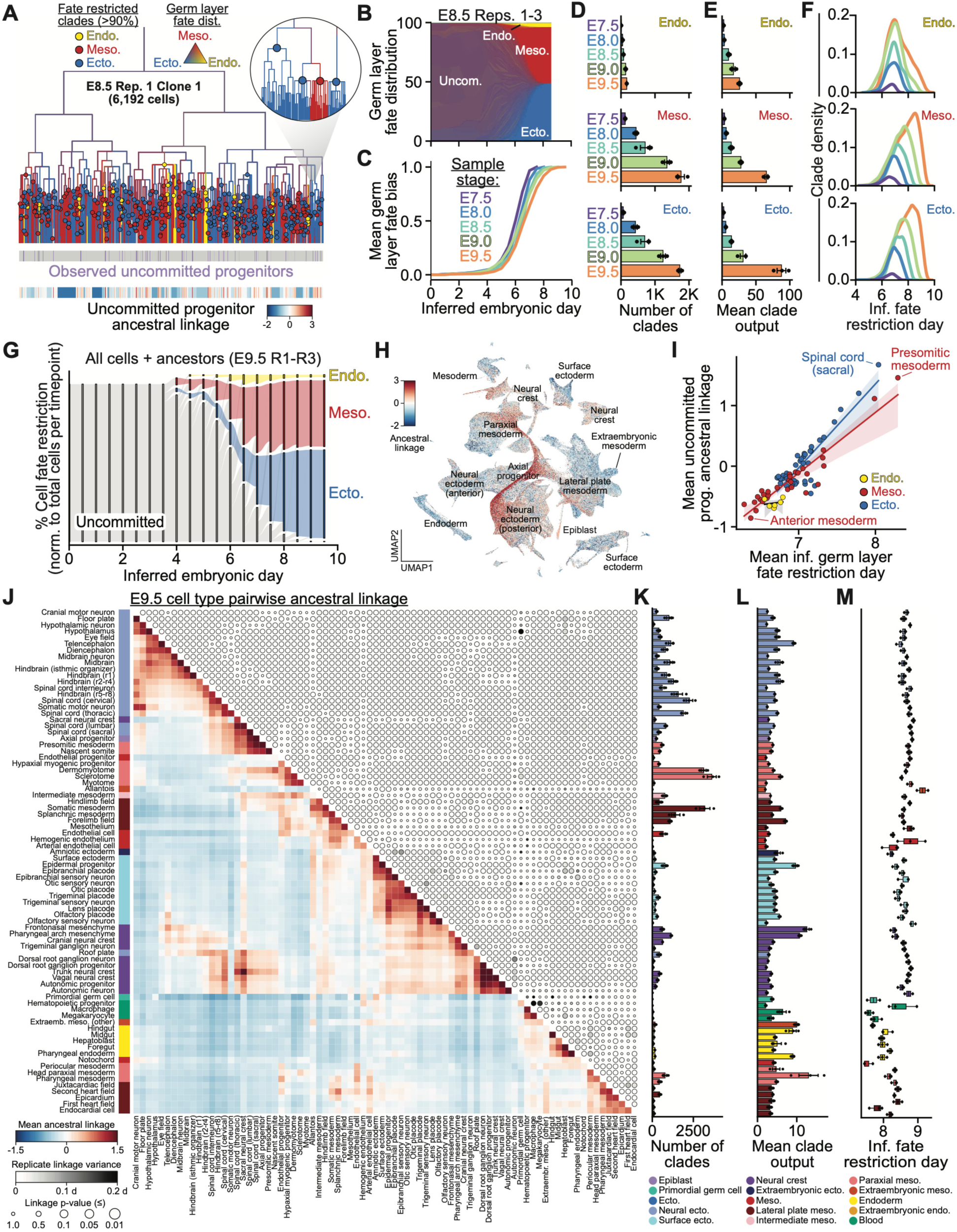
Cell fate bias, pool size, and fate restriction timing by germ layer and cell type. (**A**) Representative lineage tree for E8.5 replicate (rep.) 1 clone 1 (6,192 cells). Fate-restricted nodes (>90% representation from one germ layer) are colored: yellow = endoderm (Endo.), red = mesoderm (Meso.), blue = ectoderm (Ecto.). Branches are colored by descendant fate distribution. Inset sub-tree highlights these metrics at higher resolution. Color bars below indicate observed uncommitted progenitors and their ancestral linkage to other cells. (**B**) Relative abundance of germ layer fate distributions over time across triplicate E8.5 embryos. (**C**) Mean germ layer fate bias for triplicate embryos from E7.5-E9.5, calculated as one minus the normalized entropy of descendant fate distributions at each node relative to root. (**D**) Number of fate-restricted clades per germ layer by timepoint. (**E**) Mean number of descendant cells per fate-restricted clade. For (D,E), bars represent mean ± SEM for three biological replicate embryos. (**F**) Distribution of inferred germ layer fate restriction times across sampled stages, shown as a kernel density estimate over clades (area under each curve = 1). Curves are colored by sampled stage. (**G**) Germ layer fate restriction over time for all E9.5 cells and their ancestors across reps. 1-3. (**H**) UMAP of all sampled cells colored by ancestral linkage (**Methods**) to uncommitted progenitors. (**I**) Mean ancestral linkage of each cell type to uncommitted progenitors at E9.5 versus mean inferred (inf.) germ layer commitment timing. Points colored by germ layer assignment. (**J**) Mean pairwise ancestral linkage across all cell types. Axis color blocks indicate developmental domain; circle size indicates permutation test p-values; circle shading indicates variation between replicates. (**K**) Count, (**L**) mean number of descendants, and (**M**) commitment timing of cell type-restricted clades, colored by developmental domain. Error bars indicate ± SEM.

We next defined ancestral nodes for which >90% of descendants belonged to a single fate–here, any one germ layer–as “fate-restricted” clades (**Figure 3A**). Across timepoints, both clade number and clade size increased, particularly for ectoderm and mesoderm, with a consistent number of fate-restricted nodes at a given tree depth across triplicate embryos from E7.5 to E9.5 (**Figures 3D-F and S7B**). On average, endodermal fate restriction occurred earliest, around E7.0, whereas ectodermal and mesodermal specification often occurred later and over a broader interval, driven by continued emergence of clades contributing to a subset of cell types (**Figures 3F**, **3G, and S7C**)^15^.

To understand why germ layer fates are specified over different developmental windows, we asked whether uncommitted progenitors–defined transcriptionally as epiblast cells at E7.5 and axial progenitors at later timepoints–remain preferentially linked to specific germ layers over time. We defined an ancestral linkage metric based on the lineage distance from any source cell to the nearest member of a target population, normalized against a permuted background expectation, where positive values indicate more recent than expected shared ancestry (**Methods**)^28,57^. Germ layer fate-restriction timing correlated strongly with linkage to uncommitted progenitors and showed clear anterior-posterior organization across mesodermal and ectodermal outputs, with posterior cell types (*e.g.,* presomitic mesoderm, sacral spinal cord) sharing more recent ancestry with uncommitted progenitors and anterior populations separating earlier (**Figures 3H**, **3I, and S7D**).

Finally, we characterized how these analyses were affected by chimerism rate and confirmed their robustness to technical error. Lower-chimerism embryos yielded fewer fate-restricted clades, as expected from their smaller proportion of traced cells (**Figure S7E**). We observed a germ layer mismatch rate of ∼1%, estimated as the frequency with which a cell’s label disagreed with the most parsimonious label for its parent, indicating that misplaced cells are rare (**Figures S7F and S7G**). Clade number, output, restriction timing, and ancestral linkage were robust to this level of simulated topological error and to integration dropout assessed by leave-one-integration-out jackknife analysis (**Figures S7H–O**).

### Lineage architecture of cell type diversification

Having established the dynamics of germ layer specification, we asked how individual cell types emerge from these broad developmental compartments. We computed mean pairwise ancestral linkage across 80 abundant cell types at E9.5, generating a matrix that captures patterns of shared ancestry (**Table S3**). Clustering revealed blocks of closely related cell types, as well as additional structure within and between blocks that reflects lineage history and spatial patterning across the embryo (**Figure 3J**; see **Figures S5 and S6** for cell subtype-level linkage analysis). These relationships were remarkably consistent (<6 hour variation) across triplicate E9.5 embryos.

Neural ectodermal cell types formed a large cluster structured primarily by position along the anterior-posteior axis (**Figure 3J**). Anterior populations–including forebrain, midbrain, and hindbrain–were strongly linked, while posterior spinal cord populations generated during axial elongation^15,58,59^ occupied a distinct cluster. Differentiated neurons clustered by their regional identities, with spinal cord interneuron and motor neuron subtypes further organized by dorsal-ventral (D-V) position from floor to roof plate (**Figure S6D**). Surface ectoderm and neural crest (NC) each formed discrete yet tightly associated clusters, with NC also linked to the neural ectoderm, consistent with its origin at the boundary between neural and surface ectoderm in the developing neural plate^60–64^. Within the surface ectoderm, sensory neurons clustered tightly with their associated placodal populations, consistent with recent reports^65^.

Mesodermal cell types segregated into two distinct blocks: an anterior block comprising head, pharyngeal, and cardiac mesoderm, and a posterior block containing lateral plate and paraxial derivatives including dermomyotome and sclerotome (**Figure 3J**). Endothelial cells were associated with both the anterior and posterior clusters, reflecting their broad mesodermal origins^66^. Further, among posterior cell types, ancestral linkage organizes paraxial, intermediate, and lateral plate derivatives along the mediolateral axis. Notably, the most posterior mesodermal populations–presomitic mesoderm and nascent somites–were more strongly linked to axial progenitors and neural ectoderm than to other mesodermal cell types, consistent with axial progenitors contributing to both neural and mesodermal lineages during axis elongation^58,59^.

Finally, a subset of cell types showed minimal off-diagonal linkage (**Figure 3J**). Endodermal cell types formed a cohesive cluster organized along the A-P axis, in line with the early lineage separation observed in our germ layer analyses. Notably, notochord cells exhibited linkage to both endoderm and floor plate, despite their classification as mesoderm, reflecting their unique developmental origin and signaling role^67,68^.

### Quantification of cell fate specification

To resolve how cell fate specification unfolds over time, we asked when fate biases toward specific cell types first emerge and when these fates become restricted. We first quantified how ancestral fate distributions in E9.5 lineage trees deviated from background over time, and found that weak fate bias became detectable at ∼E4.0, a result we corroborated using a cell type label-independent measure of transcriptional distance (**Figures S8A and S8B**). To dissect the drivers of these early biases, we clustered ancestral nodes by their descendant fate distribution and identified 15 recurrent fate bias patterns present before E7.0 with the earliest biases driven primarily by mesodermal fates and some anterior ectodermal and NC cell types (**Figures S8C-I**).

We next quantified when these biases resolve into fate-restricted clades, the number of clades contributing to each cell type, and the overall output of these clades (**Table S4**). Among the earliest fate-restricted cell types were extraembryonic mesoderm and extraembryonic-derived megakaryocytes and macrophages, as well as primordial germ cells, which arise as a specific population within the posterior embryo at the earliest stages of epiblast patterning (**Figures 3K-M**)^69,70^. Because of their early specification, these populations showed minimal linkage to other developmental domains (**Figure 3J**). Clade statistics, cell type abundances, and early fate bias patterns were all highly consistent across independent embryos, indicating that progenitor timing, number, and differentiation trajectories are all stereotyped features of mammalian development (**Figures 3K-M and S8J-M**).

### Early fate restriction of the notochord

The notochord is a transient axial structure that forms along the embryonic midline during gastrulation and functions as a key signaling center for D-V patterning of the overlying neural tube and adjacent paraxial mesoderm^71^. Classical fate-mapping studies revealed how the notochord arises from organizer regions at the anterior end of the primitive streak, where cells display marked fate plasticity, and shares clonal relationships with both the definitive endoderm and the ventral neural tube^10,67,72,73^. However, these relationships have largely been inferred from sparse clonal labeling studies, leaving the relative contribution and timing of the notochord’s shared ancestry across germ layers unquantified.

Consistent with its early morphogenesis, the notochord was among the earliest-specified cell types in our dataset, with an average fate-restriction time before E7.5 across all triplicate embryos–often concurrent with the initial specification of mesodermal fate (**Figure S9A**). To resolve its developmental origins, we quantified the sibling outputs of notochord-restricted clades in our lineage trees (**Figure 4A; Methods**). The majority of sibling descendants were mesodermal, but notably 25.5% were endodermal, a 13.4-fold enrichment relative to their overall frequency (**Figures 4B and 4C**). At the cell type level, sibling outputs were enriched for gut and paraxial mesoderm, tissues that develop alongside the notochord at the embryonic midline, while ectodermal sibling outputs were predominantly floor plate (**Figure S9B**).

**Figure 4.**
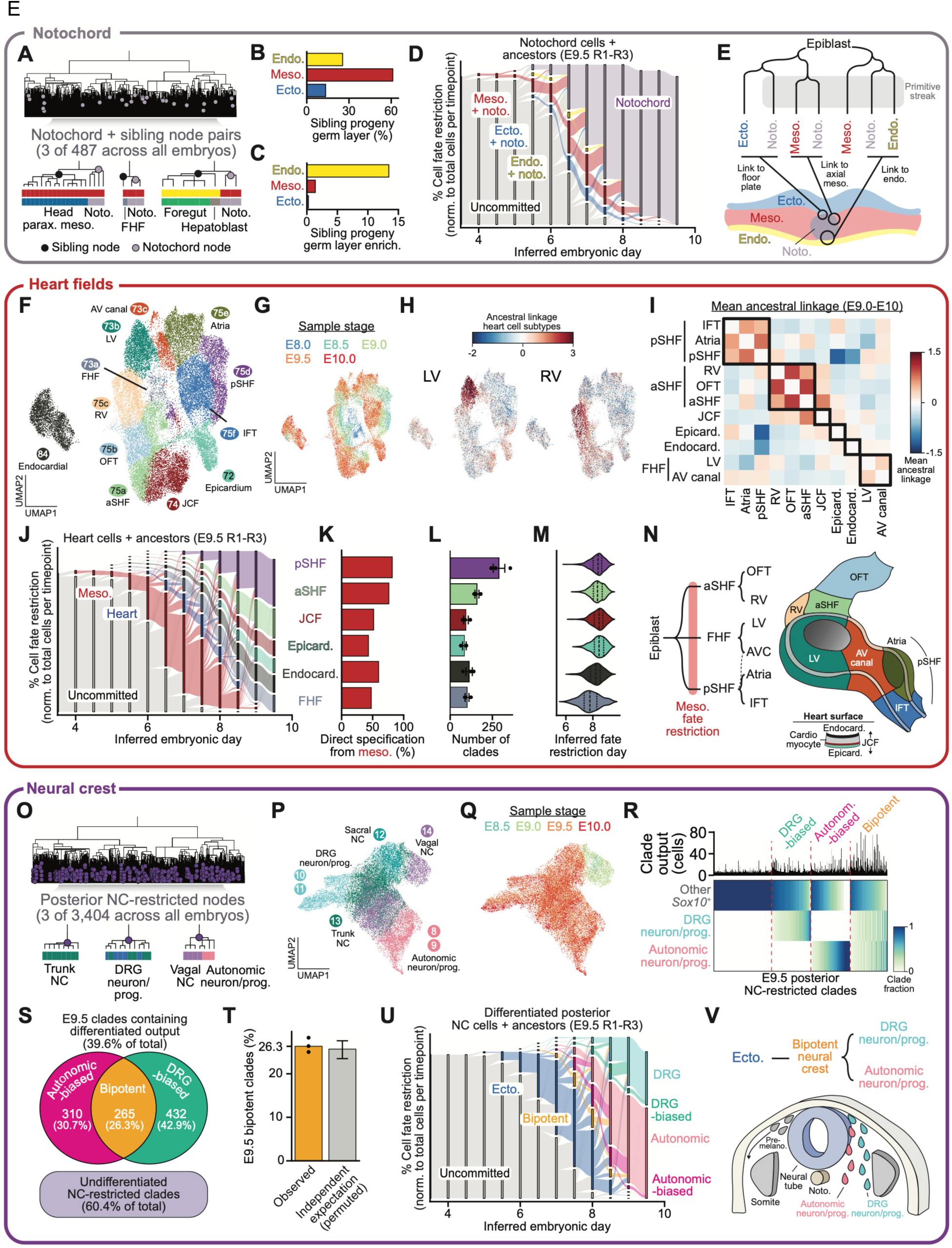
Cell fate specification dynamics in the developing notochord, heart fields, and neural crest. (**A-E**) <u>Notochord:</u> (**A**) Example tree (E9.5 replicate 2, clone 1; E9.5-R2-C1) with notochord-restricted nodes marked (grey circles). Insets show 3 of 487 notochord+sibling (black circles) pairs and descendant cell types. (**B**) Germ layer composition and (**C**) enrichment relative to background of notochord sibling clades. (**D**) Cell fate restriction over time at the level of germ layer and notochord for all notochord cells and their ancestors across E9.5 reps. 1-3. Notochord cells treated as wildcards for germ layer fate restriction. (**E**) Model of direct notochord specification from epiblast, with siblings adopting alternative fates around primitive streak ingression. (**F-N**) <u>Heart fields:</u> UMAP of 12 cardiac subtypes from E8.0-E10.0 colored by (**F**) cell type, (**G**) stage, and (**H**) ancestral linkage to left ventricle (LV) and right ventricle (RV). FHF = first heart field, AV canal = atrioventricular canal, aSHF/pSHF = anterior/posterior second heart field, IFT = inflow tract, JCF = juxtacardiac field, OFT = outflow tract. (**I**) Mean pairwise ancestral linkage across cardiac cell types (E9.0-E10.0). Boxes mark FHF, aSHF, and pSHF. Epicard. = epicardium, Endocard. = endocardium. (**J**) Cell fate restriction over time at the level of mesoderm, heart, and heart field for all heart cells and their ancestors across E9.5 reps. 1-3. (**K**) Percentage of clades directly specified from the mesoderm by heart field. (**L**) Count and (**M**) commitment timing of heart field-restricted clades. Error bars indicate ± SEM. (**N**) Model of sequential FHF, aSHF, and pSHF restriction from distinct mesodermal progenitors. (**O-V**) <u>Neural crest (NC):</u> (**O**) Example lineage tree (E9.5-R2-C1) with posterior NC-restricted nodes (purple circles) marked. Inset shows 3 of 3,404 nodes, color bars indicate descendant cell identities. DRG = dorsal root ganglia, prog. = progenitor. UMAP of posterior NC cells colored by **(P)** cell type, and (**Q**) stage. (**R**) Outputs of E9.5 posterior NC-restricted clades, grouped as DRG-biased, autonomic-biased, or bipotent; clade size shown above. (**S**) Venn diagram of outputs in (R). (**T**) Percent bipotent clades compared to expectation given statistically independent specification (clade-size-stratified permutation). Error bar indicates 95% CI from permutation test. (**U**) Cell fate restriction over time at the level of ectoderm, neural crest (DRG-biased, autonomic-biased, or bipotent), and differentiated neural crest (DRG and Autonomic) for all differentiated E9.5 NC cells and their ancestors across E9.5 reps. 1-3. (**V**) Model of bipotent trunk NC cells resolving to DRG or autonomic fates.

These data support a model in which the notochord is often specified directly from epiblast cells, with transitory shared ancestry across germ layers reflecting alternative fates adopted by sibling descendants during and after primitive streak ingression according to their spatial position (**Figures 4D and 4E**). The endodermal enrichment aligns with classical fate-mapping linking these populations^10,73^, and with their transcriptional similarity during the E7.5–E8.0 window (**Figures S9C and S9D**).

### Lineage architecture of the developing heart

The developing mouse heart arises from temporally and spatially patterned progenitors. The first heart field (FHF) contributes to the early heart tube and predominantly gives rise to the left ventricle (LV), whereas second heart field (SHF) progenitors progressively add to the arterial and venous poles to elongate the heart^74–76^. Fate-mapping studies have resolved these relationships and identified further subdivisions including distinct anterior (aSHF) and posterior (pSHF) domains of the SHF, primarily associated with right ventricle (RV) / outflow tract and atria / inflow tract, respectively^77–81^. However, these studies have largely relied on static barcoding approaches with distinct labeling efficiencies and dynamics, making it difficult to directly compare fate restriction timing, progenitor overlap, and relative contributions to different cardiac structures.

To address these questions, we annotated 12 cardiac cell subtypes spanning the major cardiac compartments from E8.0-E10.0 (**Figures 4F**, **4G, and S9E**). We then computed ancestral linkage between each population and all other heart cells, revealing lineage relationships not apparent from transcriptional data alone (**Figures 4H and S9F**). For example, despite their transcriptional similarity (*e.g., Myh7*, *Myl3* expression), RV and LV populations were not closely related, matching their aSHF and FHF origins^76,82^. Mean pairwise ancestral linkage cleanly separated the FHF, aSHF, and pSHF, with minimal linkage between the aSHF and pSHF (**Figure 4I**)^83–86^. It also linked atrioventricular canal cells to both the FHF and pSHF, reflecting their position at the atrioventricular boundary^87^, and cleanly segregated epicardial and endocardial cells^79,88^.

The early lineage separation of aSHF and pSHF prompted us to examine the dynamics of heart field specification. Tracing heart cells back through inferred time and annotating the fate-restriction of their ancestors revealed that heart field restriction frequently coincided with mesoderm restriction, with ∼50% of clades specified directly from the mesoderm rather than passing through a heart-restricted intermediate, a result also supported by quantifying heart field co-occurrence within heart-restricted and mesoderm-restricted clades (**Figures 4J**, **4K, and S9G–J; Methods**)^76,82,89^. The number of contributing clades and their restriction timing varied by heart field: the FHF comprised a small number of early-specified clades, whereas the pSHF comprised a larger number of later-specified clades (**Figures 4L and 4M**). Together, these data align with conventional marker-driven lineage tracing results demonstrating that distinct lineage identities are established alongside mesodermal fate restriction, well before transcriptionally defined heart cell types are detected (**Figure 4N**)^76,82,89^.

Finally, we asked whether lineage-related structure defines identities below the level of heart field. Descendants of 38.7% of heart-restricted clades were significantly more transcriptionally similar to one another than were randomly sampled cells of the same cell type, revealing sub-domains nested within broader cardiac identities (**Figures S9K and S9L**)^90^. Many transcriptionally diverse clades contained one or more endocardial cells, in agreement with the emergence of this cell type from multiple heart fields (**Figure S9M**)^91^. Overall, 68.7% of endocardial cells derived from endocardial-biased clades, while the rest derived from clades biased toward other heart fields, particularly the FHF (**Figures S9N and S9O**).

### Fate restriction in the neural crest

The neural crest is a transient, migratory population of multipotent progenitors that generates diverse structures across the embryo, with the trunk NC giving rise to much of the peripheral nervous system^92–94^. Competing models hold that trunk NC progenitors are either multipotent, generating both sensory and autonomic (sympathetic and parasympathetic) derivatives, or prespecified when they emerge^95–97^. To distinguish these possibilities, we identified 3,404 posterior NC-restricted clades containing differentiated dorsal root ganglion (DRG) and autonomic neurons alongside their progenitors (**Figures 4O-Q**). Vagal, trunk, and sacral NC showed sequential fate restriction, tracing a progressive wave of specification along the A-P axis (**Figure S9P**)^62,63^. Clades from early-sampled embryos were predominantly composed of *Sox10*^+^ NC progenitors, while E9.5 clades additionally contained DRG and autonomic outputs (**Figure S9Q**).

We classified clades with differentiated output at E9.5 as DRG-biased, autonomic-biased, or bipotent based on the composition of descendant cells (**Figures 4R, S9R, and 9S**). These annotations represent observed fates rather than potential, as the remaining *Sox10*+ progenitors may later differentiate into other cell types. Among clades with differentiated outputs, 42.9% were DRG-biased, 30.7% were autonomic-biased, and 26.3% were bipotent (**Figure 4S**). Notably, after controlling for the number of differentiated cells per clade, the observed bipotent fraction matched that expected under independent assortment of DRG and autonomic fates (**Figure 4T**). Earlier-restricted clades were more likely to contain a differentiated output, as expected given maturation over developmental time (**Figure S9T**).

Reconstructing differentiated NC fate trajectories from ancestral fate biases revealed that bipotent progenitors tended to be short-lived, rapidly resolving to DRG- or autonomic-restricted clades between ∼E7.5 and ∼E9.0, coincident with trunk neural crest migration (**Figure 4U**). Together, these data are compatible with a model in which commitment to DRG and autonomic fates is not prespecified at the time of trunk NC specification, but resolves over the course of NC migration (**Figure 4V**)^62^. However, since delamination is not directly measured, the precise timing of fate restriction relative to this process cannot be determined.

### Gene expression drivers of cellular diversification

Having resolved the lineage architecture underlying cell fate specification, we next sought to understand the gene regulatory programs that drive these transitions. We identified genes with heritable expression patterns and grouped them into 70 coherent lineage-associated programs (**Figures 5A and S10A; Methods**)^98^. These programs captured major axes of developmental regulation, including spatial patterning, cell type specification, and signaling responses (**Figures 5A, S10B, and S10C**). We assessed gene program activity across all cells and timepoints and imputed activity at ancestral nodes (**Methods**), enabling reconstruction of program dynamics over time (**Figures 5B**, **5C, S10D-G, and S11**).

**Figure 5.**
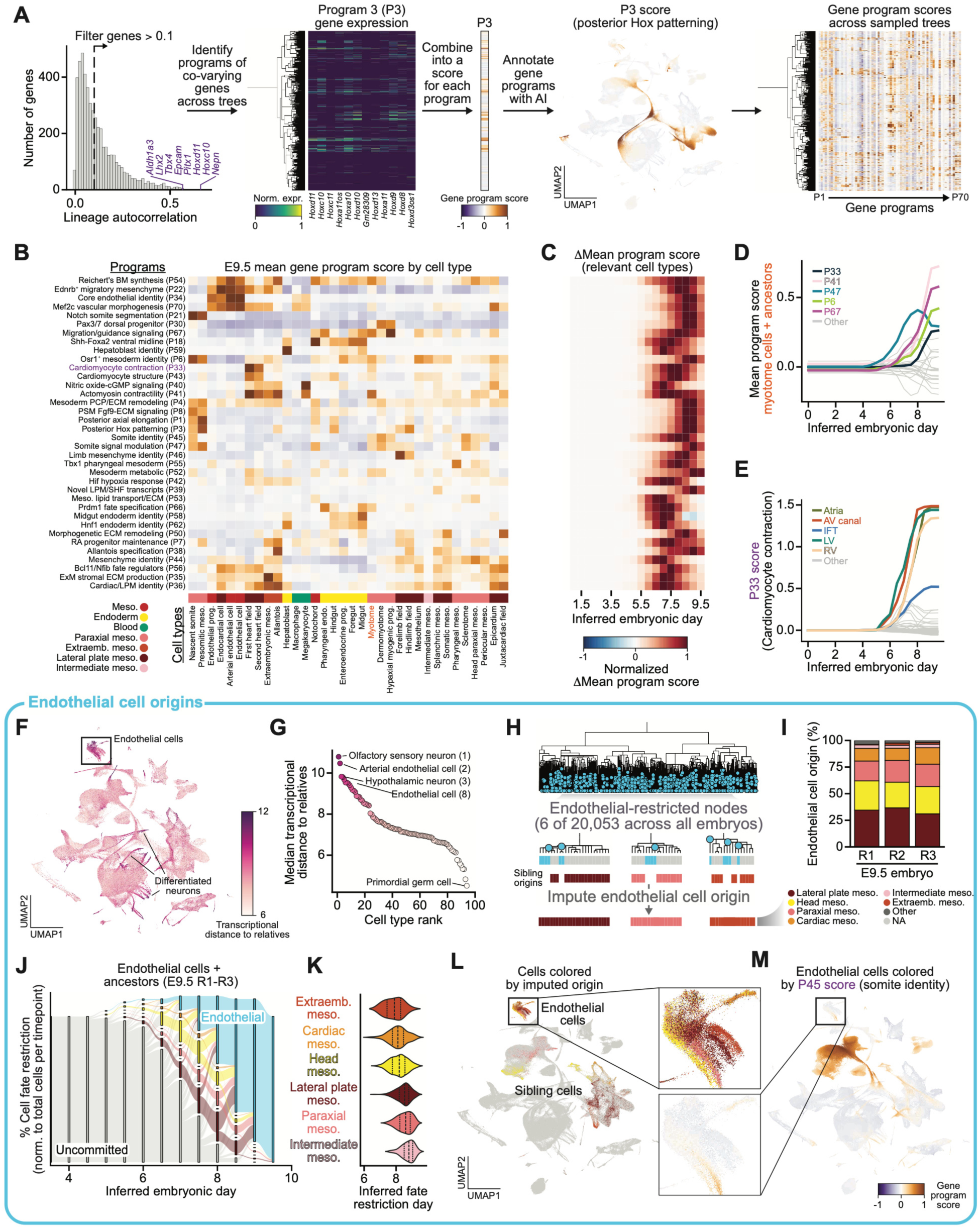
Gene regulatory programs underlying cell fate specification. (**A**) Gene program identification process. Genes with heritable expression (scaled local autocorrelation >0.1) were grouped into 70 programs by pairwise local autocorrelation^98^. Program 3 (P3; posterior Hox patterning) is shown as an example, including normalized expression of constituent genes, score across an example lineage tree (E9.5-R2-C3), and its score on the UMAP. All 70 program scores are shown on the tree at right. (**B**) Mean program score across mesodermal and endodermal cell types in E9.5 embryos. Colorbar below indicates developmental domain. Program annotation details described in **Table S5**. (**C**) Change (Δ) in mean program score over inferred embryonic day in E9.5 lineage trees for relevant cell types (mean >0.5 in ≥1timepoint), normalized per program. (**D**) Mean program score over inferred embryonic day for selected programs within the myotome lineage: P33 (cardiomyocyte contraction), P41 (actomyosin contractility), P47 (somite signal modulation), P6 (*Osr1*+ mesoderm identity), and P67 (migration/guidance signaling). (**E**) P33 score by inferred embryonic day across cardiac cell types (colored) and all other cell types (grey). (**F-M**) <u>Endothelial origins:</u> (**F**) UMAP of all sampled cells colored by mean transcriptional distance to their closest relatives (MRCA ≤1.5 days). (**G**) Medain transcriptional distance to relatives ranked across cell types. (**H**) Endothelial origin imputation: E9.5-R2-C1 tree with endothelial-restricted nodes marked (blue circles). Insets show 6 of 20,053 fate-restricted nodes and siblings. Cell type composition of siblings used to infer the developmental origin of each endothelial cell. (**I**) Proportion of endothelial cells assigned to lateral plate, intermediate, paraxial, extraembryonic, head, and cardiac mesoderm origins for E9.5 embryo reps. 1-3. (**J**) Cell fate restriction over time at the level of mesoderm type and endothelial for all endothelial cells and their ancestors across E9.5 reps. 1-3. Endothelial cells are treated as wildcards for mesoderm type fate restriction. (**K**) Endothelial commitment timing by inferred mesodermal origins. (**L**) UMAP of all cells with endothelial cells and their siblings colored by inferred developmental origin, inset shows endothelial cells and (**M**) P45 (somite identity) score.

These analyses revealed two complementary modes of gene program activity: lineage-restricted deployment within specific cell types and recurrent activation across them. Within individual lineages, gene programs showed temporally ordered activation. In the myotome, somite signal modulation (P47) preceded actomyosin contractility (P41), while cardiomyocyte contractility (P33) was activated earliest in the FHF and later across other cardiac cell types (**Figures 5D and 5E**). In contrast, other programs were reused across developmentally distinct cell types–for example, the mesenchymal program (P44) was deployed across both NC- and mesoderm-derived mesenchymal populations (**Figures 5B and S10D**).

The rapid activation of neuronal programs across ectodermal lineages (**Figures S10D-G**) suggested that some cell types undergo sharp transcriptional transitions over short developmental intervals, including the activation of convergent functional programs from independent progenitor pools. To quantify this phenomenon, we calculated the mean transcriptional distance between cells and their closest lineage relatives (**Methods**). We saw clear transcriptional divergence of neural subtypes from their progenitors, as well as notable signals from endothelial cells, consistent with their rapid deployment throughout the embryo to form the vasculature (**Figures 5F and 5G**)^99^.

### Multiple parallel developmental origins of endothelial cells

The prevailing view is that most endothelial cells (ECs) arise from lateral plate mesoderm^100,101^, but alternative sources have also been proposed, including paraxial^102–105^, and intermediate mesoderm^106^. As past studies generally focused on one developmental origin at a time, we sought to systematically trace the parallel contribution of multiple mesodermal tissues to ECs, quantifying the percentage arising from each source.

We identified 20,053 endothelial-restricted clades across all trees and inferred their developmental origins from their siblings (**Figure 5H**). Consistent with the prevailing model, the lateral plate mesoderm was the largest contributor, accounting for 34.3% of ECs^100,105^. Notably, head (22.3%), paraxial (14.5%), extraembryonic (11.8%), cardiac (10.6%), and intermediate (2.2%) mesoderm also generated ECs (**Figure 5I**)^91,103–107^. These proportions were consistent across the three E9.5 replicate embryos. Endothelial fate restriction varied systematically by developmental origin, with extraembryonic- and cardiac-derived ECs specified earliest (**Figures 5J and 5K**). Further, sibling compositions within each compartment reflected their developmental context, including enrichment of hypaxial myogenic progenitors among siblings of paraxial-derived ECs (**Figure S10H**)^104,108^.

This diversity of origins is a clear example of convergent differentiation^2^, raising the question of whether ECs from different sources are transcriptionally identical. ECs from all mesodermal sources upregulated genes in the core endothelial program (P34), forming a single cluster in the UMAP embedding (**Figures 5L and S10I**), but also retained detectable transcriptional signatures of their tissue of origin (**Figure S10J)**. For example, paraxial-derived ECs maintained residual expression of the somite identity program (P45), consistent with *in vitro* evidence indicating that intermediate differentiation trajectories shape the final transcriptional profiles of ECs (**Figure 5M**)^109^.

### Spatiotemporal patterning of the neural ectoderm

Spatial patterning is a central driver of cell type specification in the developing embryo, particularly within the neural ectoderm, which generates diverse neuronal populations along the A-P and D-V axes^61,110,111^. To understand the interplay between spatial position and cell fate, we used published spatial transcriptomics data to identify spatially predictive genes and inferred each cell’s A-P and D-V position from their expression, restricting D-V analysis to the spinal cord (**Figures 6A-E and S12; Methods**)^112–114^.

**Figure 6.**
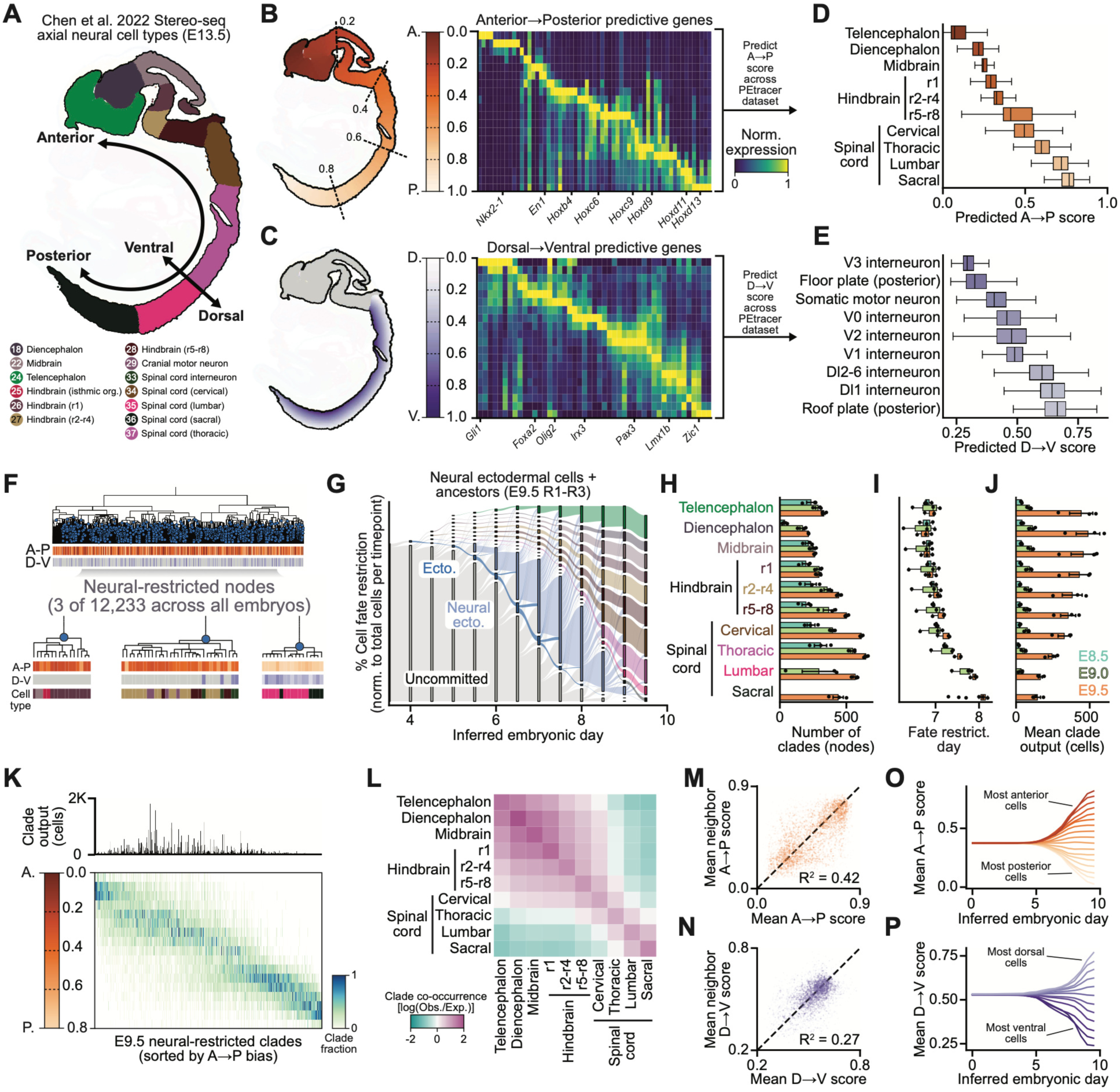
Spatial patterning and bias in the neural ectoderm. (**A**) E13.5 mouse embryo sagittal section (Stereo-seq^114^) with neural ectoderm colored by axial region and other tissues shown in the background. Cell type colors and numbers match Figure 2. A-P and D-V axes are indicated. Org. = organizer. (**B,C**) Left: Neural ectoderm colored by (**B**) A-P position and (**C**) D-V position. Right: Expression of the 61 A-P and 54 D-V patterning genes used to predict position. (**D**) Predicted A-P scores by axial region. (**E**) Predicted D-V scores by spinal cord cell subtype. For (D,E), boxes indicate mean and interquartile ranges. (**F**) Example E9.5-R1-C1 tree with neural-restricted nodes (blue circles) marked. Inset shows 3 of 12,233 neural-restricted nodes with descendant A-P score, D-V score, and identities provided. (**G**) Cell fate restriction over time at the level of ectoderm, neural ectoderm, and neural region for all neural ectoderm cells and their ancestors across E9.5 reps. 1-3. (**H**) Number of neural-restricted clades by region and timepoint, (**I**) mean fate restriction day, and (**J**) mean number of descendants. Bars in (H-J) show mean ± SEM across three replicate embryos. (**K**) A-P distributions for E9.5 neural-restricted clades with >10 descendants, ordered by mean A-P score; clade size shown above. (**L**) Neural region clade co-occurrence [log(observed/expected)] within E9.5 neural-restricted clades. (**M**) Mean A-P score versus nearest neural-restricted lineage neighbor; dashed line, y = x. (**N**) As in (M), for spinal cord D-V score. (**O**) Mean A-P score of ancestors over inferred time for neural cells grouped and colored by their observed A-P position at E9.5. (**P**) Same as (O) for spinal cord cell D-V score.

To understand how neural lineage specification varies along the A-P axis, we identified 12,233 neural-restricted clades across all embryos (**Figure 6F**). These neural clades were often specified directly from uncommitted progenitors, remaining multipotent for about a day before regional fate restriction (**Figure 6G**). For each neural region, the number of contributing clades, their mean restriction time, and their size varied systematically along the A-P axis (**Figures 6H-J**). Anterior brain regions were seeded at ∼E7.0 by ∼250 early-restricted clades whose output expanded substantially over time, whereas posterior spinal cord populations continued to acquire new fate-restricted clades through E9.5^15,115^. These patterns distinguish two modes of neural tube formation: primary anterior neurulation and secondary posterior neurulation, the latter sustained by ongoing specification from axial progenitors^58,116^.

Anterior and posterior neural clades differed in inferred spatial distribution. In E9.5 trees, anterior clades tended to span broader A-P domains, consistent with their earlier specification, and frequently spanned molecularly distinct regions, including the forebrain-midbrain and midbrain-hindbrain boundaries (**Figures 6K**, **6L, S13A, and S13B**). Within the spinal cord, clades were often biased toward specific D-V positions, with minimal association between specification timing and D-V location (**Figures S13C and S13D**).

Finally, we assessed the heritability of spatial identity by comparing the mean inferred spatial position of each E9.5 clade to that of its nearest lineage neighbor (**Figures 6M and 6N**). Inferred A-P position showed substantial heritability (R² = 0.42), while inferred D-V position was less strongly correlated (R²= 0.27). Calculating spatial scores for ancestral nodes as the mean of their descendants revealed that subtle A-P bias emerged ∼E5.0, with ancestors of the most anterior and posterior cells already slightly skewed toward their eventual positions (**Figure 6O**). D-V bias arose at a similar time; however, unlike the symmetric A-P pattern, ventral bias was initially more pronounced than dorsal, consistent with neurulation, where ventral positions are specified earlier at the base of the neural groove (**Figure 6P**)^117^. Together, these results suggest that spatial organization is established prior to neural fate specification, particularly along the A-P axis.

### Timing of neural ectoderm regionalization

Because anterior clades cross regional boundaries, anterior regional identity is not determined at the time of neural-fate restriction, raising the question of how and when these boundaries are subsequently established. Regionalization could proceed hierarchically, with an early midbrain-hindbrain split preceding subdivision of the hindbrain into rhombomeres^118,119^. Alternatively, regional identities could be specified concurrently, according to each cell’s position along the A-P axis^120^. To distinguish these possibilities, we focused on the first rhombomere (r1), since it can be clearly distinguished from the adjacent *Otx2*⁺ midbrain and the more posterior *Hoxa2*⁺ rhombomeres (r2-r4), with the *En1*⁺ isthmic organizer at the anterior edge of r1 (**Figures 7A and 7B**)^121–123^.

**Figure 7.**
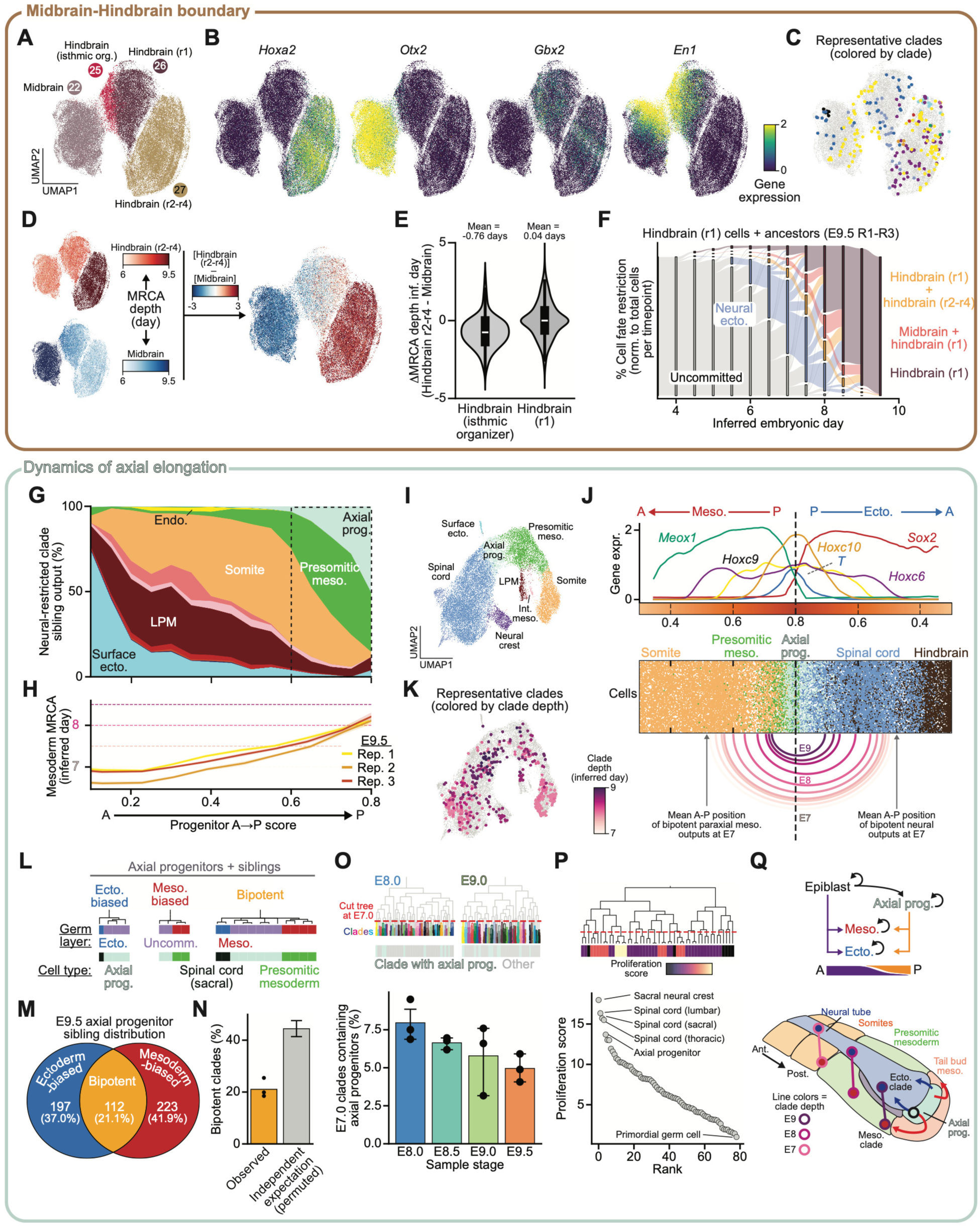
Midbrain-hindbrain boundary and the dynamics of axial elongation. (**A-F**) <u>Midbrain-hindbrain boundary:</u> UMAP of E9.5 midbrain, hindbrain isthmic organizer, r1, r2-r4 cells colored by (**A**) cell type, expression of (**B**) *Hoxa2*, *Otx2*, *Gbx2*, *En1*, and (**C**) 10 randomly sampled clades. (**D**) Inferred MRCA time between each cell and r2–r4 (top) or midbrain (bottom), and their difference (Δ), indicating relative lineage relatedness. (**E**) ΔMRCA depths (r2-r4 minus midbrain) for isthmic organizer and r1 cells. (**F**) Cell fate restriction over time at the level of neural ectoderm, r1 + r2-r4 or r1 + midbrain, and r1 only for all r1 cells and their ancestors across E9.5 reps. 1-3. (**G-Q**) <u>Axial elongation:</u> (**G**) Sibling clade outputs for E9.5 neural-restricted clades ordered by mean A-P score colored by developmental domain; paraxial mesoderm is split into somitic and presomitic groups. Box indicates region expanded in (I,J). (**H**) Mesodermal MRCA for neural ectoderm cells by inferred A-P position. Lines depict E9.5 mean ±95% CI. (**I**) UMAP of posterior neural-restricted clades and their siblings colored by developmental domain. (**J**) <u>Top:</u> Gene expression profiles of regional markers along the A-P axis showing *Meox1⁺* somites, presomitic mesoderm, *T⁺* axial progenitors, *Sox2⁺* spinal cord, and hindbrain cells. <u>Middle:</u> Cells ordered by A-P position, colored by developmental domain. <u>Bottom:</u> Bipotent clades grouped and colored by depth (E7-E9) with arcs connecting mean A-P position of mesodermal and ectodermal outputs. Dashed line indicates the posterior end of the embryo. (**K**) UMAP from (I) with representative neural-restricted clades and siblings colored by clade depth. (**L**) Three representative axial progenitor-containing clades and siblings with descendant germ layers and identities indicated. (**M**) E9.5 axial progenitor sibling outputs classified as ectoderm-biased, mesoderm-biased, or bipotent. (**N**) Percent bipotent clades compared to expectation given statistically independent specification (clade-size-stratified permutation). Error bar indicates 95% CI. (**O**) Persistence of axial progenitor-containing clades. E8.0–E9.5 trees were cut at inferred E7.0 and scored for whether they contained an axial progenitor. Bars show mean ± SEM of three replicate embryos. (**P**) Proliferation score, defined as the mean number of cells sharing a common ancestor within the previous day, ranked across E9.5 cell types. (**Q**) Model of fate specification during axial elongation and tailbud dynamics: direct epiblast specification dominates anteriorly and axial progenitors posteriorly; bipotent clades generate spatially offset neural and mesodermal outputs.

Since many neural-restricted clades span r1, we asked when r1 cells become lineage-restricted relative to adjacent regions (**Figure 7C**). We inferred the timing of each r1 cell’s most recent common ancestor (MRCA) with either midbrain or r2-r4 cells, using the difference between these times as a measure of relative lineage separation (**Figure 7D**). Cells within the isthmic organizer were more closely related to midbrain, whereas cells across the remainder of r1 showed comparable lineage relationships to both midbrain and r2-r4, with a gradient corresponding to A-P position (**Figure 7E**). This linkage reflected transient progenitor clades restricted to r1 plus midbrain, which were just as common as clades restricted to r1 plus r2-r4 (**Figure 7F**). Thus, r1 cells share recent ancestry with both midbrain and hindbrain rather than resolving through an early midbrain-hindbrain split, arguing that regional identities are specified concurrently from a shared progenitor field according to A-P position.

### Dynamics of posterior axis elongation

To understand how neural and mesodermal fates are jointly specified along the A-P axis, we examined the sibling clades of neural-restricted nodes at E9.5. Anterior siblings contributed to all three germ layers and were enriched for surface ectoderm, consistent with an epiblast origin, whereas siblings became increasingly enriched for paraxial mesoderm and axial progenitors toward the posterior end (**Figure 7G**)^15,58,116^. This trend was accompanied by progressively more recent neural-mesodermal separation, with 58% of the most posterior neural cells sharing a common ancestor with the mesoderm within the last 1.5 days (**Figures 7H and S13E**). Among these posterior bipotent clades, 42% contained an axial progenitor at E9.5 (**Figure S13F**). Transcriptionally, these cells formed a continuum from *Sox2*⁺ neural ectoderm to *T*⁺ axial progenitors to *Meox1*⁺ paraxial mesoderm, with bipotent clades ordered along the A-P axis by MRCA depth (**Figures 7I-K and S13G**)^124,125^. Notably, the inferred mean A-P position of bipotent clades at a given depth matched that of axial progenitors at the corresponding sampled stage, with mesodermal outputs adopting slightly more posterior Hox identities than their ectodermal siblings (**Figures 7J, S13H, and S13I**).

We then asked whether the bipotency of the axial progenitor pool reflects uniformly bipotent progenitors or a mixture of fate-biased progenitors. Classifying axial progenitor clades by sibling germ layer, we found 37.0% ectoderm-biased, 41.9% mesoderm-biased, and 21.1% bipotent – less frequent than expected under statistically independent specification (**Figures 7L–N**). Thus, although the axial progenitor pool collectively produces both fates, not all progenitors behave as bona fide bipotent neuromesodermal progenitors. To assess whether the axial progenitor pool is continuously replenished from many early founders or declines over time, we cut each tree at E7.0 and quantified the fraction of clades containing an extant axial progenitor. This fraction declined modestly across stages (∼8% at E8.0 vs. ∼5% at E9.5; **Figure 7O**), indicating that the axial progenitor pool remains polyclonal during elongation. Estimating proliferation rates from the number of cells sharing a common ancestor within the previous 24 hours we found that axial progenitors and posterior neural populations exhibited the highest proliferation scores in the embryo (**Figures 7P and S13J**).

Together, these findings support a model in which highly proliferative, polyclonal axial progenitors are maintained in the growing tailbud and continuously generate both neural and mesodermal lineages during elongation (**Figure 7Q**)^58,59,126^. As this pool progressively acquires increasingly posterior Hox identities, neural descendants tend to exit earlier and differentiate in the neural tube, whereas mesodermal descendants remain longer in the posterior growth zone before contributing to presomitic mesoderm. This temporal asymmetry may explain the consistent offset in Hox identity and A-P position between neural and mesodermal outputs observed in fate-mapping studies^15,127–130^.

## DISCUSSION

Here we map cell fate specification from pre-gastrulation through early organogenesis in the mouse by integrating cell state and lineage measurements at embryo scale. Using continuous cell marking with PEtracer, we reconstruct lineage trees for 16 embryos comprising >1.4 million cells sampled at half-day intervals from E7.5 to E10.0, resolving >2.1 million cell edges (∼75% of the total). Near-complete cellular sampling captures how cell fate decisions unfold throughout development, while deep transcriptional profiling links lineage architecture to cellular identity. This dataset can be explored at mela.wi.mit.edu.

Classic fate-mapping approaches have provided mechanistic insights into specific lineages and remain foundational for defining progenitor potential and tissue relationships^16,131,132^. More recent lineage tracing studies have resolved lineage relationships across multiple tissues, but have been limited to describing high-level relationships because of their sparse labeling^2,28,33,133^. Building on this body of work, our densely sampled lineage trees resolve the majority of cell divisions across embryogenesis, enabling direct quantification of progenitor-progeny relationships across the embryo without prior knowledge of lineage-defining markers or when individual specification events are expected to occur. We measure progenitor dynamics, including when fates become restricted and how many clades contribute to each cell type, all with empirical estimates of reproducibility. Below we revisit fundamental principles of mammalian development for which our data offer new quantitative insight.

First, developmental lineage architectures are highly reproducible across timepoints and embryos. Fate restriction trajectories, cell type ancestral linkages, progenitor clade numbers, clonal outputs, and commitment timing showed minimal variation across replicates, with pairwise linkage estimates differing by <6 hours. This reproducibility reflects the highly coordinated, regulative nature of mammalian development^1,134^, in which robust population-level outcomes emerge from individual cells acquiring fates in response to their position and local signaling environment.

Second, germ layer specification begins before gastrulation and extends beyond it. Subtle biases in epiblast fate distributions can first be detected at ∼E4.0 based on branch length estimates–around the time of implantation but well before the onset of gastrulation–suggesting that spatial organization and signaling in the post-implantation epiblast begin to bias progenitor outputs^10,11,135–138^. Specification persists at the posterior end of the embryo; our data show that bipotent axial progenitors continue generating additional mesodermal and ectodermal progenitors through E9.5, ultimately producing more germ-layer-restricted clades than are specified directly from the epiblast^15,58,59,126,139,140^.

Third, cell types can converge on shared transcriptional programs from distinct lineage origins. Convergence has long been described qualitatively; our approach makes it quantitative, resolving the full set of contributing origins along with their relative contributions and timing^2^. Endothelial cells arise from multiple mesodermal sources–including lateral plate, intermediate, and paraxial mesoderm–that each activate core endothelial genes while retaining subtle signatures of their developmental origins^100,103–105,141^. This convergence can be obscured by approaches that infer lineage from transcriptional similarity alone (*e.g.,* optimal transport), underscoring the importance of direct quantification of cell lineage dynamics^22,26^.

Finally, spatial position can outweigh lineage identity in determining cell fate. At the midbrain-hindbrain boundary, clades frequently span both regions despite sharp transcriptional differences, indicating that regional boundaries arise from downstream molecular specification rather than early lineage compartmentalization^118,120,122^. Similarly, trunk neural crest progenitors retain the capacity to generate both DRG and autonomic neurons at the time of fate restriction, with these identities resolving over time in response to local signaling cues likely encountered during migration^62–64,95^. The transient nature of these multipotent progenitors likely explains the discrepancy with recent static barcoding results^97^, highlighting the value of continuous lineage recording for capturing short-lived intermediates.

This work, and other concurrent efforts^53^, provide a foundation for future studies integrating spatial information, additional timepoints, and perturbations. Our A-P and D-V scores offer a preview of what imaging-based readout of PEtracer will reveal: with precise spatial positions, lineage relationships could be mapped onto tissue architecture, resolving how local signaling, morphogen gradients, and cell-cell interactions shape fate *in vivo*^43,47^. Extending this framework to later stages and combining it with genetic and environmental perturbations will define how cell fate programs are established across development and reactivated or dysregulated in adult homeostasis, regeneration, and disease^26,131,142–145^. Integrating our paired lineage and transcriptional measurements across time could further enable computational models that learn the regulatory logic and genetic programs driving fate specification. This work represents an important step toward a lineage-resolved, embryo-scale reference for cell fate specification across mouse development.

### Limitations of the study

Reconstructed lineage trees are not a perfect record of developmental history. Unmarked divisions and incomplete cellular sampling can obscure some progenitor-progeny relationships, and branch lengths rely on the assumption that edit accumulation is approximately proportional to elapsed time, so time estimates should be interpreted accordingly. Cell type annotations are limited by the resolution of current references and the possibility of residual doublets in our dataset. Spatial coordinates were imputed rather than measured directly. Further, these trees capture cell fates rather than their developmental potential–cells observed to adopt a given fate here may retain the capacity to generate others if placed in an ectopic environment^146^. Finally, the embryos profiled here are chimeras seeded by 2-6 donor mESCs rather than derived from a single zygote, which may introduce slight differences in developmental timing and progenitor allocation. While host-derived wild-type cells provide an internal control for developmental competence, with cell type distributions closely matching established references, the earliest fate decisions should be interpreted in this engineered context. Additionally, while we observe no overt effects of lineage tracing on development, subtle effects of tracing activity cannot be fully excluded.

## RESOURCE AVAILABILITY

### Lead contact

Further information and requests for resources and reagents should be directed to and will be fulfilled by the lead contact, Jonathan S. Weissman.

### Materials availability

- Plasmids generated in this study have been deposited to Addgene: Pooled library 257853 = Non-silencing mESC piggybac LTC library; Plasmid 257851 = Rosa26 PEmax-T2A-GFP knock-in cassette; Plasmid 257852 = Upstream U6 24-mer acceptor backbone
- Engineered mouse embryonic stem cell lines carrying the PEtracer system generated in this study are available from the lead contact with a completed materials transfer agreement.

### Data and code availability

- Single-cell RNA-seq data have been deposited at GEO as <u>GSE342607</u> and are publicly available as of the date of publication.
- Processed data have been deposited at Zenodo and are publicly available as of the date of publication at https://doi.org/10.5281/zenodo.19892784.
- All original code has been deposited on Zenodo at https://doi.org/10.5281/zenodo.22833553 and is also publicly available at of the date of publication. The version-controlled repository is additionally maintained at https://github.com/jweissmanlab/MELA-2026
- Any additional information required to reanalyze the data reported in this paper is available from the lead contact upon request.

## ACKNOWLEDGMENTS

We thank members of the Weissman, Smith, Loh, and Yosef labs for helpful discussions and intellectual/technical support, including N. Dias, M. Carlino, A. Masaltseva, R. Jokhai, R. Saunders, and P. Zheng. We also thank the Whitehead Institute Mouse Genetically Engineered Models (GEM) Center including Randy Curry for colony management support, the Whitehead Flow Cytometry Core, the Whitehead Genome Technology Core, the Broad Institute Clinical Labs, the Broad Institute Klarman Cell Observatory, and the Yale Animal Research Center (YARC). We also thank J. Feidler, A. Landsberger, M. Peetz, and K. Zill of the MPI Transgene unit for their support and expertise, as well as the MPIMG Microscopy, Sequencing, and Flow Cytometry Cores.

## AUTHOR CONTRIBUTIONS

Conceptualization: LWK, WNC, JSW

Methodology: LWK, WNC

Investigation: LWK, WNC, JV, TCJH, MW, GG, WC, LAS, DC, KS, AY, LW

Software: WNC, GG

Formal analysis: WNC, LWK, GG

Visualization: WNC, LWK, GG

Funding acquisition: ZDS, JSW

Resources: LWK, JV, TCJH, MW, WC, SM, KEY

Supervision: ZDS, JSW, NY, XZ

Writing –original draft: LWK, WNC, JSW

Writing –review & editing: LWK, WNC, ZDS, JSW, KML, JV, MW, TCJH, GG, KEY, WC, LAS, DC, KS, AY, SM, NY

## DECLARATION OF GENERATIVE AI AND AI-ASSISTED TECHNOLOGIES IN THE WRITING PROCESS

During the preparation of this work the author(s) used ChatGPT and Claude to generate code, perform automated cell type and gene program annotation, and for manuscript proofreading. After using this tool/service, the author(s) reviewed and edited the content as needed and take(s) full responsibility for the content of the published article.

## SUPPLEMENTAL INFORMATION

Document S1. Supplementary Figures S1-S13.

Table S1. Embryo-level sequencing and tracing statistics, related to Figure 2.

Table S2. List of cell subtypes with markers and rationale, related to Figure 2.

Table S3. E9.5 pairwise linkage values with annotation, related to Figure 3.

Table S4. E9.5 fate-restricted clade statistics, related to Figure 3.

Table S5. Gene programs with genes and descriptions, related to Figure 5.

## METHODS

## KEY RESOURCES TABLE

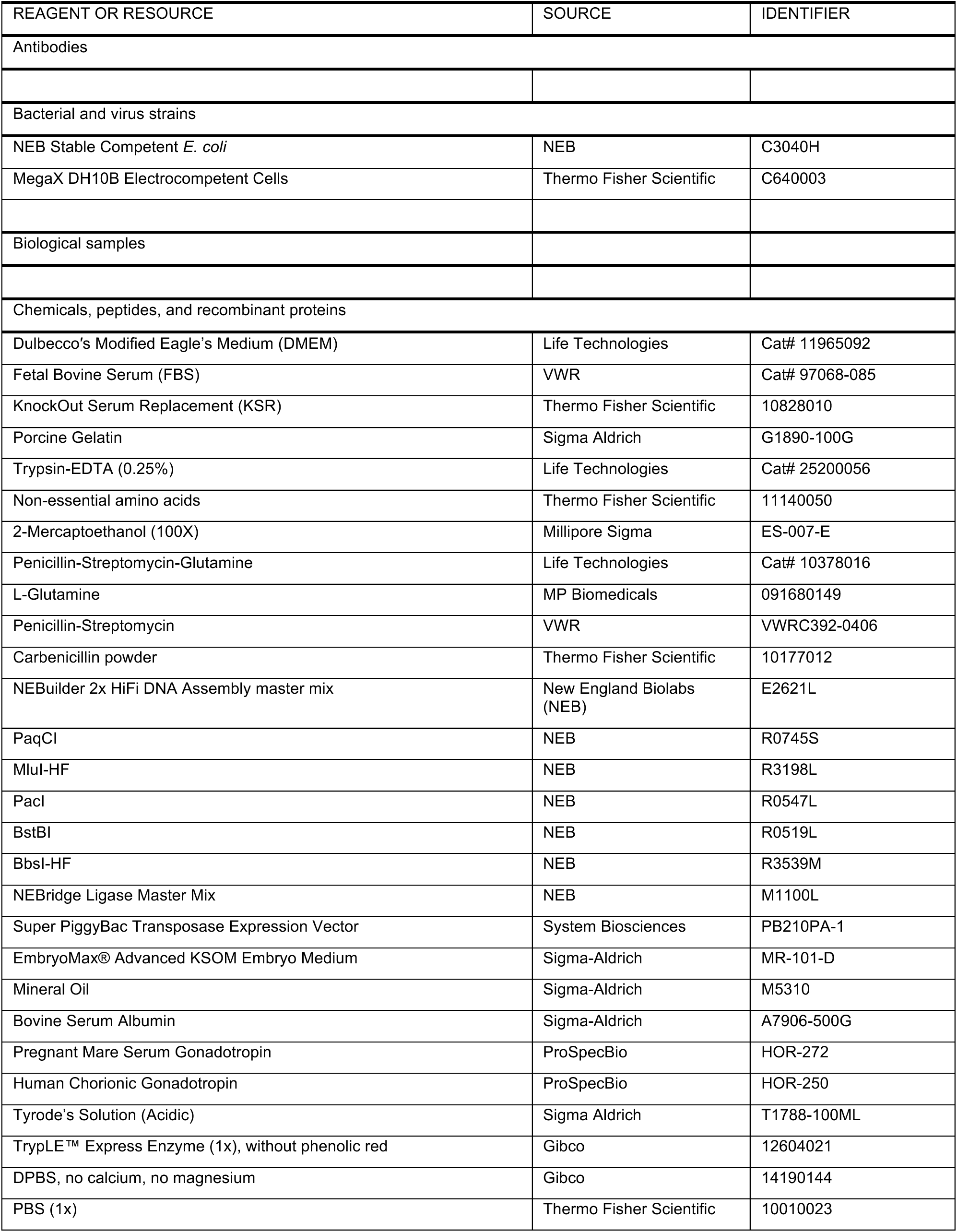
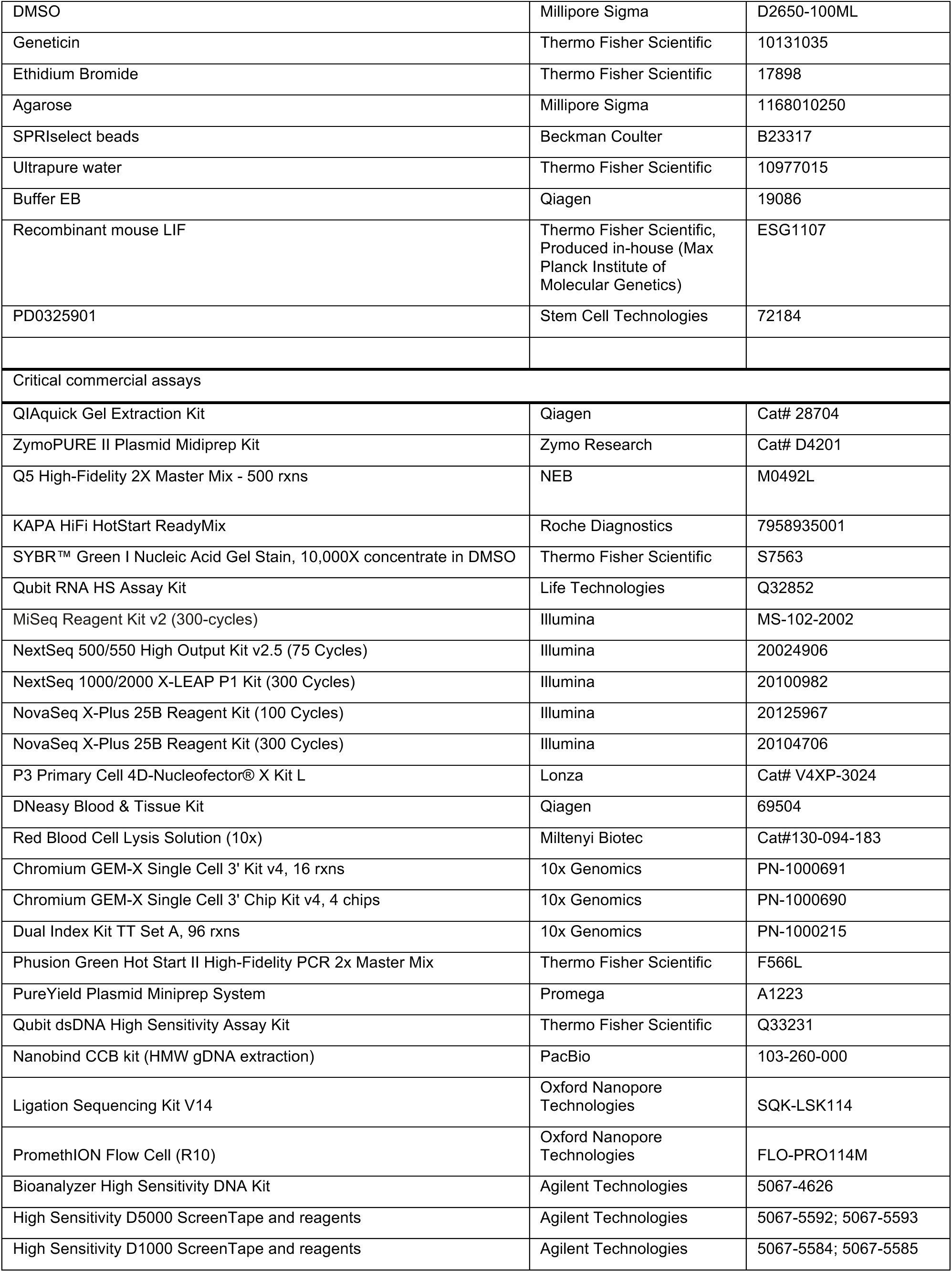
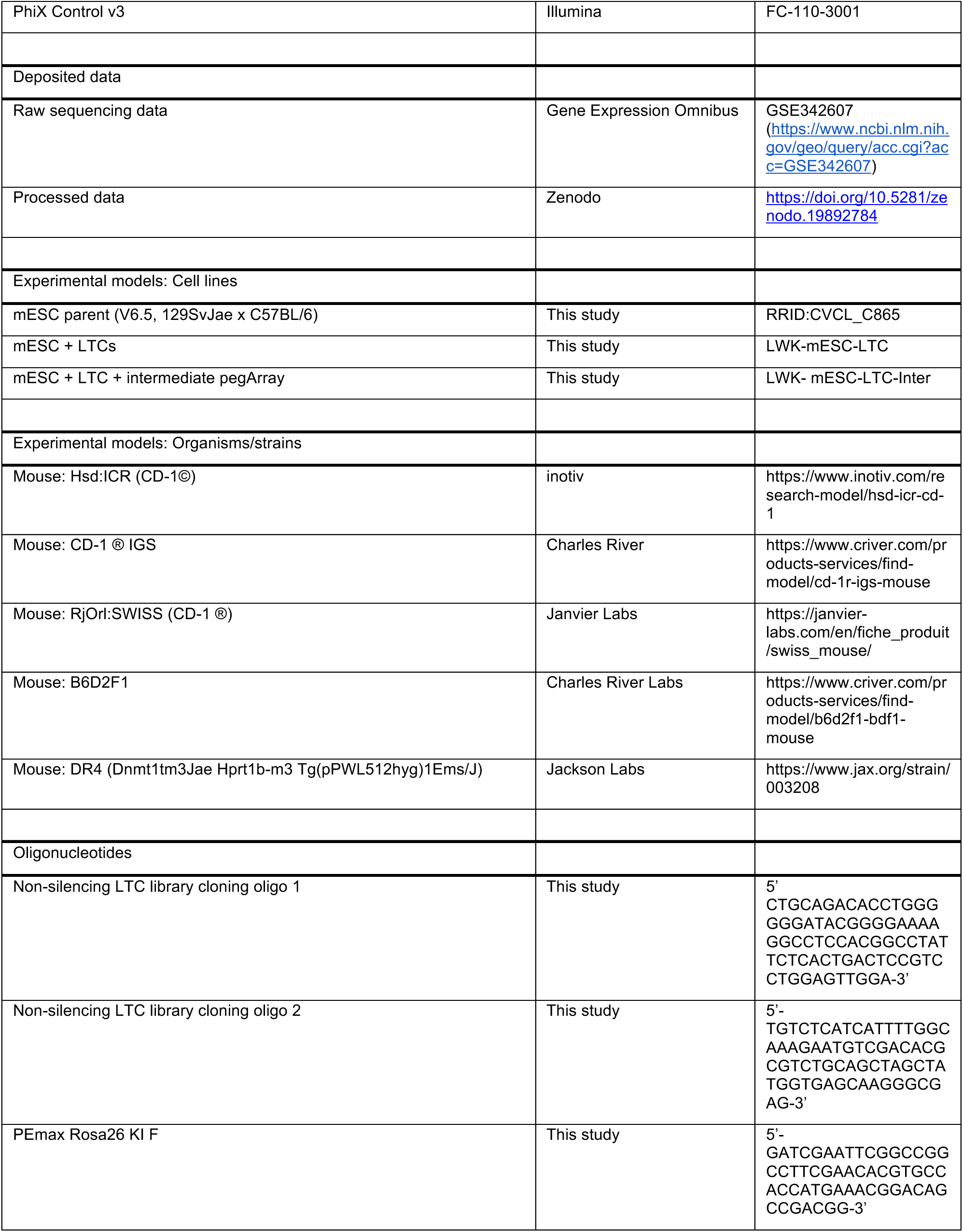
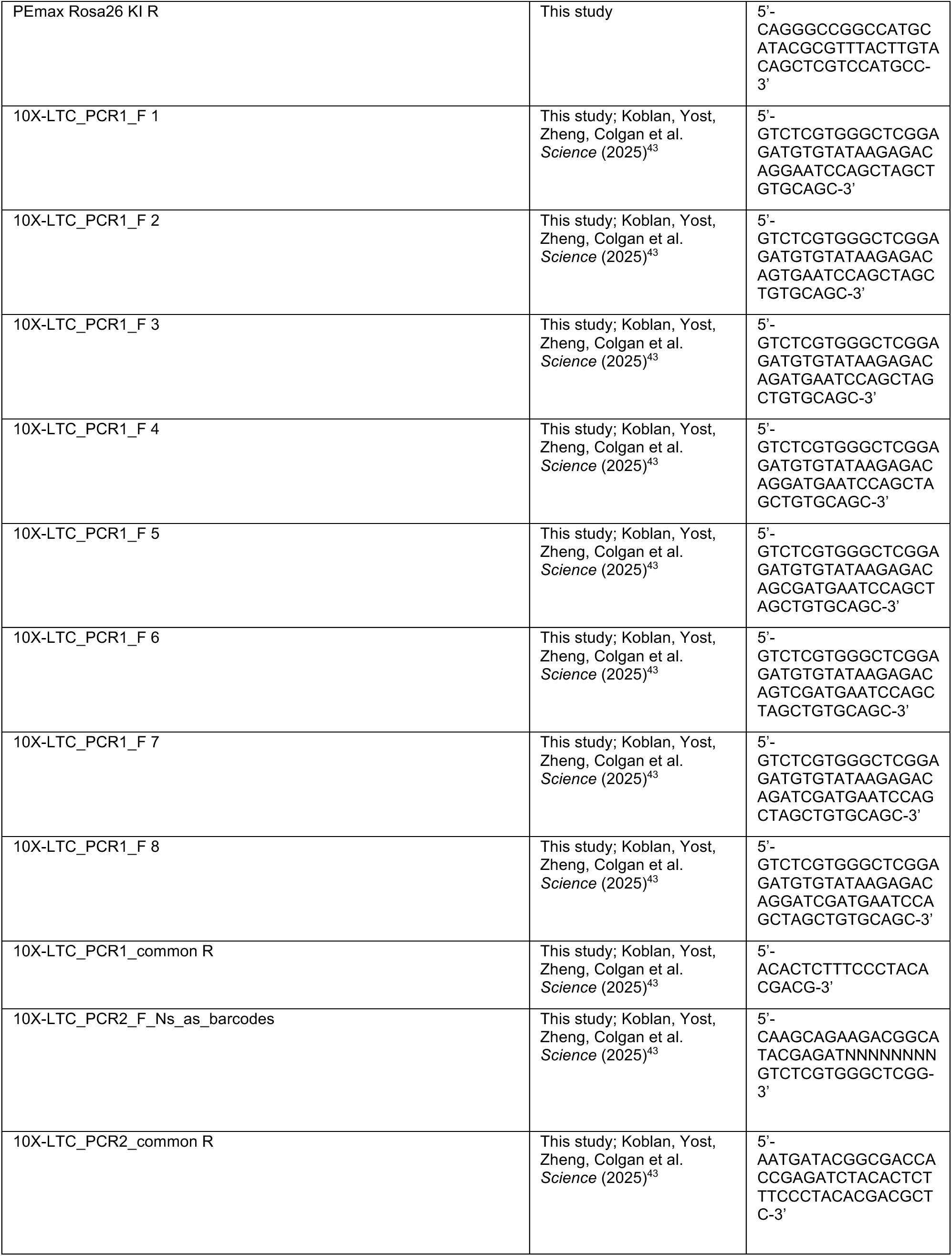
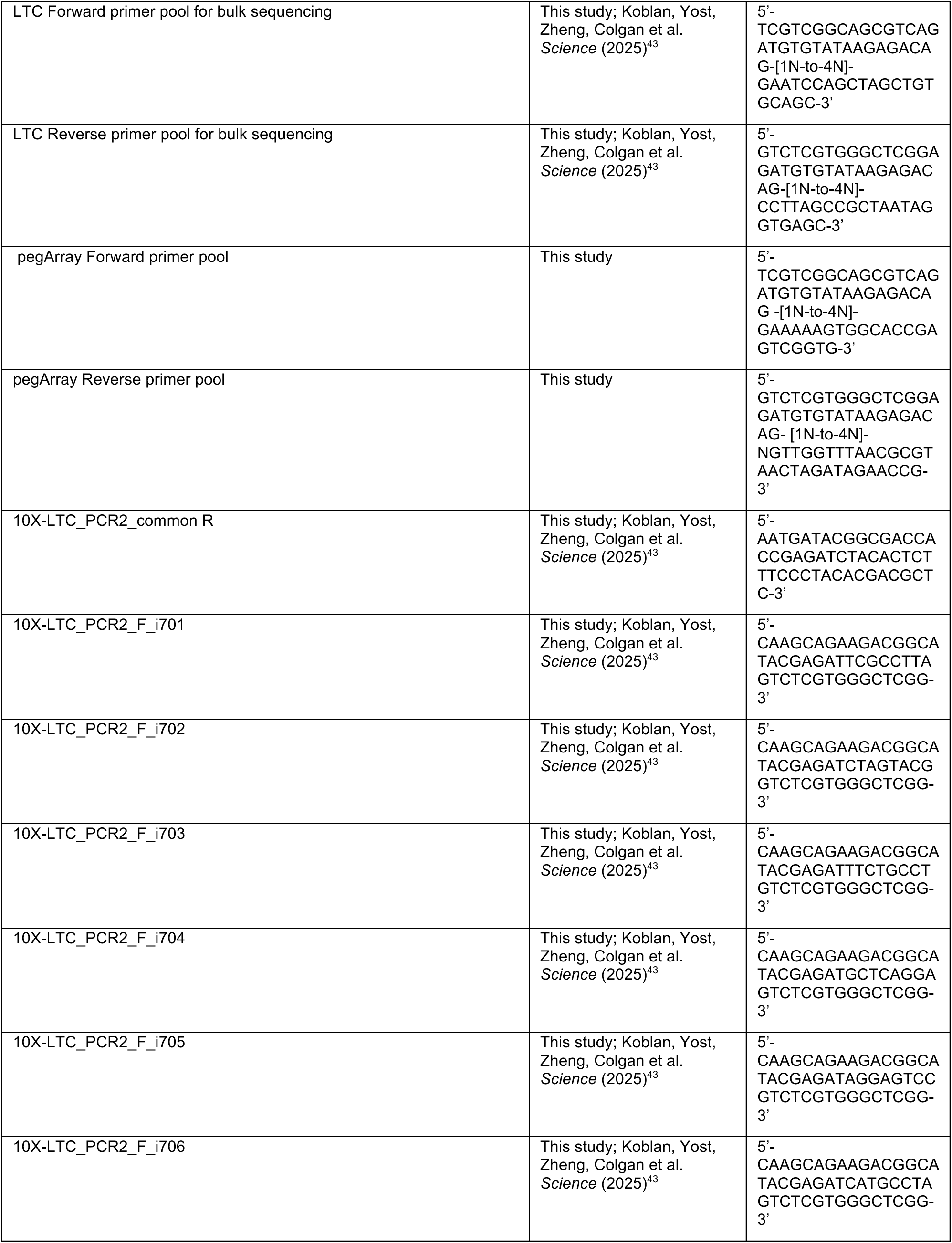
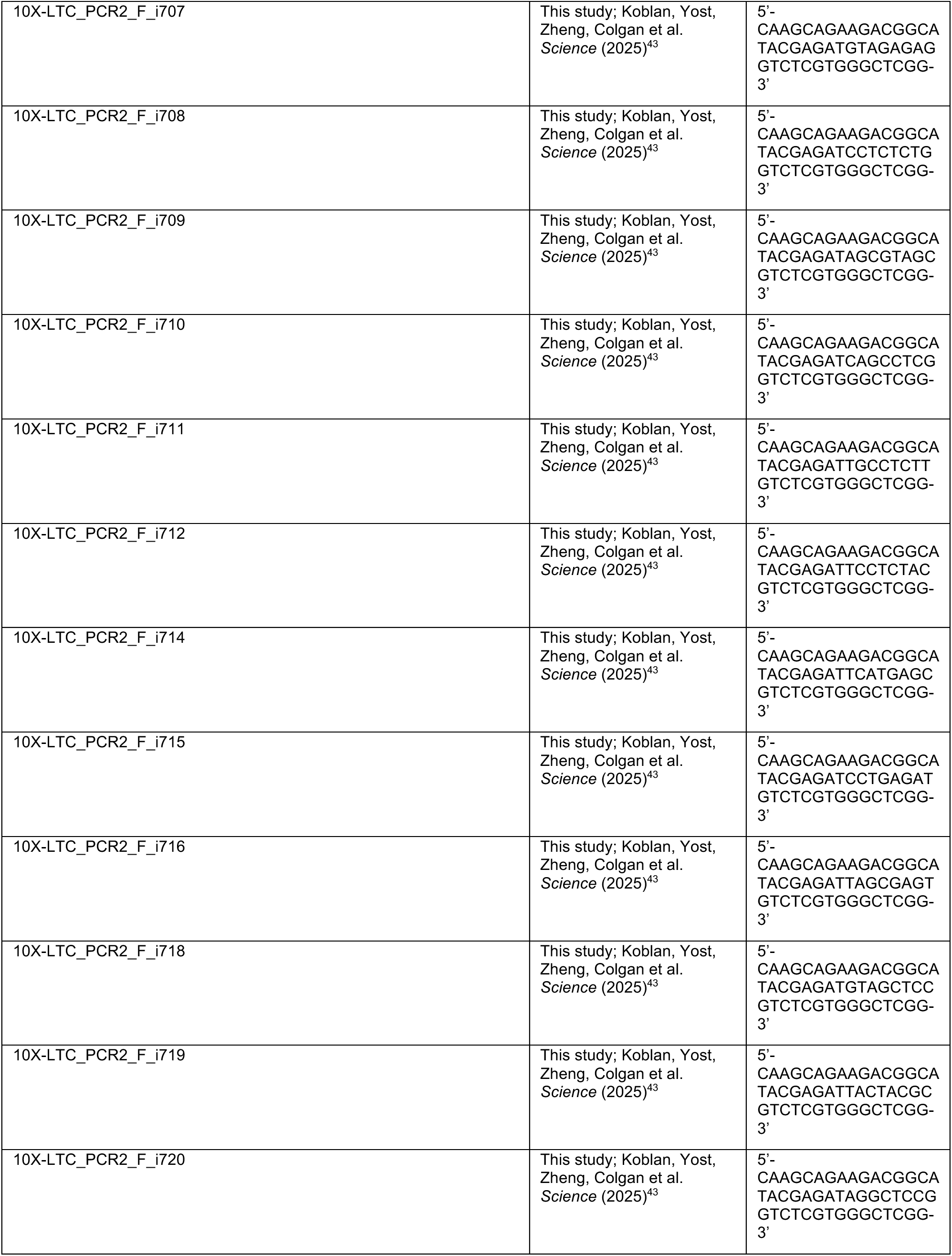
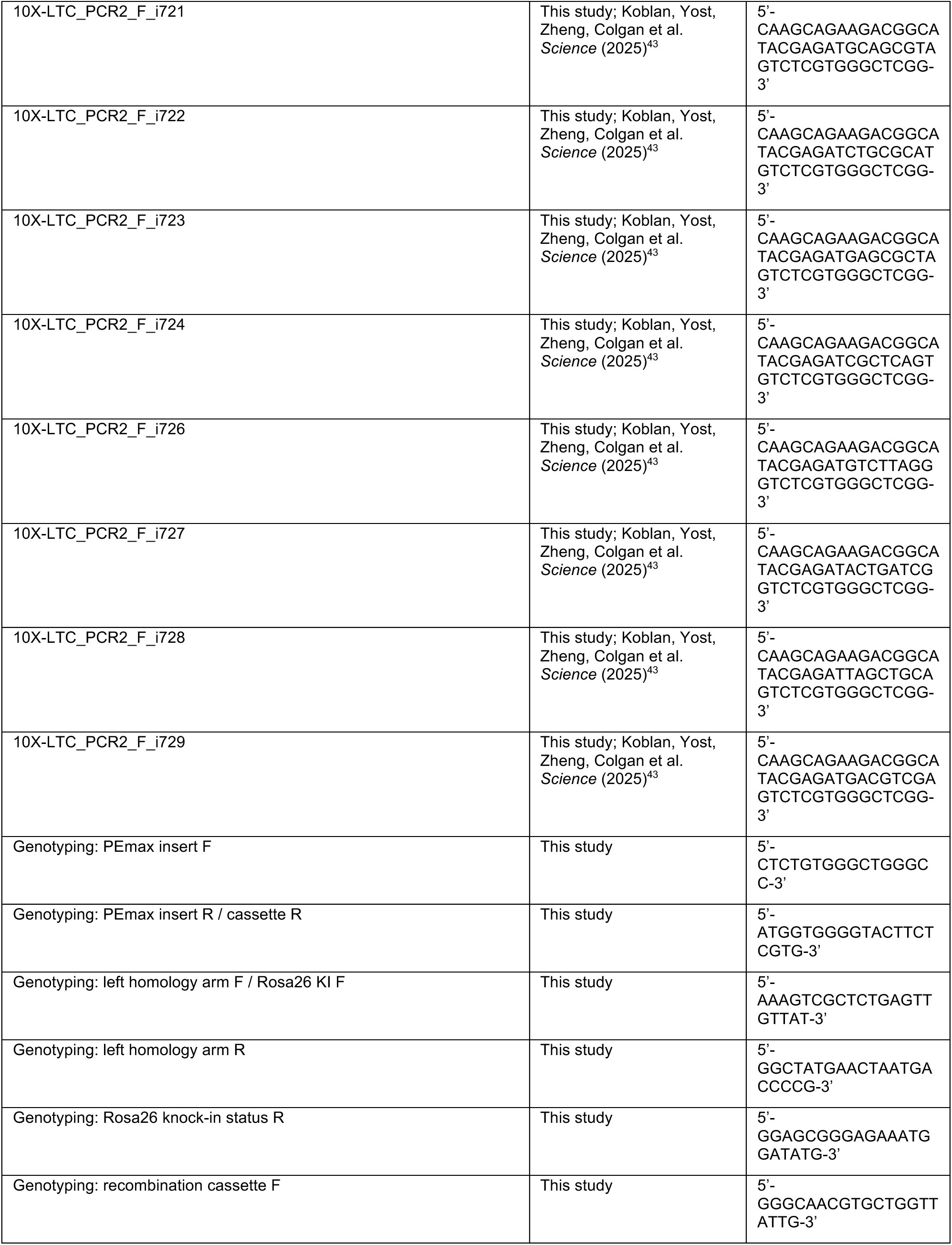
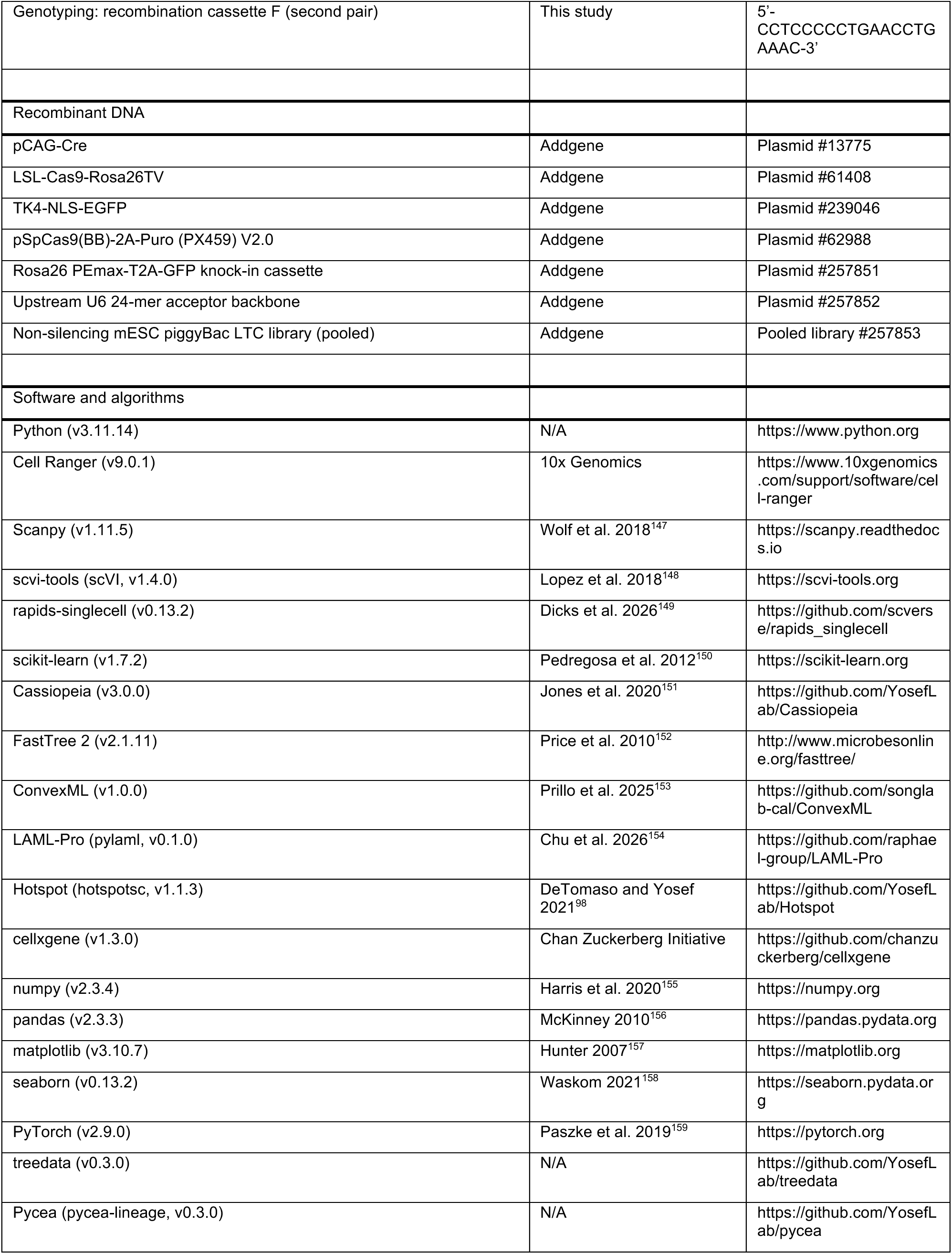
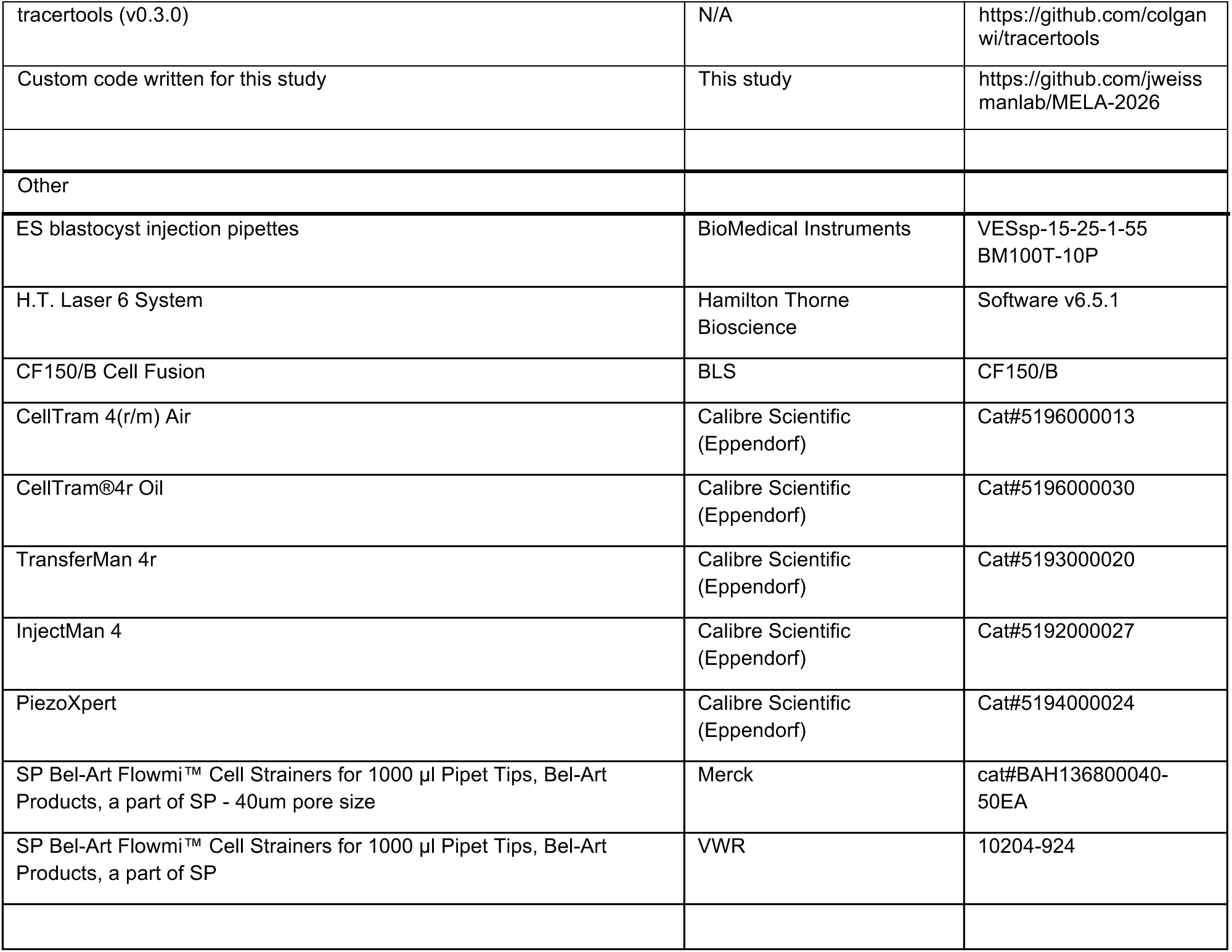

## EXPERIMENTAL MODEL AND STUDY PARTICIPANT DETAILS

### Mice

All procedures were performed in specialized specific-pathogen-free facilities, following all relevant animal welfare guidelines and regulations, with mice kept on a 12 h light-dark cycle from 6 am to 6 pm, and provided a standard diet ad libitum. CD-1 females were 8–32 weeks of age when used to generate donor embryos and 8–15 weeks (28–40 g) when used as pseudopregnant recipients for embryo transfer; male CD-1 mice used for matings were between 8 weeks and 1.5 years of age. All donor cells in the study with ongoing lineage tracing are male. Chimeric embryos were collected between E7.5 and E10.0 for single-cell analysis, and at E12.5, E15.5, and E18.5/term to assess development into later gestation. All mouse experiments described in this study were approved by the Massachusetts Institute of Technology Institutional Animal Care and Use Committee (IACUC, protocol numbers 2311000598 and 2208000407), the Yale University Committee on the Use of Animals in Research and Teaching (IACUC, Protocol #2026-20357) or by the LaGeSo Berlin (Protocol #G0098/23).

### METHOD DETAILS

#### Cloning of the Rosa26 PEmax-T2A-GFP knock-in cassette

Gibson assembly was used to assemble DNA fragments encoding PEmax-T2A-GFP into a digested backbone designed for knock-in of genetic cargos into the mouse genomic Rosa26 locus (Addgene plasmid #61408)^160^. Briefly, backbone plasmid was digested with MluI (NEB) and BstBI (NEB) according to manufacturer’s recommendations. PEmax-T2A-GFP was amplified using Q5 High-Fidelity 2X Master Mix (NEB). Fragments were assembled into complete plasmids using 5 μL of NEBuilder 2X HiFi DNA Assembly master mix (NEB) combined with 0.05pmol backbone and 0.15pmol of each insert fragment brought to a final volume of 10 μL with ultrapure nuclease-free water. Assemblies were incubated for 60 minutes at 50°C prior to transformation into NEB stable competent *E. coli* (C3040, NEB). Transformation included a 15-minute incubation of 50 μL of competent cells and 5 μL of assembly mix prior to a 30-second heat shock at 42°C and a subsequent 5-minute incubation on ice before plating on 100 μg/mL carbenicillin-containing agar plates.

#### Cloning of upstream U6-containing 24-mer pegArrays

24-mer pegArrays were cloned as previously described^43^, substituting a novel backbone that includes two U6 promoters that do not express any gRNA upstream of the 24-mer array. This strategy was used to minimize positional effects that decrease the expression levels of the first two pegRNAs in our arrays. Briefly, 24-mer assemblies require four pieces of DNA that include a PaqCI digested 24-mer backbone and a PaqCI digested fragment for each edit site 8-mer prepared as described previously. PaqCI digestions were performed overnight using 4 μL of CutSmart buffer, 4 μL of PaqCI activator, 2 μL of PaqCI, and 30 μL of the plasmid of interest. These digested fragments were gel extracted using the QIAquick Gel Extraction Kit (Qiagen) following manufacturer’s recommendations. Digested fragments were assembled into 30 μL PaqCI golden gate assemblies with 3.33 nM of the PaqCI digested 8-mer fragments (typically 1 μL of the ∼4,000nt fragment diluted to 150 ng/μL), 1 μL of PaqCI pre-digested 24-mer acceptor backbone diluted to 25 ng/μL, 2.5 μL PaqCI enzyme, 1.25 μL PaqCI activator, 10 μL NEBridge Ligase Master Mix (M1100L), and water to bring the reaction to 30 μL total volume. Assemblies were carried out following manufacturer’s recommendations, carrying out 99 cycles of 5 minutes at 37°C and 5 minutes at 16°C before a 10-minute 60°C incubation and subsequent 4°C hold. These reactions were transformed into NEB Stable Competent *E. coli* (NEB, C3040), incubating on ice for 20 minutes prior to a 30 second heat shock at 42°C. After heat shock, bacteria were incubated on ice for 5 minutes prior to plating on 100 μg/mL carbenicillin-containing agar plates. Plates were incubated overnight at 30°C prior to picking colonies into 5 mL of 100 μg/mL carbenicillin-containing terrific broth, outgrowth overnight at 30°C, and DNA isolation using the Promega PureYield DNA Miniprep System. Some bacteria were expanded into 100 mL cultures overnight while isolated plasmids were diluted to 100 ng/μL and submitted for whole plasmid sequencing. DNA was isolated from these larger cultures using the ZymoPure II Plasmid Midiprep Kit. Plasmids were sequence confirmed by whole plasmid sequencing and further validated to have the correct insert by gel electrophoresis to be the correct size using restriction enzyme digestion.

#### Cloning of a library of non-silencing PEtracer lineage tracing cassettes

We cloned our previous library of PEtracer lineage tracing cassettes (LTCs) into a backbone compatible with PiggyBac transposition, which leveraged recent developments in non-silencing promoter elements derived from Addgene plasmid #239046^161^. The cloned parent backbone was digested with PacI and the LTC library was introduced via Gibson assembly as previously described^43^. Libraries were transformed into MegaX electrocompetent cells as previously described, and DNA was isolated using the ZymoPure II plasmid midiprep kit (Zymo Research). The resulting plasmid library was sequence confirmed by whole plasmid sequencing (Plasmidsaurus, Quintara) prior to introduction into mouse embryonic stem cells (mESCs).

#### Plasmid availability

The cloned plasmids and LTC library listed above are available through Addgene:

- Rosa26 PEmax-T2A-GFP knock-in cassette (Plasmid #257851)
- Upstream U6 24-mer acceptor backbone (Plasmid #257852)
- Non-silencing mESC piggybac LTC library (Pooled library #257853)

#### General Mouse Embryonic Stem Cell Culture Conditions

Mouse embryonic stem cells (mESCs) were maintained on mitotically inactivated DR4 mouse embryonic fibroblast (MEF) feeders in mESC medium, as previously described^162^. Briefly, cells were cultured in DMEM (Gibco) supplemented with 10% fetal bovine serum (HyClone), 1,000 units/mL leukemia inhibitory factor (ESGRO; Sigma), 2 mM L-glutamine (MP Biomedicals), nonessential amino acids (Gibco), penicillin-streptomycin (VWR), and β-mercaptoethanol. Cultures were maintained at 37°C in 5% CO_2_.

MEFs were mitotically inactivated by gamma irradiation and cryopreserved prior to use. For ESC culture, dishes were coated with 2% gelatin at room temperature for ≥30 min. MEFs were plated 24-48 hours before ESC seeding at the following densities (cells/well): 6-well: 5 x 10^3^; 12-well: 2.5 x 10^3^; 48-well: 6.25 x 10^3^; and 10-cm dish: 3 x 10^6.^

Dissociation of ESCs was carried out every 2–3 days by using 0.25% trypsin (LIFE technologies) following a wash with HEPES-buffered medium (in-house HEPES buffer was prepared containing 18.4 mM HEPES, 133.5 mM NaCl, 5.2 mM KCl, 5.3 mM dextrose, 0.42 mM KH_2_PO_4_, and 0.29 mM Na_2_HPO_4_·7H₂O, phenol red, pH 7.25-7.35, 285–295 mOsm/kg) and incubation at 37°C for 5-10 min. Trypsinization was quenched with serum-containing medium (2x or more the volume of trypsin), and cells were triturated to a single-cell suspension before being pelleted by centrifugation (200 RCF, 5 min).

For cryopreservation, an equal volume of 2x ESC freezing medium (50% FBS, 30% mESC medium, 20% DMSO) was added to the cell suspension. Cells were transferred to -80°C in a slow-cool chamber (Nalgene), for at least 24 hours before storage in liquid nitrogen.

#### Generation of PEmax-T2A-GFP mESCs

PEmax-T2A-GFP mESCs were generated by homology-directed repair using two plasmids: (i) a Rosa26-PEmax-T2A-GFP knock-in donor plasmid (described above) and (ii) a plasmid encoding S. pyogenes Cas9 and a Rosa26-targeting sgRNA to induce a double-strand break at the Rosa26 locus. The Cas9/sgRNA plasmid was derived from Addgene plasmid #62988, into which the Rosa26 sgRNA sequence (GACTCCAGTCTTTCTAGAAGA) was cloned using BbsI-HF restriction sites. Low passage (p17) V6.5 (129SvJae×C57BL/6)^163^ ESCs were thawed three days prior to nucleofection. On the day of the nucleofection, cells at 60–70% confluency were dissociated using trypsin (see above) and pre-plated on 2% gelatin-coated dishes for 30 min to deplete MEFs. Non-adherent ESCs were collected, counted, and 1 x 10^6^ cells were pelleted for nucleofection. For nucleofection, cell pellets were resuspended in 100 µL of P3 Nucleofector solution (Lonza, V4XP-3024) and mixed with 0.5 µg of the Cas9/sgRNA plasmid and 1.5 µg of the Rosa26-PEmax-T2A-GFP donor plasmid. Nucleofection was performed using a 4D-Nucleofector X Unit (Lonza) with program CG-104. Immediately after nucleofection, cells were plated at clonal density (1,000–5,000 cells per 10-cm dish) onto DR4 MEF feeders. Selection with G418 (300 µg/mL) was initiated 48 hours post-nucleofection and maintained for 6 days. Individual colonies (n = 48) were manually picked and expanded in 96-well plates.

#### Genotyping of PEmax-T2A-GFP mESCs

Colonies were isolated following HDR engineering and selection using neomycin at 300 µg/ml. Individual colonies were expanded and genomic DNA was extracted using QuickExtract DNA Extraction Solution (Biosearch Technologies, QE09050) following manufacturer’s protocols; specifically, a 15-minute incubation at 65°C followed by a 5-minute incubation at 95°C followed by a hold at 12°C. Genotyping was carried out using the Phusion Green Hot Start II High-Fidelity PCR 2x Master Mix with the following cycle conditions: 98°C for 2 minutes, [98°C for 10 seconds, 65°C for 20 seconds, 72°C for 60 seconds]x34, 72°C for 2 minutes. Multiple primer pairs were used to check the integrity of multiple portions of the knocked-in sequence.

1. To check the presence of the PEmax insert: 363 bp amplicon with CTCTGTGGGCTGGGCC + ATGGTGGGGTACTTCTCGTG
2. To check the integrity of the left homology arm sequence: 283 bp amplicon with AAAGTCGCTCTGAGTTGTTAT + GGCTATGAACTAATGACCCCG
3. To check the status of a Rosa26 genomic knock-in: 607 bp if there is no knock-in or a heterozygous knock-in; no amplification if homozygous knock-in with AAAGTCGCTCTGAGTTGTTAT + GGAGCGGGAGAAATGGATATG
4. To check the status of the recombination cassette: 1550 bp if LSL is intact, 882 bp if LSL is recombined with GGGCAACGTGCTGGTTATTG + ATGGTGGGGTACTTCTCGTG
5. An additional primer pair to check the status of the recombination cassette: 1271 bp if not recombined, no amplification if recombined with CCTCCCCCTGAACCTGAAAC + ATGGTGGGGTACTTCTCGTG

#### Creation of a homozygous PEmax-T2A-GFP mouse strain

mESC clones G3 and F3, confirmed to be heterozygous for PEmax expression, were used for blastocyst injection as previously described^17^. Briefly, ESC clones were thawed and cultured for 3 days prior to dissociation for microinjection. Donor embryos were obtained from B6D2F1[BDF1] females (C57BL/6J x DBA2, Charles River) following superovulation induced by pregnant mare serum (7.5 IU PMS, ProSpecBio), followed 48 hours later by human chorionic gonadotropin (7.5 IU hCG, ProSpecBio), and subsequent overnight mating with BDF1 males. Eight-cell stage embryos were collected at embryonic day 2.5 (E2.5) and cultured overnight in EmbryoMax®KSOM Advanced Embryo Medium (MilliporeSigma) under oil to the blastocyst stage. For injections, 4-8 ESCs were introduced into the blastocoel cavity of each embryo. A total of 20 injected blastocysts were transferred into the uterine horn of each pseudopregnant CD1 recipient female at day 2.5 post coitum.

#### Isolation of homozygous PEmax-T2A-GFP mESCs

Morula-stage embryos were isolated from PEmax-T2A-GFP homozygous mice that had been superovulated (5 IU PMS followed by 5 IU hCG 48h later) and paired with homozygous males. Embryos were harvested 2 days post coitum and cultured overnight to the blastocyst stage in EmbryoMax®KSOM Advanced Embryo Medium (MilliporeSigma) under oil. ESC derivation was performed as previously described^164^. Briefly, blastocysts were exposed to Acid Tyrode’s solution (MilliporeSigma) for 30-60 seconds to remove the zona pellucida, followed by two to three washes in serum-containing mESC medium, from now on referred to as mESC medium. Individual embryos were then transferred into wells of a 96-well plate pre-seeded with MEFs. Embryos were cultured undisturbed for 5 days in serum-free medium supplemented with 15% KnockOut Serum Replacement (KSR) and 5 µM of the Mek1 inhibitor PD98059 (Cell Signaling Technology) from now on referred to as mouse ESC derivation medium (mESCD medium). On day 5, outgrowths were identified and dissociated following a single wash with HEPES-buffered medium, incubation with trypsin (5 min), and quenching with mESC medium. The entire cell suspension was transferred to a 48-well plate containing derivation medium and cultured until ESC colonies emerged. Three to four days later, ESCs were passaged onto one or two wells of a 12-well plate, depending on cell density, and subsequently maintained in mESC medium. Established cell lines were cryopreserved for later use and injection into blastocysts to generate chimeras. Clone sex was determined by PCR, and male clones were selected for karyotyping.

#### Karyotyping analysis

Cells/clones were thawed into T25 flasks covered with DR4 MEFs and cultured for 4-5 days in mESC Medium in order to prepare for submission. Cell lines were submitted to Cell Line Genetics according to supplier’s protocols for karyotyping analyses.

#### Establishing PEmax-T2A-GFP mESCs with high numbers of integrated LTCs

PEmax-T2A-GFP mESCs were thawed 3 days prior to nucleofection. After dissociation, 2 x 10^6^ cells were resuspended in 100 µL P3 Nucleofector solution (Lonza, V4XP-3024) containing 5 µg of donor plasmid and 1 µg of Super PiggyBac transposase expression vector. Nucleofection was carried out using a 4D-Nucleofector X Unit (Lonza; program CG-104) and all cells were immediately plated in a single 6-well covered with MEFs. Cells were passaged 1:1 to a 10cm plate 2 days after. After nucleofection, cells were cultured for 7 days with passaging every 2-3 days to allow recovery and expansion. The top 5% of mCherry⁺ cells were isolated by fluorescence-activated cell sorting (FACS; BD FACSAria II, BD Biosciences). Sorted cells (2,000 per 10-cm dish; three dishes per clone) were plated onto MEF feeders and cultured for 5-7 days to allow colony formation. Individual colonies were manually picked into 96-well plates containing MEFs. The following day, cells were passaged onto fresh 96-well plates seeded with half-density MEFs and cultured for 2 days, followed by a 1:3 split and further expansion for 3 days. Three replicate plates were cryopreserved, two frozen for future expansion and genomic DNA was isolated from the third.

#### Screening for Sorted PEmax-T2A-GFP mESCs with high numbers of integrated LTCs

Genomic DNA was isolated from mESC clones using QuickExtract DNA isolation solution according to manufacturer’s protocols (Millipore Sigma). The number of integrated LTCs per clone was determined following amplification and bulk sequencing as previously described^43^. Briefly, two successive rounds of PCR were used to amplify lineage tracing cassettes with 2 μL of gDNA and 500 nM primer mix using the Phusion Green Hot Start II High-Fidelity PCR 2x Master Mix with the following cycle conditions: 98°C for 2 minutes, [98°C for 10 seconds, 65°C for 20 seconds, 72°C for 30 seconds]x28, 72°C for 2 minutes. PCR1 was conducted using a pool of forward and reverse primers where each primer was pooled in an equimolar fashion to enable sequencing on an Illumina sequencer with minimal need for added PhiX. All PCR1 amplicons were checked on 2% agarose (VWR) gels with 1 μL of ethidium bromide (Thermo Fisher Scientific) per 10 mL of 1x Tris-Acetate-EDTA (TAE) buffer. PCR2 amplification was carried out using Nextera adapter PCR primers (listed in Key Resources Table) with the following conditions: 98°C for 2 minutes, [98°C for 10 seconds, 61°C for 20 seconds, 72°C for 30 seconds]x8, 72°C for 2 minutes. 5 μL of PCR1 products were used for each PCR2 reaction and 500 nM final primer concentrations were used. Following PCR2, all reactions for a given genomic locus were pooled together, run on a 2% agarose gel, and gel extracted using the QIAquick gel extraction kit (Qiagen). Concentrations of purified libraries were determined using a Qubit double-stranded DNA high sensitivity Kit (Thermo Fisher Scientific) according to the manufacturer’s instructions. Libraries were diluted to appropriate concentrations for sequencing on an Illumina MiSeq, NextSeq, or NovaSeq.

**LTC Forward primer pool:** TCGTCGGCAGCGTCAGATGTGTATAAGAGACAG-[1N to 4N]-GAATCCAGCTAGCTGTGCAGC

**LTC Reverse primer pool:** GTCTCGTGGGCTCGGAGATGTGTATAAGAGACAG-[1N to 4N]-CCTTAGCCGCTAATAGGTGAGC

Clones with desired integration levels were thawed and subsequently expanded (48-well to 12-well to 6-well format), cryopreserved, and subjected to karyotype analysis.

#### Generation of tracing-competent PEtracer mESCs

PEmax-T2A-GFP homozygous mESCs harboring 35 unique LTCs using PiggyBac transposition were karyotyped prior to introducing slow, intermediate, and fast kinetics 24-mer pegArrays. Electroporations were conducted as above, where 2 x 10^6^ cells were nucleofected with 5 μg of transfer plasmid and 1 μg of Super PiggyBac Transposase Expression Vector (System Biosciences, PB210PA-1) using the CG104 protocol on the Lonza 4D-electroporation system. Cells were allowed to recover and expand over two weeks prior to FACS enrichment for the top 10% BFP-expressing cells. Individual clones were subsequently isolated and karyotyped prior to further use. Further, clones were evaluated by bulk sequencing for the presence and relative abundance of each pegRNA in the 24mer array. These PCRs were conducted as described above using two successive rounds of PCR with 2 μL of gDNA and 500 nM primer mix using the Phusion Green Hot Start II High-Fidelity PCR 2x Master Mix with the following cycle conditions: 98°C for 2 minutes, [98°C for 10 seconds, 70°C for 20 seconds, 72°C for 45 seconds]x29, 72°C for 2 minutes. PCR1 was conducted using a pool of forward and reverse primers where each primer was pooled in an equimolar fashion. All subsequent steps were carried out as described above in “Screening for Sorted PEmax-T2A-GFP mESCs with high numbers of integrated LTCs”.

**pegArray Forward primer pool:** TCGTCGGCAGCGTCAGATGTGTATAAGAGACAG -[1N-to-4N]-GAAAAAGTGGCACCGAGTCGGTG

**pegArray Reverse primer pool:** GTCTCGTGGGCTCGGAGATGTGTATAAGAGACAG-[1N-to-4N]-NGTTGGTTTAACGCGTAACTAGATAGAACCG

#### Long-read whole genome sequencing of homozygous PEmax-T2A-GFP, high-copy LTC mESCs

Following karyotype analysis, we sought to further characterize our lead line with 36 integrated LTCs using long-read whole genome sequencing. This line was thawed, expanded, and >10^6^ mCherry^+^ cells were isolated by FACS prior to genomic DNA isolation using the PacBio NanoBind CCB kit (102-301-900). Genomic DNA libraries for Oxford Nanopore sequencing were prepared using the Ligation Sequencing Kit V14 (SQK-LSK114; Oxford Nanopore Technologies) according to the manufacturer’s protocol. Briefly, high-molecular-weight genomic DNA was first assessed for quality and quantity using NanoDrop, and samples were normalized to the recommended input mass (2ug). DNA underwent end-repair and dA-tailing. Following a bead-based cleanup step to remove enzymes and short fragments, sequencing adapters supplied in the kit were ligated to the prepared DNA. Adapter-ligated libraries were purified using magnetic beads, and the final library was eluted in an elution buffer provided in the kit. Prepared libraries were loaded onto a PromethION flow cell with R10 nanopores and sequenced on the Oxford Nanopore P2 Solo platform. Sequencing runs were conducted using default parameters in MinKNOW software, with real-time basecalling enabled to generate FASTQ files for downstream analysis.

#### *In vitro* activation and collection for bulk sequencing

PEtracer mESCs were nucleofected and FACS-sorted for mCherry and GFP double-positive populations as described below (see “Generation of PEtracer mESC-derived chimeric embryos”). Following sorting, cells were plated back to culture instead of being used for aggregation. For time-course collection, independent nucleofections were performed for each biological replicate (n = 3). From each nucleofection batch, cells were cultured in parallel, with routine passaging every 2 to 3 days, and harvested at days 0, 2, 4, 6, and 8 post-sorting. At each time point, MEF-depleted cell pellets were collected for bulk sequencing. Genomic DNA was extracted from flash-frozen cell pellets using the DNeasy Blood and Tissue Miniprep kit (Qiagen, 69504), and LTCs were amplified and subjected to bulk sequencing as described above (see “Screening for Sorted PEmax-T2A-GFP mESCs with high numbers of integrated LTCs”).

#### Generation of PEtracer mESC-derived chimeric embryos

PEtracer mESCs (1-2 million cells) were nucleofected with 10 µg pCAG-Cre plasmid (Addgene plasmid #13775) using the P3 Primary Cell 4D-Nucleofector®X Kit L (Lonza, V4XP-3024) and the CG-104 program on a Lonza 4D-Nucleofector X system, following the manufacturer’s instructions and as previously described^165,166^. At 24 hours post-nucleofection, cells were trypsinized and sorted for mCherry and GFP double-positive populations using a BD FACSAria™II Cell Sorter (BD Biosciences) operated with BD FACSDiva software (v8.0.1). Cells were first gated to exclude MEFs, debris, and doublets based on forward and side scatters. Fluorescence was detected using a 510/20 filter for GFP and a 615/20 filter for mCherry. GFP-positive cells were defined relative to non-transfected control cells to establish background fluorescence, and the top approximately 80% of GFP-expressing cells within the mCherry^+^/GFP^+^ double-positive population were sorted and maintained on ice until aggregation. Subsequently, 6–8 healthy PEtracer mESCs were injected into precompaction 4- to 8-cell stage embryos derived from natural matings of CD1 mice for high-efficiency incorporation into the inner cell mass ∼24 hours later^167–169^. For tetraploidization experiments, 2-cell stage embryos were tetraploidized by electrofusion as described previously using the CF150/B Cell Fusion instrument (BLS)^48^. Injections were performed using ES blastocyst injection pipettes (BioMedical Instruments; VESsp-15-25-1-55; inner diameter 15 µm, bent angle 25°, bent length 1 mm, pipette length 55 mm; glass: BM100T-10P), in conjunction with an H.T. Laser 6 system (software version 6.5.1) and a CellTram injection system. Injected embryos were cultured overnight in EmbryoMax®Advanced KSOM Embryo Medium (Sigma-Aldrich, MR-101-D), overlaid in pretested embryo-grade mineral oil (Sigma-Aldrich M5310), and transferred into the uteri of pseudopregnant CD-1 strain females around 18 hours post-injection. Embryos were subsequently collected at the indicated developmental stages for downstream chimerism scoring and analyses.

#### Bulk sequencing analysis of chimeric E9.5 mouse embryos

Genomic DNA was isolated from intact chimeric E9.5 embryos using the DNeasy Blood and Tissue Miniprep kit (Qiagen, 69504). LTCs were amplified and sequenced as described above (see “Screening for Sorted PEmax-T2A-GFP mESCs with high numbers of integrated LTCs”). Edit saturation and LM distributions were calculated as previously described^43^.

#### Imaging-based documentation of experimental embryos

All whole embryo images captured prior to scRNA-seq were performed using a Zeiss Observer 7 with a 5x or 10x air objective.

#### scRNA-seq of experimental embryos

Whole embryos (E7.5-E10.0, 12 hr increments) were isolated and cleaned of maternal and extraembryonic tissue in a petri dish with 1x DPBS, then serially washed through several drops to remove maternal tissue. Complete embryos were initially screened by fluorescence imaging to identify high-grade chimeras, followed by documentation at 10x magnification. Following this initial selection, each embryo was individually transferred into a 200-500 µl drop of 0.2% BSA/PBS and dissociated into single cells by replacing the 0.2% BSA/PBS with an equal volume of TrypLE (Invitrogen) and incubated for 10 minutes at 37°C. After the incubation, embryos were mechanically dissociated using a P200 pipette and incubated >2x at 5-minute intervals. The cell suspension was quenched by serially adding and collecting 200 µl of 0.2% BSA/PBS and carefully resuspending the collected cells into a LoBind Eppendorf tube until the final volume was ∼1 mL (Eppendorf). A 1x Red Blood Cell lysis solution (Miltenyi Biotec) was used to treat cells from E8.5-E10.0 embryos. Cells were filtered into a new LoBind Eppendorf tube using a Scienceware Flowmi cell strainer, 40-70 µm (VWR) and centrifuged for 5 min at 1200 rpm at 4°C. Following centrifugation, ∼250 µl of supernatant was removed and cells were resuspended. This process was repeated to concentrate the cell pellet. Cells were resuspended in 0.2% BSA/PBS.

Approximately 5% of the cell suspension was then used to determine the overall percent chimerism of candidate embryos by flow analysis using a BD Celesta according to pre-defined gating parameters. Embryos with >70% chimerism were chosen as high-quality candidates for full embryo sequencing, followed by an estimation of the total cell count using a hemocytometer. For each embryo, the entire cell suspension was processed for single-cell RNA sequencing (10x Genomics, Chromium GEM-X Single Cell 3’ Chip Kit v4), distributing cells across multiple reactions to remain within the maximum recommended input of cells per reaction. During optimization, multiple PEtracer lines with distinct editing rates were tested and sequenced with recovery targets of ∼10-20,000 cells using the 10x Chromium NextGEM v3.1 3’UTR capturing technology (10x Genomics, Chromium Single Cell NextGEM v3.1). Single-cell libraries were prepared following the manufacturer’s protocol with cDNA (Step 2) and indexing amplification (Step 3) PCR cycle numbers of 10 each. Primary cDNA and final libraries were examined on a D5000 and D1000 TapeStation System, respectively (Agilent Technologies).

#### Target site library preparation

Following Step 2 of the 10x Genomics GEM-X workflow, a portion of amplified cDNA was used to generate target site libraries using pooled equimolar staggered forward primers and a common reverse primer (Key Resources Table). The optimal PCR1 cycle number was first determined by qPCR on a 1:8 cDNA dilution using KAPA HiFi HotStart ReadyMix (Roche, 07958935001) and SYBR Green (Invitrogen, S7563; 95°C 3 min, [98°C 20 sec, 65°C 15 sec, 72°C 15 sec]x40, 72°C 1 min); PCR1 was then performed on undiluted cDNA using the same primer set and cycling conditions for two fewer cycles than the qPCR plateau (typically 22 cycles). A second PCR on a 1:25 dilution of the PCR1 product incorporated Nextera i7 indices and sequencing adapters (Key Resources Table; 95°C 3 min, [98°C 20 sec, 72°C 30 sec]x14, 72°C 1 min). The final libraries were purified by 0.9x SPRI bead cleanup and eluted in Buffer EB. Libraries were assessed for size and purity by BioAnalyzer (Agilent) and quantified using a Qubit double-stranded DNA high sensitivity Kit (Thermo Fisher Scientific) to determine the concentrations. Libraries were diluted to 4 nM and sequenced on a MiSeq, NextSeq 2000, or NovaSeq X Plus where appropriate (Illumina). All PCRs were first monitored to determine optimal cycle number by qPCR by adding 0.6x SYBR Green (Thermo Fisher Scientific) to the PCR reactions.

## QUANTIFICATION AND STATISTICAL ANALYSIS

### Mapping LTCs using long-read whole genome sequencing

FASTQ files were aligned to the mouse reference genome (GRCm39) using the minimap2 (v2.28) aligner, and resulting alignments were processed to identify LTC integration sites. We leveraged the characteristic size of the LTC (∼7 kb) to detect insertions directly from individual long-read alignments. Specifically, we scanned CIGAR strings to identify large insertion events (6.9–7.3 kb) that were flanked on both sides by aligned sequence, ensuring confident anchoring of the insertion to the reference genome. For each such event, we recorded the genomic insertion position, extracted the inserted sequence from the read, and retained the longest qualifying insertion per read. Extracted insertion sequences were oriented to a consistent strand and parsed to recover the integration barcode sequence by identifying sequence motifs flanking the barcode region. The inferred sequences were then error-corrected by matching to a predefined intBC whitelist. This approach enabled direct linkage of individual LTC integrations to genomic coordinates. Final insertion calls were manually inspected using IGV to confirm alignment quality and validate integration sites.

### scRNA-seq data processing

All scRNA-seq data were processed using Cell Ranger (v9.0.1). Gene expression and lineage tracing libraries were each aligned to a custom reference genome built by appending 2,171 possible LTC sequences to the mouse GRCm39 assembly, enabling joint recovery of transcriptional and lineage information from the same cells. Gene expression was quantified against GENCODE vM37 gene annotations. Integration barcodes (intBCs) and lineage marks (LMs) were called from LTC-aligned reads using a custom Python script, as described previously^43^.

### scRNA-seq quality control

To generate a high-quality dataset with accurate lineage trees, we applied a multistep quality control workflow to remove low-quality cells, low-confidence LTC reads, ambient RNA-derived signal, and doublets. Because the abundance and quality of both transcriptome and LTC reads varied across captures, we used adaptive thresholding wherever possible. Specifically, when defining data-driven cutoffs, we fit a two-component Gaussian mixture model to the relevant one-dimensional distribution within each capture and removed observations corresponding to the lower peak of the distribution.

#### Removal of low-quality cells

We first filtered low-quality cells using transcriptome-derived QC metrics. For each cell, we calculated total UMI counts, total mitochondrial counts, and the fraction of mitochondrial transcripts. Within each capture, adaptive thresholding was applied to the distribution of mitochondrial counts among cells with detectable mitochondrial signal, and then to the distribution of total UMI counts among cells passing the mitochondrial filter and having less than 2% mitochondrial reads. Cells falling below either threshold were removed. A minimum total UMI cutoff of 5,000 was enforced across all captures, except for the shallowly sequenced preliminary data, where a threshold of 100 UMIs was used.

#### Removal of ambient LTC reads and PCR artifacts

We next filtered LTC alleles to remove low-support calls consistent with PCR artifacts or ambient RNA. For each allele, we calculated reads per UMI and retained only alleles with at least 5 UMIs and at least 4 reads per UMI. Within each cell–integration barcode (intBC) pair, we then removed alleles with weak support relative to the dominant allele. Specifically, we retained alleles only if their reads-per-UMI value was at least 25% of the maximum observed for that intBC in that cell, unless they had at least 10 reads per UMI, and we further required their UMI count to be at least 25% of the maximum UMI count within that same cell–intBC pair.

#### Identification of donor, host, and doublet cells

We then distinguished donor- and host-derived cells based on LTC expression and single-nucleotide polymorphisms (SNPs) detected in transcriptome reads. For each cell, total LTC UMI counts were summed and normalized within each capture by the median value among retained cells. Donor and host seed populations were defined from the upper and lower tails of the LTC UMI distribution, respectively. Host-specific SNPs were identified by comparison between host and donor seed populations and used to quantify host-like signal in each cell. Donor cells were then called using adaptive thresholding on the ratio of normalized LTC counts to normalized transcriptome counts. Cells classified as donor but showing excess host SNP signal were labeled as donor–host doublets. In captures where the host-specific SNP signal was weak and host cells were female, Xist expression was used as an orthogonal marker to identify donor–host doublets and validate our general thresholds.

We next used lineage information to identify donor–donor doublets based on conflicting LTC alleles. For each allele, we calculated the fraction of UMIs it contributed within its cell-intBC pair and used this information to select a dominant allele. We then quantified allele conflict rates as the fraction of reads assigned to non-dominant alleles. Baseline conflict rates were estimated across low-conflict cells and intBCs, and per-cell conflict fractions were recalculated using only integrations with low background conflict. Cells with allele conflict fractions greater than 3% were classified as donor–donor doublets.

Finally, we identified host-host doublets using an approach based on total UMI counts, as these cannot be resolved using SNPs or lineage information. Donor cells previously classified as singlets or doublets were ordered by normalized transcriptome counts, and the local doublet frequency was estimated using a rolling window. The normalized count threshold at which the local donor doublet frequency exceeded 50% was then used to define a cutoff for host doublets. Host cells above this threshold were labeled as host–host doublets. These and all previously identified doublets were removed from the dataset.

### Lineage tree reconstruction

Lineage trees were reconstructed independently for each embryo using the set of donor cells with >75% intBC detection (95% of cells passed this threshold; per embryo statistics in **Table S1**). For each cell, LTC alleles were converted into a character matrix in which rows corresponded to cells and columns corresponded to editable sites across intBCs. A scalable hybrid approach was used to construct the >100,000-cell lineage trees presented in this study, combining a greedy algorithm that resolves high-confidence splits (supported by ≥3 edits) at the top of the trees with a heuristic neighbor-joining algorithm to reconstruct subtrees within these clades.

To identify high-confidence splits, we applied a greedy recursive partitioning procedure based on shared mutations. Within each set of cells, candidate mutations were enumerated and ranked by prevalence. A seed mutation was selected, and additional mutations with highly overlapping cell sets (Jaccard similarity >0.9) were grouped to define a provisional split. This mutation set was refined by identifying mutations that were highly prevalent within the resulting subset of cells (present in >90% of cells, treating missing data as compatible), and cells matching this refined mutation set were assigned to a child clade. This procedure was applied iteratively, with each split required to be supported by ≥3 shared edits and to contain ≥1% of cells, ensuring that only well-supported divisions were resolved. Recursive splitting was limited to a maximum depth of 3 to prevent over-partitioning and to ensure that downstream reconstruction operated on sufficiently large clades. A small number of cells that did not match any high-confidence clade were removed as likely doublets.

Within each high-confidence clade, lineage trees were reconstructed using the heuristic neighbor-joining algorithm implemented in FastTree 2 (v2.1.11)^152^. This approach is highly scalable because it avoids explicit computation of the full pairwise distance matrix and has been shown to accurately reconstruct cellular lineage trees^170^. To run FastTree, lineage marks were encoded as amino acid states. Maximum-likelihood optimization and support-value estimation were disabled, such that topology was inferred using the underlying heuristic neighbor-joining procedure. A custom edit-distance matrix was supplied to reflect the structure of lineage recording data, in which only unedited-to-edited transitions occur, and missing values were retained during inference. Only characters detected in >50% of cells and with no single lineage mark represented in >95% of cells were used for subtree reconstruction.

Subtrees were rooted using a character-based outgroup strategy. For each clade, we identified the most frequent non-missing edited state across all characters and treated cells carrying this state as a proxy outgroup. For each edge in the unrooted tree, we evaluated the partition it defined by comparing the overlap between this proxy outgroup and the sets of descendant cells on either side of the edge using a Jaccard-like criterion. The tree was rooted at the edge that maximized this overlap, yielding a root placement consistent with the most globally shared mutation pattern.

Clade-level subtrees were subsequently combined to generate embryo-scale lineage trees. Because each embryo contained multiple independently seeded donor clones, the resulting trees naturally comprised multiple top-level branches corresponding to these clones. To enable clone-resolved analyses, embryo-level trees were partitioned into clone-level trees by splitting at the root, with each direct descendant of the root defining an independent clone-specific lineage tree.

### Branch length estimation

For each clone-level tree, ancestral lineage mark (LM) states were reconstructed using the Sankoff algorithm as previously described^43^. Following ancestral state inference, edges without any inferred mutations were collapsed to simplify the tree topology. This step removes branches that are not supported by any edits, merging nodes with identical inferred states. As a result, the trees contain multifurcations, where a single node gives rise to more than two descendants. These multifurcations reflect unresolved branching order due to insufficient mutational information and ensure that only biologically meaningful branches supported by at least one edit are retained.

Branch lengths were estimated using a maximum-likelihood framework implemented in ConvexML (v1.0.0)^153^, which models mutation accumulation as an independent exponential process across sites. Under this model, edits arise along each branch according to a Poisson-like process, such that the expected number of mutations is proportional to branch length scaled by site-specific mutation rates, providing a molecular clock–like interpretation in which branches with more edits correspond to longer durations. To account for heterogeneity in editing kinetics, we specified relative mutation rates (0.3, 0.4, and 1.2, repeated across sites). A pseudocount of 1 mutation per edge was included to stabilize estimates for branches with sparse signal. Missing and unedited states were explicitly incorporated into the model, and branch lengths were inferred jointly across the entire tree using convex optimization. Finally, branch lengths were rescaled such that the total depth of each tree matched the known sampling time of the embryo, and each internal node was assigned an inferred embryonic day.

### *In silico* benchmarking

Ground-truth lineage trees were simulated using the birth-death process implemented in Cassiopeia (v3.0.0)^151^. Waiting times between divisions were drawn from a log-normal distribution (σ = 0.1). To emulate the acceleration in proliferation observed over developmental time, the division rate was increased 2.5-fold (μ = 0.4 to μ = 1.0) for lineages more than two time units after the start of the simulation. Unless otherwise noted, each simulation was run until 10,000 extant cells were reached. Simulated lineage-tracing data with characteristics matching the empirical data (35 LTCs, 8 LMs, 50% edit saturation) were generated for each simulated tree using Cassiopeia’s stochastic tracing model.

#### Reconstruction algorithm benchmarking

To evaluate the scalability and accuracy of candidate reconstruction algorithms, trees were simulated at ten sizes spanning 200 to 200,000 extant cells, with 10 replicates per condition. Six algorithms were compared: Neighbor Joining, UPGMA, and Greedy solvers implemented in Cassiopeia; LAML-Pro (v0.1.0)^154^ maximum likelihood reconstruction (ultrametric = True, max iterations = 5000); and FastTree^152^, with and without the initial Greedy step used in the hybrid approach described above. For each run, reconstruction accuracy was quantified as the normalized Robinson-Foulds distance from the ground-truth topology. Wall-clock runtime and peak memory usage were also recorded. Runs were terminated if they exceeded a maximum runtime of one day.

#### Tracing-capacity benchmarking

To assess how recording capacity and detection efficiency affect reconstruction accuracy, 10,000-cell trees were simulated and reconstructed using the FastTree solver while sweeping the number of LTCs (5-50) and the missing-data rate (0-20%), with 10 replicates per condition. For each replicate, the underlying ground-truth tree was held fixed across conditions by seeding the simulation with the iteration index, such that only the character matrix size and missing-data rate varied. Reconstruction accuracy was quantified as the normalized Robinson-Foulds distance from the ground-truth topology.

#### Branch-length estimation benchmarking

Branch-length accuracy was evaluated on the true, rather than reconstructed, tree topology to isolate the performance of the estimation procedure from reconstruction error. Trees of 10,000 cells were simulated while sweeping the number of cassettes (5-50) and the variance in division timing, parameterized as the log-normal σ of the birth-waiting distribution (0.1-0.5), with 10 replicates per condition. Branch lengths were estimated using ConvexML, as described above, and LAML-Pro (minimum branch length of 0.001). Accuracy was quantified as the mean absolute error (MAE) between estimated and true simulated node times.

### Number of extant cells over time

The number of extant cells over time was calculated by traversing each reconstructed lineage tree and counting the number of branches present at each inferred timepoint. For each edge, the branch was considered extant from the time of the parent node (inclusive) until the time of the child node (exclusive). These counts were then summed across all branches to obtain the number of extant lineages at each timepoint.

### Percent of branches marked by an edit

We quantified the fraction of lineage branches supported by at least one edit by comparing the number of observed branches in the reconstructed trees to the total number of branches expected under a fully bifurcating topology. Because all mutationless edges were collapsed during tree reconstruction, every observed edge in the final tree corresponds to a lineage branch supported by one or more inferred edits. However, collapsing mutationless edges introduces multifurcations, in which a node has more than two children, representing unresolved sequences of cell divisions. Each multifurcation with k descendants implies the presence of *k-2* unresolved (hidden) branches that are not supported by detectable edits. The total number of branches was therefore defined as the sum of observed branches and these inferred hidden branches, and the overall fraction of branches marked by an edit was calculated as the ratio of observed edges to total branches. Equivalently, the fraction of unmarked branches is given by the inferred hidden branches divided by the total number of branches. These and other tree statistics are reported in **Table S1**.

To examine how this fraction varies over developmental time, we additionally computed the fraction of marked branches within discrete time intervals. Observed branches were assigned a time corresponding to the midpoint between their parent and child nodes. Hidden branches arising from multifurcations were similarly assigned approximate times based on the parent node and the distribution of child node times, with multiple hidden branches distributed between these values to reflect their implied ordering. Within each time bin, the number of observed (marked) and hidden (unmarked) branches was counted, and the percent of branches marked by an edit was calculated as the fraction of observed branches among all branches in that interval.

### Normalization and highly-variable gene selection

Single-cell analysis was performed using Scanpy (v1.11.5)^147^. Gene expression counts were normalized by scaling total UMIs per cell to 20,000 (selected based on mean UMIs/cell) followed by log1p transformation, and cell cycle scores were computed using curated S phase and G2/M gene sets. To identify highly variable genes while minimizing confounding effects, we excluded genes exhibiting strong bias with respect to donor versus host identity, developmental stage, cell cycle, or sex. Donor- and host-biased genes were identified by differential expression (t-test), and the top 500 genes ranked by absolute effect size were removed. For each developmental stage, the top 20 stage-enriched genes were also excluded. In addition, all cell cycle genes and genes located on the X and Y chromosomes were removed, and genes expressed in fewer than 1,000 cells were filtered out. Highly variable genes were then selected from this filtered gene set using a variance-based approach that accounts for mean expression, selecting the top 5,000 genes while controlling for batch effects across developmental stages; these genes were used for downstream dimensionality reduction and clustering analyses.

### Embedding with scVI and UMAP

Cells were embedded using scVI (scvi-tools v1.4.0)^148^, which learns a low-dimensional latent representation of gene expression while accounting for technical and biological variability. The model was trained using 50 latent dimensions, 2 hidden layers of 1024 hidden units, 15 epochs, and a batch size of 512, with developmental stage specified as a batch covariate. UMAP embeddings were generated using RAPIDS single-cell (v0.13.2)^149^ by constructing a 50-nearest neighbor graph from the scVI latent space and applying UMAP with default parameters and a fixed random seed for both global and subcluster visualizations.

### Transcriptional similarity between timepoints and with reference datasets

We measured transcriptional similarity between datasets by estimating distributional distances between cell populations in an integrated scVI latent space. We first co-embedded datasets using the same scVI parameters as above, with the dataset label specified as a batch covariate. We then computed transcriptional similarity using Sinkhorn similarity, defined as a normalized Sinkhorn divergence, where the minimum divergence across all pairs of embryos was scaled to 1 and the maximum divergence was scaled to 0. Sinkhorn divergence was estimated using p=2 and regularization of ε = 0.025, and the mean across 50 minibatches of size 512. We used the Sinkhorn implementation from geomloss^171^.

### Cell type annotation

Cell types were annotated at the level of cell subtypes using a combination of reference-based label transfer, manual curation, and automated approaches. Where a single anatomical tissue comprises cells of more than one embryonic origin, cell type labels refer to the molecularly defined population rather than the composite tissue, consistent with usage in other single-cell atlases^22^. For example, “pharyngeal arch mesenchyme” denotes the neural crest-derived mesenchymal population, whereas the mesoderm-derived cells of the pharyngeal arches are annotated separately as “pharyngeal mesoderm”. Initial annotations were obtained by transferring labels from published mouse embryo reference atlases, including Pijuan-Sala et al., 2019^21^ and Qiu et al., 2024^22^, based on similarity in gene expression profiles. These labels were refined through manual inspection and cluster-level analysis using cellxgene (v1.3.0), guided by canonical marker genes and literature review. For each cluster, marker genes were identified in two categories: “global” markers distinguishing the cluster from all other cells, and “local” markers distinguishing the cluster from the three most transcriptionally similar cell subtypes. In parallel, automated annotation was performed using Claude Opus (v4.5) by providing cluster-level marker genes as input. Final subtype annotations were determined by integrating evidence from label transfer, manual curation, and AI-based predictions; no annotation was assigned by label transfer or AI prediction alone. Because these lines of evidence were combined through expert judgment rather than applied as sequential automated steps, final labels do not correspond to discrete AI-only or transfer-only assignments. In **Table S2,** claude_description and claude_rationale columns were generated using Claude Opus (v4.5) and provided as lightly reviewed model output to aid interpretation. Subtypes were subsequently grouped into broader cell types and further annotated with germ layer and developmental domain information.

### Quantifying germ layer mismatch

To estimate the rate of leaf misplacement in our lineage trees, we quantified how often a cell’s germ layer disagreed with that of its siblings. For each clone-level tree, ancestral germ-layer states were reconstructed using the Fitch–Hartigan algorithm and the mismatch rate was defined as the fraction of annotated leaves whose germ layer differed from the reconstructed state of their parent node. Presomitic mesoderm and sacral spinal cord cells were excluded prior to this calculation, as their axial progenitor origin places them adjacent to cells of the other germ layer even when correctly reconstructed. This yielded a mean mismatch rate of 0.97% for our E9.5 trees.

### Calculating cell fate bias

Cell fate was defined operationally as the distribution of cell states among descendant cells within the reconstructed lineage trees. For each internal node, a fate distribution was computed by tallying the number of descendants belonging to each annotated cell type (or other fate category, such as germ layer). To characterize how these fate distributions evolve over developmental time, we split the tree into discrete time bins (0.5-day intervals for all analyses) and assigned each node’s contribution across these bins based on the temporal span of its lineage. Specifically, each node contributes its fate distribution to all timepoints between the time of its parent (inclusive) and its own time (exclusive). This reflects the biological interpretation that a branch defined by a node persists over the interval from its birth until a division event gives rise to child nodes. Thus, this approach captures how fate bias accumulates along continuous developmental trajectories rather than at discrete branching points.

Fate bias was quantified using a normalized entropy-based score applied to each node’s fate distribution. Specifically, for a node with descendant proportion vector *p*, the fate bias score was defined as *1 - H(p)/H(q)*, where *H(p)* is the entropy of the node’s distribution and *H(q)* is the entropy of the root distribution for the corresponding tree, both of which are estimated via a plug-in estimator. This score ranges from 0 (no bias relative to the root distribution) to 1 (complete restriction to a single fate). To summarize trends over time, a weighted mean fate bias was calculated within each time bin, where each node was weighted in proportion to its number of descendant cells. Statistical significance of fate bias at each timepoint was assessed using permutation testing with 100 permutations: descendant fates were resampled under a null model that preserves total descendant counts per node and the overall root distribution, and weighted mean fate bias was recomputed for each permutation. Observed values were then compared to this null distribution to obtain p-values, providing a measure of when lineage-wide fate bias becomes significantly elevated above what would be expected by random subsampling. The black line in plots with permutation testing indicates the mean across all permutations.

### Identifying fate-restricted clades

Fate-restricted clades were identified from reconstructed lineage trees based on the fate distribution of each node. For each fate category, such as a given cell type or germ layer, we defined a node as fate-restricted if at least 90% of its descendant cells belonged to that fate. To avoid overcounting nested clades, fate-restricted clades were selected using a two-pass procedure applied separately for each fate. First, nodes were evaluated from the leaves upward to identify all nodes whose descendant composition met the 90% threshold. Second, nodes were traversed from the root downward, and the highest qualifying node for each fate was selected; once a node was selected, its descendants were excluded from further consideration for that same fate. This yielded the earliest detectable ancestor of each fate-restricted clade while preventing descendant nodes from being counted repeatedly. The 90% threshold was used to account for minor labeling and tree reconstruction errors–in practice, the vast majority of fate-restricted clades are 100% pure for the given fate.

Multifurcations were handled explicitly because they reflect unresolved branching order rather than true simultaneous multi-way divisions. For nodes with more than two children, if multiple child branches independently qualified for the same fate, the parent node was identified as the fate-restricted ancestor for those children. Biologically, this assumes the most parsimonious scenario—that fate restriction occurred once at an unresolved split, rather than independently in each child branch. This approach avoids artificially inflating the number of fate-restricted clades due to limited lineage resolution, but may miss true cases where multiple independent fate-restriction events occur within a multifurcation. Accordingly, the number of fate-restricted clades should be interpreted as a lower bound on the number of progenitors for a given fate due to unresolved branches and embryos not being 100% chimeric.

### Visualizing hierarchical fate restriction over time

To visualize the progressive restriction of lineages over developmental time, fates were organized as an ordered hierarchy of leaf annotations running from coarse to fine, and each node was assigned the label at the most specific level at which a single label accounted for >90% of its annotated descendants, matching the threshold used to define fate-restricted clades above, or labeled uncommitted if no level qualified. Cells lacking an annotation at a given level were excluded from both the numerator and the denominator at that level, acting as wildcards compatible with any label. A subsequent top-down pass enforced monotonic restriction, with each node inheriting its parent’s commitment unless it qualified at a strictly more specific level.

Transitions between commitment states were counted across half-day intervals, using the branch-extant convention described above. Each branch at a given timepoint contributes one transition from the branch at the preceding timepoint, so a source branch that divides within an interval contributes multiple transitions. Commitment annotation was always performed on the complete trees, so that all descendants contribute to the composition of each internal node. However, when restriction toward a particular lineage was displayed, transition counts were computed on the subset of leaves belonging to that lineage together with their ancestral branches, so that the diagram traces the commitment history of the lineage of interest rather than the composition of the embryo as a whole. Counts were pooled across clone-level trees and embryos, normalized within each timepoint, and rendered as a Sankey diagram in which ribbon widths are proportional to the fraction of extant lineages following each path.

This framework was applied to several fate hierarchies. Germ-layer restriction used a single-level hierarchy of ectoderm, mesoderm, and endoderm, with epiblast and primordial germ cells excluded. Notochord restriction used a two-level hierarchy of germ layer and notochord identity, with notochord treated as a wildcard at the germ-layer level. Endothelial restriction likewise used a two-level hierarchy of mesoderm subtype and endothelial identity, with endothelium treated as a wildcard at the first level. Heart-field restriction used a three-level hierarchy of germ layer, heart, and heart field, where heart-field labels were defined as pSHF (IFT, atria, pSHF), aSHF (RV, OFT, aSHF), JCF, epicardium, endocardium, and FHF (LV, AV canal). Neural-ectoderm restriction used a three-level hierarchy of germ layer, neural ectoderm, and neural region, with cell types not corresponding to a region (e.g., roof plate) treated as wildcards at the region level. Neural-crest restriction used a four-level hierarchy of germ layer, neural ectoderm, biased output, and differentiated output, where biased-output categories were defined as DRG-biased (DRG together with other non-autonomic neural crest), autonomic-biased (autonomic together with other non-DRG neural crest), and bipotent (both DRG and autonomic, together with other neural crest). Midbrain–hindbrain boundary restriction used a hierarchy of neural ectoderm, an intermediate level comprising two alternative groupings of r1 with its neighbors (r1 with r2–r4, or r1 with midbrain), and r1 alone, with isthmic organizer cells merged into r1.

### Ancestral linkage analysis

To assess how closely each cell was related to a given target category (e.g., a cell type or germ layer), we computed an ancestral linkage score based on lineage distance within the reconstructed trees^57^. For each cell, we identified the most recent common ancestor (MRCA) shared with the nearest cell belonging to the target category and used the depth of this ancestor as a measure of relatedness, with deeper MRCAs indicating more recent shared ancestry. To account for differences in the abundance and distribution of the target population within the tree, this value was normalized by subtracting a permuted background expectation generated by randomly reassigning non-target labels while preserving the target population. The resulting ancestral linkage score therefore reflects how much more closely related a given cell is to the target population than expected by chance. To summarize these relationships at the population level, per-cell linkage scores were aggregated by grouping cells according to a source category of interest (e.g., cell type) and computing the mean linkage within each group.

To provide a global view of lineage relationships across categories (e.g., cell types or germ layers), this framework was extended by iteratively treating each category as the target and computing mean ancestral linkage from all source categories to that target, yielding a pairwise category-by-category matrix. As these relationships are inherently asymmetric, the resulting matrix was symmetrized by averaging reciprocal source–target values. Statistical significance was assessed using permutation testing (100 permutations), applying the same non-target shuffling scheme described above. To quantify variability, pairwise linkage was computed independently for each embryo, and variation was evaluated across biological replicates from the same developmental timepoint. Mean pairwise linkages at E9.5 are reported in **Table S3** with all statistically significant values >0.5 annotated with a proposed rationale and novelty score (0 = textbook to 5 = completely novel) generated by Claude Opus (v4.8).

### Leave-one-integration-out jackknife stability analysis

To confirm that the reconstructed trees and the lineage statistics derived from them are stable to the particular set of integrations recovered, we performed a leave-one-integration-out jackknife for each E9.5 embryo (E9.5-R1, E9.5-R2, E9.5-R3). For each iteration, one intBC was removed from the character matrix and the tree was reconstructed from the remaining sites using the full pipeline described above. Topological stability was quantified as the normalized Robinson-Foulds distance between each jackknife tree and the tree reconstructed from the complete character matrix. To establish that downstream conclusions are likewise insensitive to intBC composition, fate-restricted clades and ancestral linkage were recomputed for each replicate as described above, then the number of fate-restricted clades, mean clade output, and mean restriction time were summarized per germ layer and pairwise germ-layer ancestral linkage was averaged across replicates within each embryo. Variability across replicates reported as the 95% percentile interval.

### Simulating topological error

To assess the sensitivity of our clade- and linkage-based statistics to residual reconstruction error, we simulated the effect of leaf misplacement on trees with known fate structure. Trees of 10,000 extant cells were simulated as described in the “*In silico* benchmarking” section and germ-layer identities were assigned by simulating commitment down a predefined trajectory using Cassiopeia^151^. In this trajectory, the uncommitted state gives rise to endoderm (probability 0.2) or a mesectoderm intermediate (probability 0.8) at scaled time 0.6, with the intermediate resolving into ectoderm (probability 0.6) or mesoderm (probability 0.4) at scaled time 0.7. Topological error was then introduced by permuting germ-layer labels among a randomly chosen fraction of leaves (0-20%), emulating cells placed at the wrong tip of an otherwise correct tree. As the nominal permutation rate overstates the perturbation actually introduced, each condition was instead characterized by its germ layer mismatch rate, calculated as described above. Fate-restricted clades were then identified and the total number of clades, mean clade output, and mean restriction time were expressed as percent deviations from the unperturbed labeling of the same topology.

### Fate bias pattern analysis

To characterize recurrent patterns of early fate bias, we analyzed cell type–level fate distributions at ancestral nodes within reconstructed lineage trees. For each node, descendant cell types were aggregated to generate a count vector, which was then normalized to proportions to define the node’s fate distribution. Only the composition of descendant cell types was considered; the topology of the descendant lineage tree was not used in this analysis. These distributions were expanded across developmental time by assigning each node to all time bins spanned by its incoming branch, as described above, using half-day time intervals. Analyses were restricted to nodes from E9.5 embryos, focusing on early developmental timepoints (≤E7.0) and requiring a minimum of 50 descendant cells per node to ensure robust estimation of fate composition.

For each selected node, fate distributions were used as feature vectors to construct a nearest-neighbor graph (k = 20), followed by dimensionality reduction using UMAP and clustering using the Leiden algorithm (resolution = 0.5). Clusters identified in this space represent recurrent “fate bias patterns”, defined as groups of ancestral nodes with similar descendant fate compositions. Pattern identities were assigned based on Leiden cluster membership and analyzed to identify common and distinct patterns of early fate bias across embryos.

To quantify the temporal dynamics of fate bias within each pattern, we computed a weighted fate bias score across time. For each node, the fate bias score was multiplied by the number of descendant cells to obtain a weighted contribution, reflecting both the strength of bias and the size of the corresponding lineage. These values were then summed across all nodes within each pattern and time bin and normalized to the maximum observed value across patterns. This analysis captures how strongly and how extensively each pattern contributes to fate bias over developmental time, enabling comparison of the timing and magnitude of bias emergence across distinct fate bias patterns.

### Sibling output analysis

To investigate the fates of sibling lineages, we analyzed the outputs of clades adjacent to fate-restricted nodes within the reconstructed lineage trees. For each fate-restricted clade, the sibling clade was defined as the set of descendant cells arising from the nearest branching point not assigned to that fate. For directly identified clades, sibling clades were defined as the descendants of the parent node excluding the focal clade, whereas for clades identified at unresolved multifurcations, the sibling clade corresponded to the descendants of the same node. To ensure comparable lineage depth and to avoid inclusion of overly broad clades, sibling groups with substantially larger descendant populations than the focal clade were excluded. The composition of sibling clades was then quantified in terms of cell type or germ layer identity and compared to the overall background distribution across the dataset. This analysis provides a measure of how frequently specific fates co-occur within closely related lineages, offering insight into shared developmental origins and spatial or lineage coupling between emerging cell populations.

### Clade co-occurrence

To quantify the extent to which distinct categories (e.g., cell subtypes or heart fields) co-occur within fate-restricted clades, we computed a pairwise co-occurrence statistic. Each clade was treated as a unit, and for each clade we recorded the presence or absence of each category of interest. This yielded a binary clade-by-category matrix indicating whether a given category was represented among the descendants of each clade. Co-occurrence was quantified using pointwise mutual information (PMI), which measures the deviation of observed co-occurrence frequency from that expected under independence. Specifically, for each pair of categories *i* and *j*, PMI was defined as

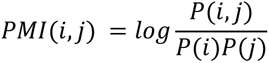

where *P(i,j)* is the probability that both categories are present within the same clade, and *P(i)* and *P(j)* are their marginal probabilities across clades. Positive PMI values indicate that two categories co-occur more frequently than expected by chance, whereas negative values indicate mutual exclusivity.

### Quantifying neural crest fate restriction dynamics

To characterize fate restriction dynamics within the neural crest (NC), we analyzed trunk NC-restricted clades and quantified the emergence of differentiated outputs over time. For each NC-restricted clade, descendant cells were annotated by differentiated output (DRG versus autonomic), and clades were binned by inferred restriction time using half-day intervals. Within each time bin, we calculated the probability that a clade contained each differentiated output, as well as the probability of producing multiple outputs (bipotency), defined as the fraction of clades containing both categories. To assess whether co-occurrence of distinct outputs deviated from statistical independence, we compared the observed frequency of bipotent clades to that expected under independent occurrence of each output type, calculated as the product of their marginal probabilities. Because larger clades are more likely to contain both outputs by chance, we additionally applied a size-stratified permutation test to E9.5 clades producing at least one differentiated output: clades were divided into ten equal-frequency bins by number of descendant cells, and the presence or absence of each output was independently permuted among clades within each bin (100 permutations), preserving the frequency of each output within each size stratum while breaking any association between them. The observed bipotent fraction was then compared to this null distribution, summarized as the 95% percentile interval.

### Quantifying within-clade transcriptional similarity

To assess whether heart-restricted clades exhibit additional transcriptional structure beyond annotated cell subtypes, we quantified the similarity of descendant cells within each clade using a low-dimensional embedding. For each clade, pairwise transcriptional similarity between cells was computed from the scVI latent space using cosine similarity, and the mean pairwise similarity across all cell pairs within the clade was used as a summary statistic.

To determine whether observed within-clade similarity exceeded that expected by chance, we performed a stratified permutation test. For each clade, a null distribution was generated by randomly sampling sets of cells of equal size from the most common cell subtype within the clade. Mean pairwise similarity was computed for each permuted set, and the observed value was compared to this null distribution to obtain a p-value and z-score. Clades with significantly higher similarity than expected (FDR-corrected across all clades) were interpreted as exhibiting additional transcriptional substructure, consistent with the presence of lineage-defined sub-domains within cell subtype identities.

### Identifying gene programs with Hotspot

Lineage-associated gene programs were identified using Hotspot (v1.1.3)^98^. Neighbor weights were computed for the five nearest neighbors in the lineage tree using an exponential kernel with σ = 2.0 applied to inferred tree distances. Normalized expression values were used to calculate gene-level local autocorrelation under the ‘normal’model with 30 parallel jobs. Here, C denotes a normalized, correlation-like measure of the extent to which a gene’s expression covaries among neighboring cells in the lineage tree, with higher values indicating stronger positive local autocorrelation. Genes with local autocorrelation C >0.1 were retained for pairwise analysis.

Pairwise local correlations were then computed among retained genes using Hotspot. Genes were grouped into coherent programs based on their local correlation Z-scores. Coarse clusters were first identified by hierarchical clustering of the Z-score matrix, and gene programs were subsequently defined within each cluster using an agglomerative procedure in which genes were merged when their mean pairwise Z-score was ≥80. Programs containing at least four genes were retained. This approach identified lineage-associated gene programs whose expression covaries among closely related cells in the lineage tree.

Gene program activity was scored in each cell as described previously^172^. For each program, the score was computed as the average expression of program genes minus the average expression of expression-matched control genes sampled from 25 expression bins. Program scores were used for downstream visualization and lineage-based analyses.

Gene program annotation was performed in a fully automated manner using Claude Opus (v4.6) by providing, for each program, the constituent genes and the set of cell types in which activity was enriched. The model generated a concise name and a brief biological description summarizing the dominant functional theme of each program. These annotations, reported in **Table S5**, were not manually curated or experimentally validated and should be considered provisional, model-derived summaries intended to aid interpretation rather than definitive biological assignments.

### Inferring ancestral program scores and rates of change

Gene program activity was imputed for internal nodes of the E9.5 lineage trees as the mean score of descendant cells, enabling reconstruction of program dynamics along lineage trajectories. To quantify these dynamics, 100 cells per cell type were randomly sampled, and program scores were computed across half-day intervals from root to leaf, with each internal node assigned a score spanning the interval between its parent and itself, as described above. Mean program activity over time was then obtained by aggregating trajectories within each cell type.

Rates of change in program activity were estimated using a smoothed finite difference of program scores over time. Scores were first smoothed with a centered rolling mean (window = 2 bins), and derivatives were then estimated at each time point as the difference between the subsequent and preceding smoothed values divided by the elapsed time between them (i.e., a centered two-point slope). To generate the rate-of-change heatmap, mean rates were computed across cell types in which a given program reached a maximum score >0.5; if no cell types met this threshold, the highest-scoring cell type was used. For comparability across programs, rate profiles were normalized by their maximum value.

### Estimating transcriptional divergence

To quantify how rapidly transcriptional states diverge among closely related cells, we measured the mean distance in scVI latent space between each cell and its nearest lineage neighbors, defined as cells sharing a common ancestor within a fixed temporal window (maximum lineage distance of 3, corresponding to approximately 1.5 days). For each cell, Euclidean distances to these neighbors were averaged to obtain a transcriptional divergence score, where lower values indicate that closely related cells remain similar and higher values indicate rapid divergence following lineage separation. This metric reflects local dispersion in transcriptional state among related cells rather than directional change. To compare divergence across cell types, per-cell scores were aggregated by taking the median within each annotated cell type.

### Inferring spatial positions using reference spatial-transcriptomics

To infer spatial positions of cells along the anterior-posterior (A-P) and dorsal-ventral (D-V) axes, we leveraged a published spatial transcriptomics dataset from a sagittal section of an E13.5 mouse embryo^114^. This section was selected because it is the earliest stage in the reference dataset that captures the full axial extent of the neural ectoderm within a single midline plane. Because we define the A-P coordinate as the normalized cumulative distance along a continuous anterior-to-posterior midline trajectory (see below), complete axial coverage is required for the coordinate to be consistently defined across the full axis. The larger embryo size and greater cell number at E13.5 additionally provide more labeled positions for training the A-P and D-V regressors. Although the developmental stage does not exactly match that of our dataset, key spatial patterning programs–particularly Hox gene expression–are established early and remain broadly stable across stages.

Within the spatial dataset, brain regions were manually annotated based on anatomical definitions from the Allen Brain Atlas^173,174^. A continuous midline trajectory was defined by tracing the center of the neural ectoderm from anterior to posterior. Each spatial location was projected onto the nearest point along this trajectory to obtain an A-P coordinate, defined as cumulative distance along the path and normalized to the interval [0,1]. D-V coordinates were computed as the signed perpendicular distance from each cell to the midline, with sign determined by the local normal vector; these values were also normalized to [0,1].

To extend these spatial coordinates to single-cell data, we trained regression models to predict A-P and D-V positions from gene expression using the spatial reference. Spatially informative genes were selected based on prior knowledge of developmental patterning systems: A-P prediction relied primarily on Hox and other axial patterning genes, while D–V prediction used canonical dorsal–ventral markers in the spinal cord. For each axis, a random forest regressor was trained with five-fold cross-validation, and performance was evaluated using cross-validated R^2^. The trained models were then applied to the single-cell dataset to infer spatial coordinates for each cell. These inferred positions recapitulated known spatial organization across the developing brain and spinal cord and were used for downstream analyses of spatial patterning and lineage relationships.

### Heritability and bias for spatial positions

To quantify the heritability of spatial identity, we assessed the similarity of inferred spatial positions between closely related clades within reconstructed lineage trees at E9.5. For each internal node corresponding to a neural-restricted clade, we identified its nearest neighboring internal node (also restricted to neural clades) using a graph-based nearest-neighbor approach on the tree. We then compared the mean inferred A–P and D–V positions of the descendant cells for each clade to those of its nearest neighbor. Heritability was quantified as the correlation (R²) between these paired values, providing a measure of how strongly spatial identity is preserved across nearby branches in the lineage tree.

To determine when spatial biases first emerge during development, spatial coordinates were inferred for internal nodes by averaging the positions of their descendant cells. These ancestral spatial scores were then expanded across time by assigning each node’s value to all time bins spanning from its parent to itself, as described above. Temporal trends in spatial bias were visualized by grouping nodes according to the spatial position of their descendant cells and computing the mean ancestral position within each time bin. This analysis captures the progressive emergence of spatial organization, revealing when lineage-associated biases along the A–P and D–V axes first become detectable over developmental time. Notably, the most posterior value predicted in our data was ∼0.8, consistent with the additional posterior growth that occurs from E10.0 to E13.5.

### Quantifying the dynamics of axial elongation

Throughout these analyses, “axial progenitor” denotes a transcriptionally defined *Sox2*⁺/*T*⁺ cell, whereas “neuromesodermal progenitor” is reserved for a progenitor demonstrated to be bipotent by lineage.

#### Sibling clade analysis

To identify and characterize bipotent axial progenitors, we analyzed the lineage relationships and descendant compositions of neural-restricted clades in E9.5 embryos. For each neural-restricted clade, sibling clades were defined as described above, and the distribution of non-neural developmental domains among sibling descendants was quantified. These distributions were examined as a function of the mean A-P position of the corresponding neural-restricted clade to assess how lineage outputs vary along the body axis. To visualize transcriptional relationships among these populations, we constructed a UMAP embedding, as described above, of cells belonging to neural-restricted clades and their sibling lineages. This visualization was restricted to posterior neural clades (mean A-P score >0.6), revealing the transcriptional continuum linking neural ectoderm, axial progenitors, and mesodermal derivatives.

#### Timing of bipotent clades

To quantify the timing of neural–mesodermal fate separation along the A-P axis, we calculated the inferred time of the most recent common ancestor (MRCA) with mesoderm for every neural ectodermal cell and summarized these values as a smoothed mean along the A-P axis. To further stratify these relationships, lineage trees were partitioned at discrete timepoints between E7.0 and E9.0 using inferred branch lengths (0.2-day intervals). At each timepoint, clades were defined by cutting the tree at the corresponding depth, and each clade containing at least two neural ectoderm cells was classified as bipotent if it also contained mesodermal descendants. The fraction of bipotent clades was then calculated as a function of both clade depth and the mean A-P position of their neural descendants. To relate these lineage-defined bipotent clades to transcriptionally defined axial progenitors, we focused on clades identified at E8.0 and calculated the fraction that contained at least one cell annotated as an axial progenitor.

#### Relative inferred position of axial progenitor output

To compare the spatial outputs of bipotent progenitors, we analyzed the inferred A-P positions of neural and mesodermal descendants. Because paraxial mesoderm represents the primary mesodermal output, we focused on these cells and calculated A-P scores as described above. For mesodermal cells, A-P scores primarily reflect Hox gene expression domains rather than precise anatomical position, as these domains are modestly shifted relative to those in the neural tube^130^. For each neural and paraxial mesoderm cell, we determined the inferred time of the MRCA between these two populations. Cells were grouped by this divergence time (E7.0–E9.0, 0.2-day intervals), and the mean A-P position of the neural and mesodermal populations was calculated within each bin, enabling comparison of the spatial positioning of lineage-related neural and mesodermal outputs over developmental time.

#### Quantifying axial progenitor bias

To determine whether the bipotency of the axial progenitor pool reflects uniformly bipotent progenitors or a mixture of fate-biased progenitors, we identified axial progenitor-restricted clades as described above and characterized the germ-layer output of their sibling clades. Each sibling clade was classified as ectoderm-biased, mesoderm-biased, or bipotent according to whether its descendants comprised ectodermal, mesodermal, or both germ layers. To assess whether the observed frequency of bipotent clades departed from statistical independence, we applied the size-stratified permutation test described above for the neural crest: analysis was restricted to clades producing at least one germ-layer output, clades were divided into ten equal-frequency bins by number of descendant cells, and the presence or absence of each output was independently permuted among clades within each bin (100 permutations). The observed bipotent fraction was compared to the resulting null distribution, summarized as the 95% percentile interval.

#### Clonal persistence of the axial progenitor pool

To assess whether the axial progenitor pool is continuously replenished from many early founders or declines in clonal diversity over time, lineage trees from E8.0, E8.5, E9.0, and E9.5 embryos were each cut at an inferred time of E7.0. For each stage, we calculated the fraction of the resulting clades that contained at least one cell annotated as an axial progenitor at the time of sampling, providing a measure of how many independent E7.0 lineages remained represented in the progenitor pool at each subsequent stage.

#### Proliferation score

To estimate relative proliferation rates across cell populations, we quantified the number of close lineage relatives for each cell within reconstructed lineage trees. Close relatives were defined as cells sharing a common ancestor within the previous 24 hours based on branch length estimates. While this metric is noisy at the single-cell level due to imperfect branch length estimation and subsampling, averaging across thousands of cells representing each cell type yields robust estimates of relative proliferation. This metric reflects the net expansion rate of a lineage over the preceding 24 hours, integrating both cell division and cell loss. Thus, populations with higher proliferation scores have generated more surviving close relatives over that interval, whereas lower scores may reflect slower proliferation, greater cell loss, or both.

## ADDITIONAL RESOURCES

These data can be interactively explored at the following link: mela.wi.mit.edu

## Supplemental information

**Figure S1.**
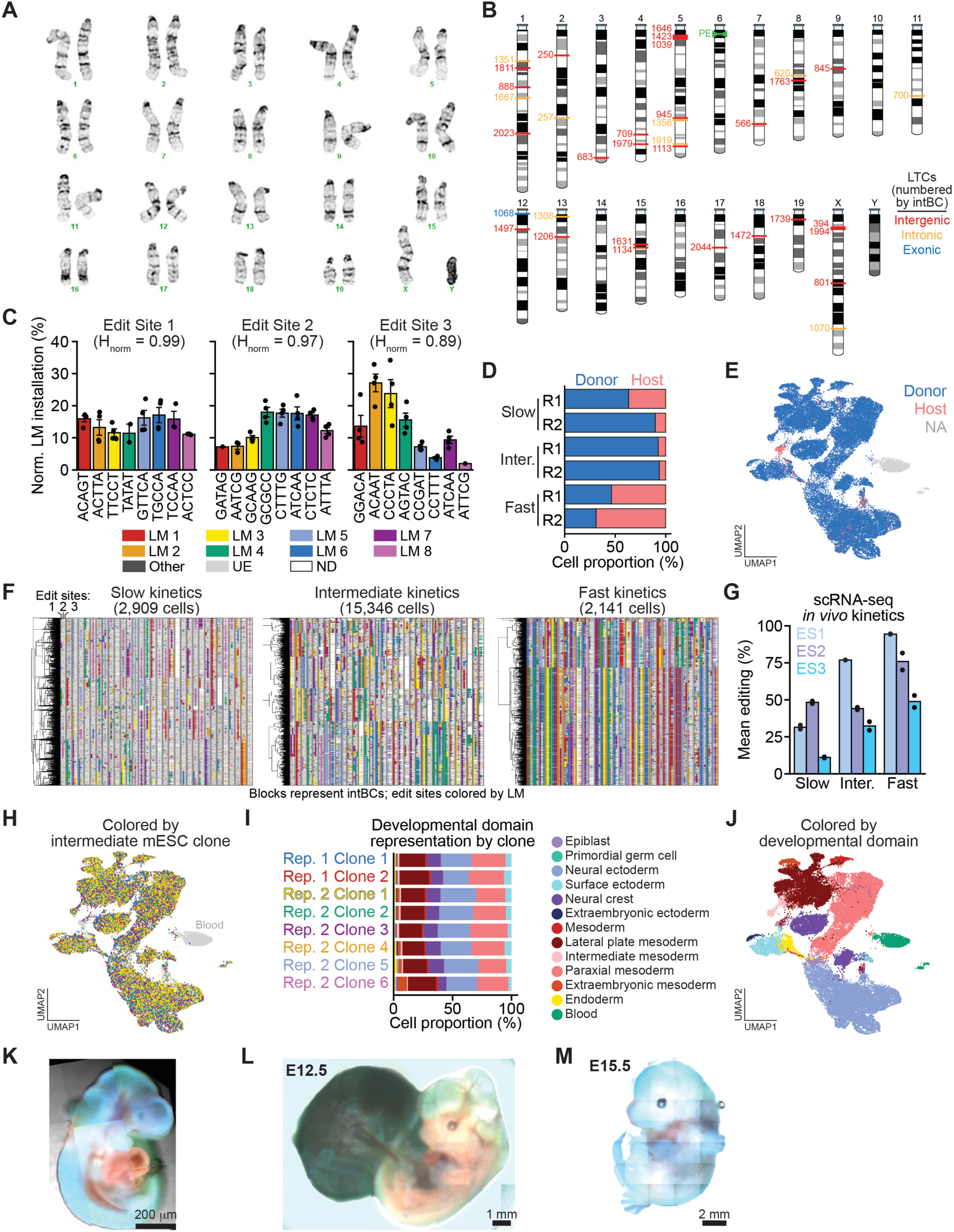
Validation of PEtracer for mouse developmental studies, related to Figure 1. (**A**) Karyotype analysis of mouse embryonic stem cells harboring a homozygous Rosa26 knock-in of Cre-activatable PEmax-T2A-GFP and 35 unique, genomically integrated lineage tracing cassettes (LTCs). (**B**) Genomic positions of integrated LTCs numbered by integration barcode (intBC) and colored by genomic context: intergenic (red), intronic (orange), a single exonic integration in *Ncoa1* (blue), and Rosa26 PEmax-T2A-GFP knock-in on chromosome 6 (green). (**C**) Normalized (Norm.) lineage mark (LM) installation efficiencies for all eight LMs at each of the three edit sites (ESs) in intermediate-kinetics E9.5 embryos from bulk sequencing. Bars represent the mean of four biological replicates ± standard error of the mean; normalized entropy (Hnorm) values provided for each ES. LMs are colored and ordered by their position in the pegArray. (**D**) Proportion of tracing donor cells (blue) and non-tracing host cells (salmon) assessed by single-cell RNA sequencing (scRNA-seq) for two biological replicates of E9.5 embryos generated from slow-, intermediate-(inter.), and fast-kinetics mESC lines. (**E**) UMAP embedding of two intermediate-kinetics E9.5 embryos colored by donor and host cell identity with primitive erythrocytes excluded. (**F**) Cellular phylogenies and character matrices for representative sub-sampled slow-, intermediate-, and fast-kinetics E9.5 embryos. Character matrix blocks represent LTCs with distinct intBCs and are colored by LM identity at ESs 1-3; cell number is indicated for each tree. (**G**) *In vivo* LM installation rates for donor cells from the two E9.5 embryo replicates in (D). (**H**) UMAP embedding from (E) colored by contributing mESC clone. (**I**) Proportions of cell types in E9.5 embryos derived from the mESC clones in (H). (**J**) UMAP embedding from (E) colored by developmental domain. Representative embryos derived from injection of activated PEtracer cells into tetraploidized host morulae at (**K**) E9.5, (**L**) E12.5, and (**M**) E15.5.

**Figure S2.**
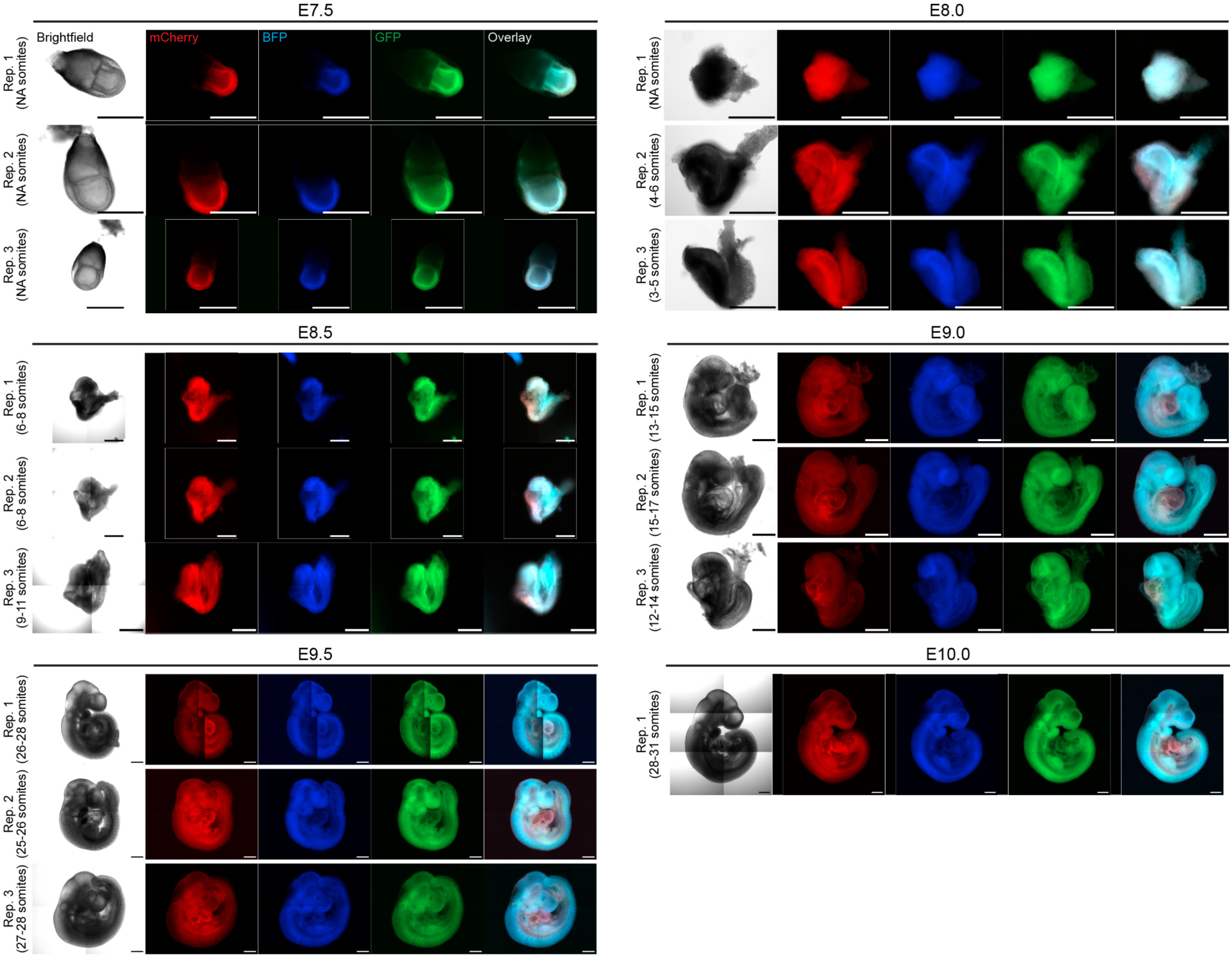
Fluorescent characterization of PEtracer embryos collected across E7.5-E10.0, related to Figure 1. Representative imaging of PEtracer embryos spanning E7.5 to E10.0 prior to dissociation and sequencing. For each embryo, four channels are shown: brightfield, PEmax-T2A-GFP (green), mCherry-linked LTCs (red), and BFP-linked pegArray (blue), along with an overlay of all fluorescent signals. Embryos are displayed in chronological order of developmental stage to illustrate progressive changes in morphology and reporter expression across gastrulation and early organogenesis. Biological triplicates were collected for each developmental stage except E10.0. All scale bars = 500 μm. Somite counts are indicated for each embryo where determinable (NA, not assessable).

**Figure S3.**
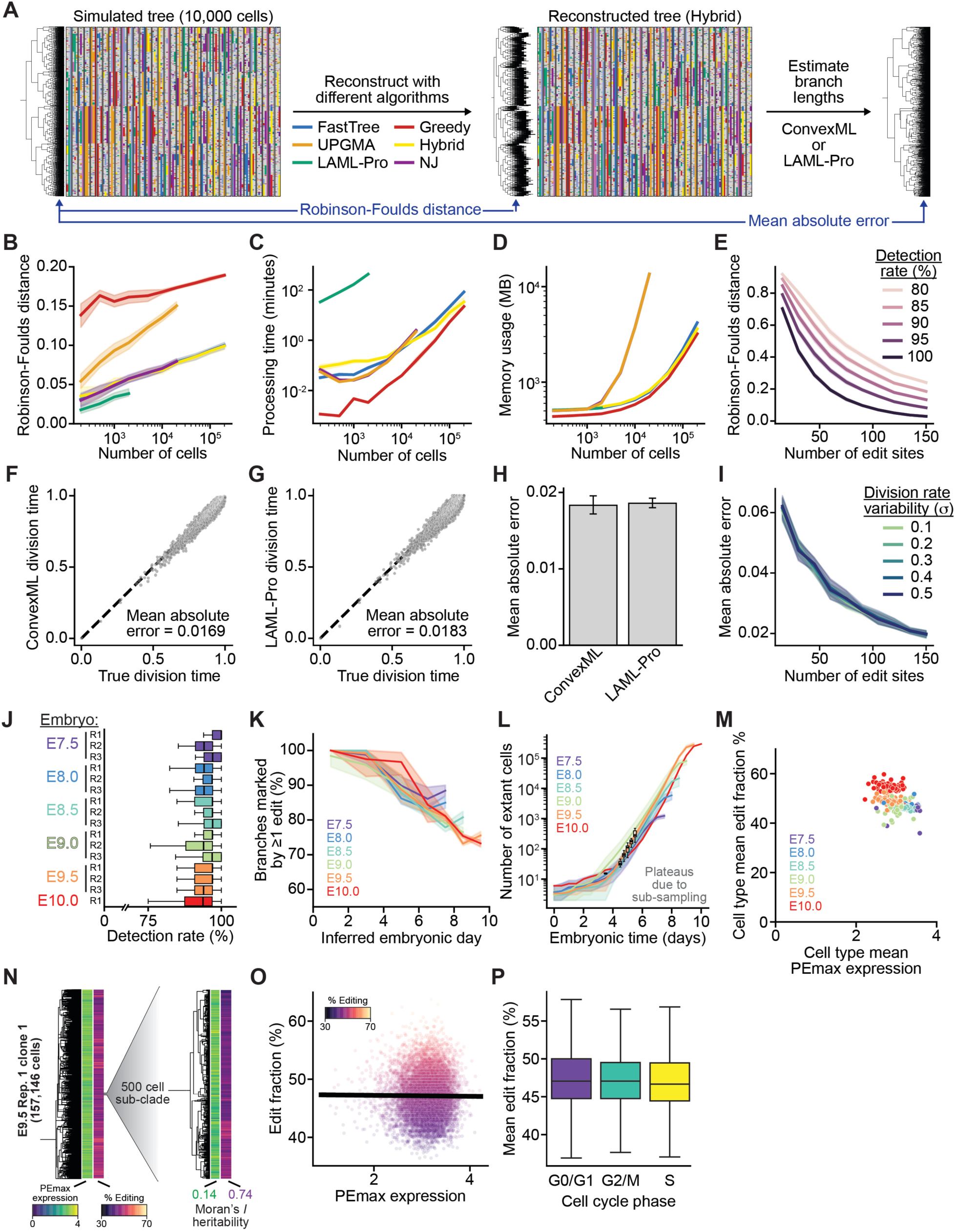
Benchmarking lineage tree reconstruction on simulated data and validation in PEtracer embryos, related to Figure 1. (**A**) Schematic of the simulation-based benchmarking framework. Ground-truth lineage trees and tracing data were simulated (**Methods**), then reconstructed using six algorithms: FastTree (blue)^1^, UPGMA (orange), LAML-Pro (green)^2^, Greedy (red)^3^, Hybrid (yellow), and Neighbor-Joining (NJ; purple). Reconstruction accuracy was quantified as the normalized Robinson-Foulds (RF) distance between the simulated and reconstructed topologies. Branch lengths were subsequently estimated using either ConvexML^4^ or LAML-Pro^2^, and division-timing accuracy was quantified as the mean absolute error (MAE) relative to ground truth. (**B**) Normalized RF distance between simulated and reconstructed trees as a function of the number of cells for each reconstruction algorithm. (**C**) Processing time (minutes) and (**D**) peak memory usage (MB) for each algorithm as a function of the number of cells. (**E**) Normalized RF distance for the Hybrid approach as a function of the number of edit sites, colored by LM detection rate. (**F, G**) Inferred versus ground truth division time for each internal node using (**F**) ConvexML (MAE = 0.0169) and (**G**) LAML-Pro (MAE = 0.0183); dashed line indicates y = x. (**H**) MAE of inferred division times for ConvexML and LAML-Pro across 10 replicates. (**I**) MAE of ConvexML-inferred division times as a function of the number of edit sites, colored by division rate variability (σ = 0.1 - 0.5). For (B–E) and (I), lines represent the mean and shaded regions the 95% confidence interval across 10 simulated replicates. (**J**) Lineage mark (LM) detection rate by scRNA-seq across all sampled embryos; R = replicate. (**K**) Mean fraction of branches marked by at least one edit across all sampled embryos, colored by timepoint. (**L**) Mean estimated number of extant cells per embryonic timepoint as a function of inferred time based on branch length estimates. Boxplots indicate published epiblast cell count estimates between E3.5 and E5.5^5^. (**M**) PEmax expression versus mean edit fraction of cell types colored by embryonic timepoint. (**N**) Representative cellular lineage tree for E9.5 R1 clone 1, comprising 157,146 cells with a 500 cell sub-clade also shown. Color bars indicate PEmax expression (left) and percent of edit sites marked with an LM (% editing, right), for each cell. Moran’s *I* heritability provided for both PEmax expression (green) and edit fraction (purple). Edit fraction versus (**O**) PEmax expression and (**P**) by cell cycle phase for all cells in E9.5 R1 clone 1.

**Figure S4.**
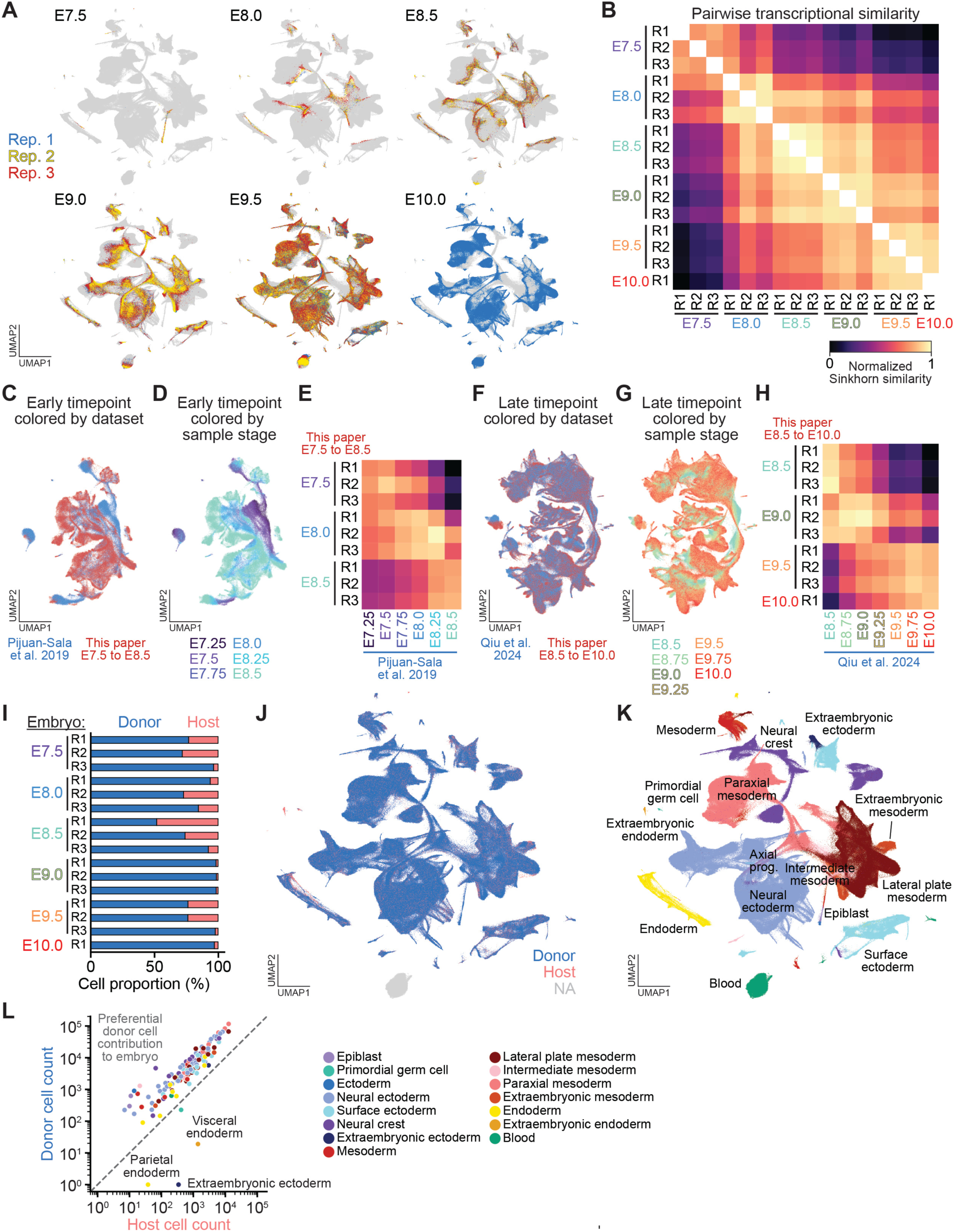
Transcriptional staging and chimerism quantification of PEtracer embryos, related to Figure 2. (**A**) UMAP embedding of all 1,792,520 cells from 16 embryos spanning E7.5 to E10.0 (see Figure 2A). Cells from each of three biological replicates for the indicated timepoint are colored separately (Rep. 1, blue; Rep. 2, yellow; Rep. 3, red), with cells from the other timepoints shown in gray. (**B**) Heatmap of pairwise transcriptional similarity between all replicates across timepoints, quantified by normalized Sinkhorn similarity. High within-stage similarity and graded between-stage similarity reflect the continuous progression of embryonic development. (**C-E**) Integration of the early timepoints from this study (E7.5-E8.5) with the reference dataset from Pijuan-Sala et al., 2019^6^. (**C**) Joint UMAP colored by dataset of origin (Pijuan-Sala et al., blue; this paper, red). (**D**) The same UMAP colored by developmental timepoint. (**E**) Pairwise Sinkhorn similarity heatmap comparing replicates from this study (rows) to timepoints from Pijuan-Sala et al. (columns). (**F-H**) Analogous integration of the late timepoints from this study (E8.5-E10.0) with the reference dataset from Qiu et al., 2024^7^. (**F**) Joint UMAP colored by dataset. (**G**) The same UMAP colored by developmental timepoint. (**H**) Pairwise Sinkhorn similarity heatmap comparing replicates from this study (rows) to timepoints from Qiu et al. (columns). (**I**) Proportion of tracing donor and non-tracing host cells across all 16 embryos (mean 85%, range 52-99%). (**J**) UMAP of all cells colored by donor (blue) versus host (salmon) cell identity, excluding primitive erythrocytes. (**K**) UMAP colored by developmental domain. (**L**) Relative contributions of donor and host cells to each developmental domain across all embryos.

**Figure S5.**
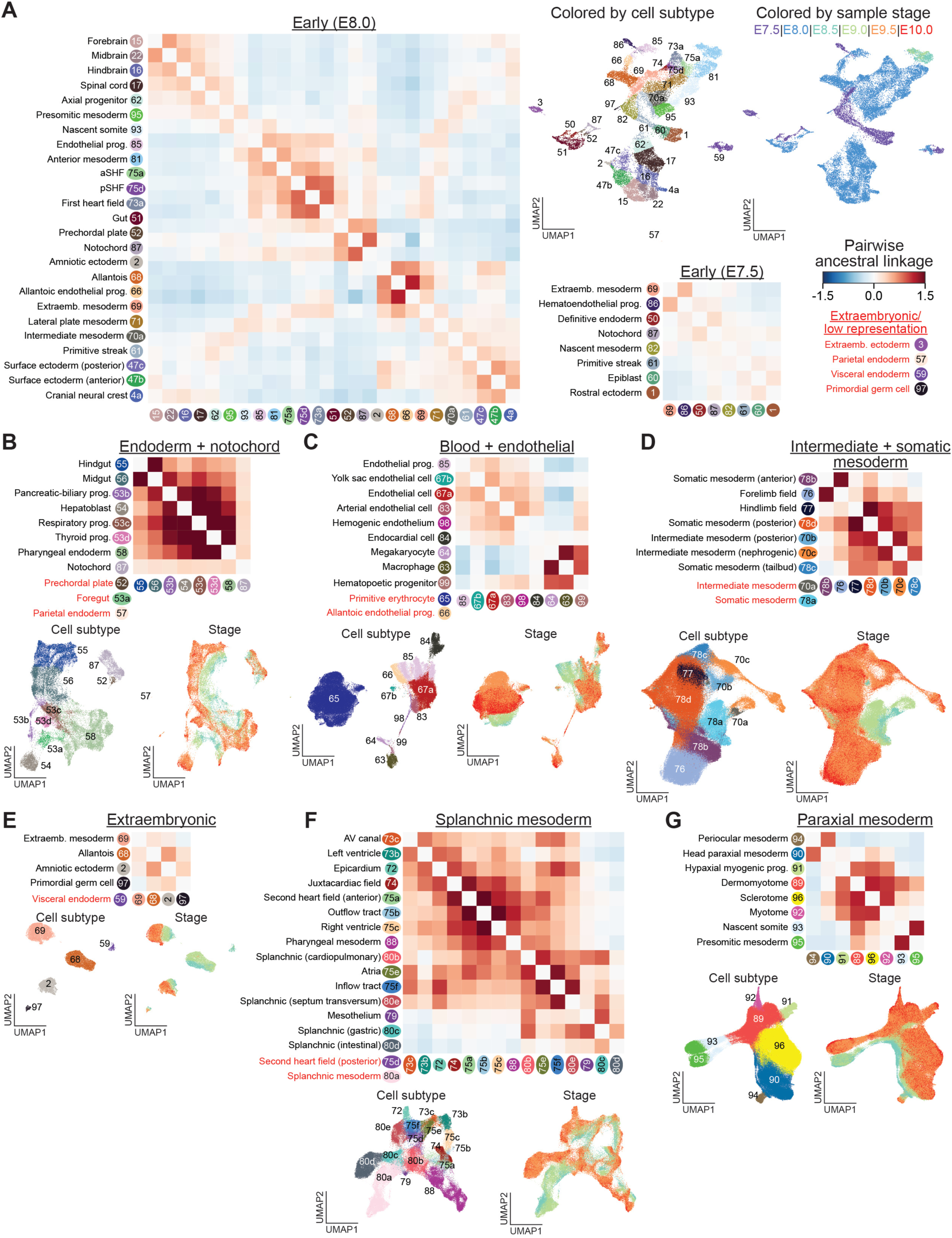
Cell subtype ancestral linkage and embryonic staging, related to Figure 2. Ancestral linkage heatmaps with cell subtype labels and UMAP embeddings colored by cell subtype and sample stage for the following developmental domains: (**A**) Early embryonic timepoints (E7.5 and E8.0), (**B**) endoderm and notochord, (**C**) blood and endothelial, (**D**) intermediate and somatic mesoderm, (**E**) extraembryonic, (**F**) splanchnic mesoderm, and (**G**) paraxial mesoderm. Cell subtypes included in UMAPs but excluded from the ancestral linkage heatmaps due to low representation are colored in red.

**Figure S6.**
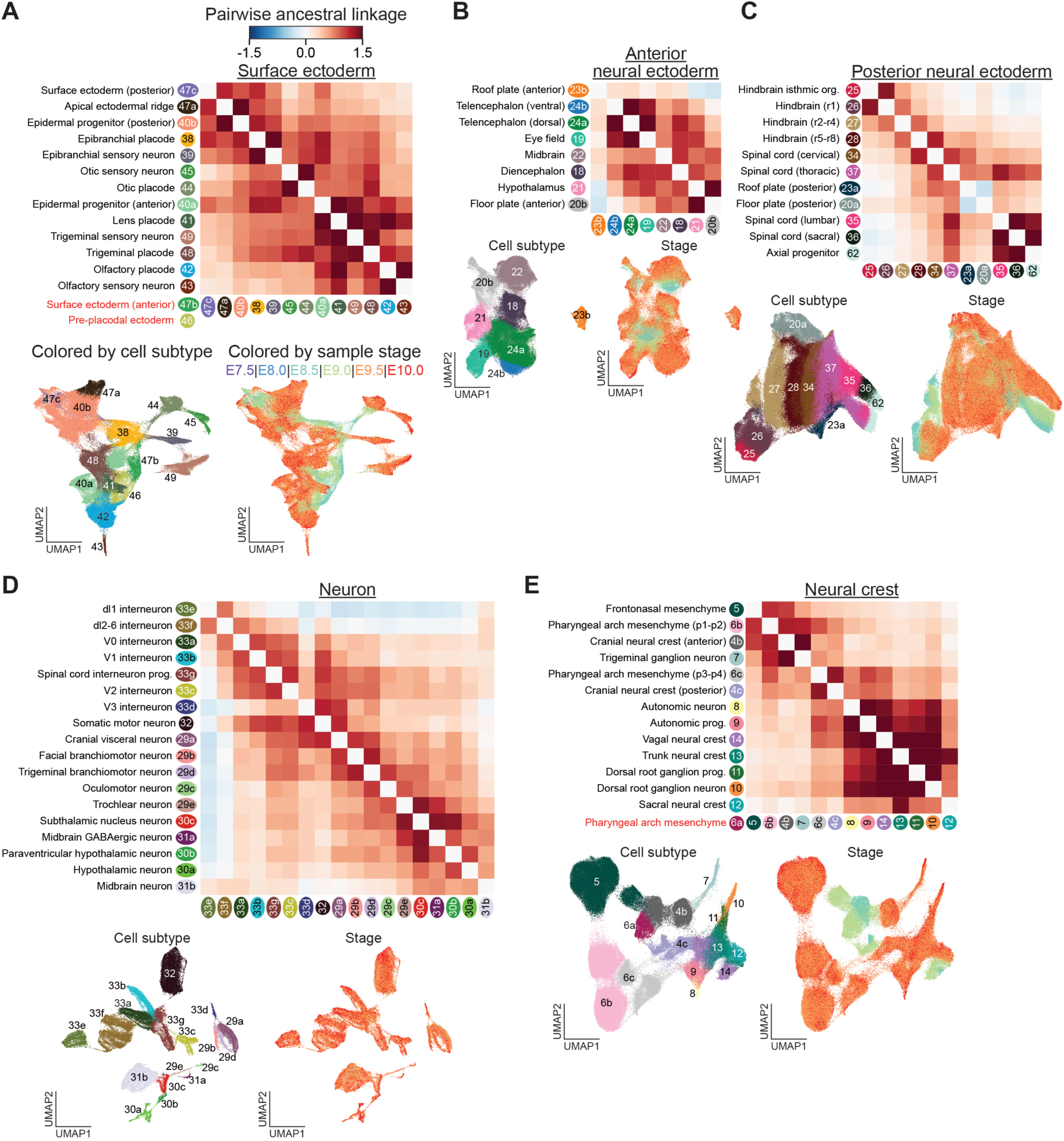
Cell subtype ancestral linkage and embryonic staging, related to Figure 2. Ancestral linkage heatmaps with cell subtype labels and UMAP embeddings colored by cell subtype and sample stage for the following developmental domains: (**A**) surface ectoderm, (**B**) anterior neural ectoderm, (**C**) posterior neural ectoderm, (**D**) neurons, and (**E**) neural crest. Cell subtypes included in UMAPs but excluded from the ancestral linkage heatmaps due to low representation are colored in red.

**Figure S7.**
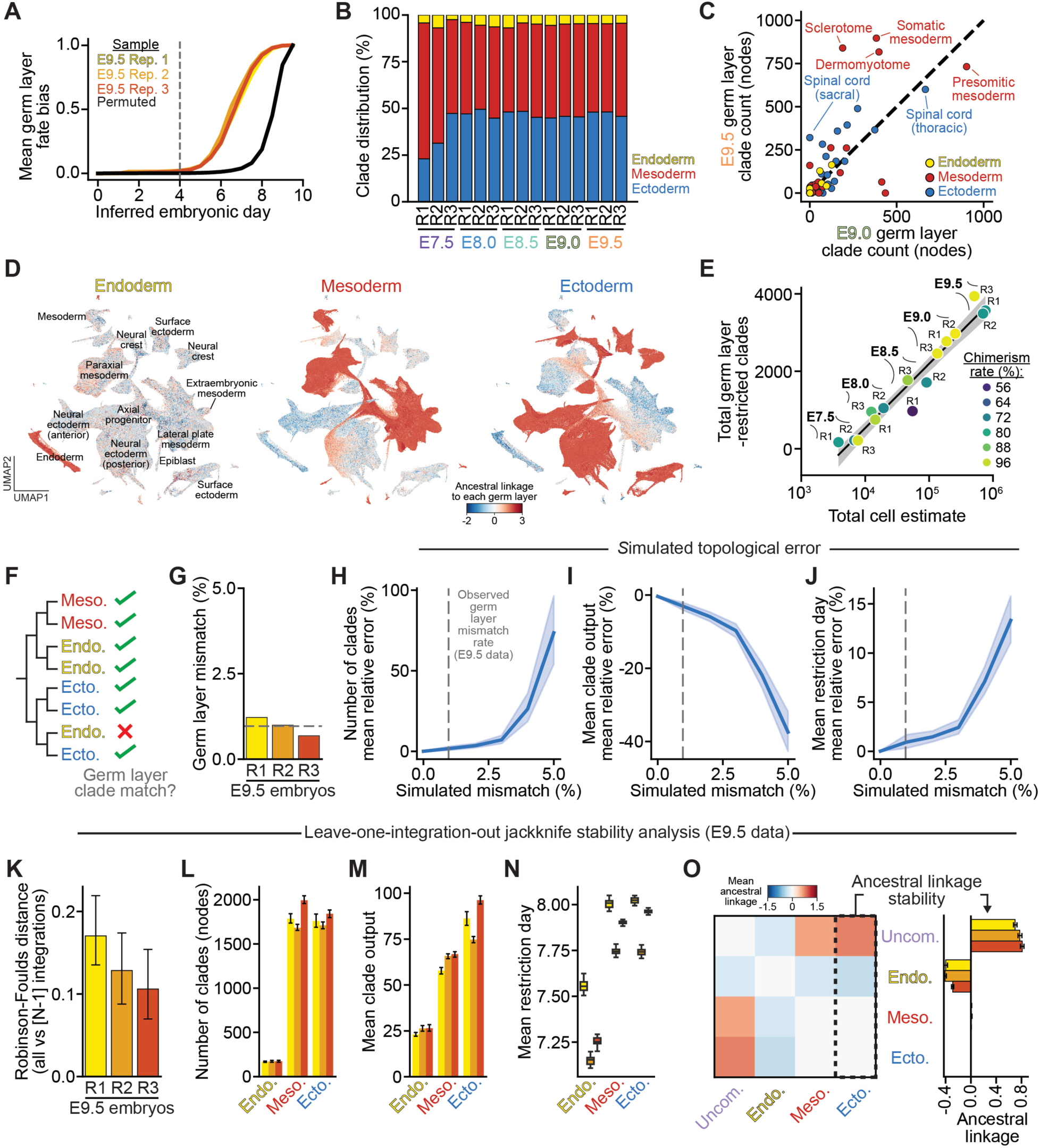
Quantification of germ layer specification dynamics with stability analysis, related to Figure 3. (**A**) Mean germ layer-level fate bias over inferred embryonic day for three E9.5 replicates and a permuted control (black). The dashed line marks the earliest time at which fate bias is statistically significant (permutation test) across all three replicates. (**B**) Distribution of germ layer-restricted clades for each replicate (R1–R3) at each sampled timepoint (E7.5–E9.5), shown as the percentage of clades assigned to endoderm (yellow), mesoderm (red), or ectoderm (blue). (**C**) Number of germ layer-restricted clades contributing to each cell type at E9.0 vs. E9.5. Cell types are colored by germ layer assignment. (**D**) UMAP of all lineage traced cells colored by ancestral linkage to each germ layer (endoderm, mesoderm, ectoderm). Red indicates linkages that are closer than would be expected by chance and blue indicates the opposite. (**E**) Total number of germ layer-restricted clades versus estimated total cell number for each embryo, labeled by timepoint (E7.5–E9.5) and replicate (R1–R3) and colored by chimerism rate. Line and shading indicate the linear fit and 95% CI. (**F**) Schematic of the percent germ layer mismatch metric: the germ layer label of each leaf is compared to the most parsimonious (Fitch-Hartigan) label for its parent. (**G**) Observed germ layer mismatch rate for E9.5 replicates R1-R3; dashed line indicates the mean (0.97% of cells). (**H-J**) Mean relative error in (**H**) the number of fate-restricted clades, (**I**) mean clade output, and (**J**) mean germ layer restriction day as a function of simulated topological error (germ layer mismatch rate, 0-5%). Line and shading indicate mean and 95% CI; vertical dashed line marks the observed E9.5 mismatch rate (**G**). (**K-O**) Leave-one-integration-out jackknife analysis (E9.5 data): trees were reconstructed after removing each intBC in turn (N-1 integrations) and compared to the full-data tree. (**K**) Normalized Robinson-Foulds (RF) distance between full and N-1 jackknife trees for R1-R3; mean ± 95% CI. (**L**) Number of fate-restricted clades, (**M**) mean clade output, and (**N**) mean restriction day for endoderm (Endo.), mesoderm (Meso.), and ectoderm (Ecto.), colored by replicate (R1-R3). (**O**) Left, mean pairwise ancestral linkage between uncommitted progenitors (Uncom.), Endo., Meso., and Ecto. across jackknife trees. Right, linkage of each population to ectoderm (boxed column) across jackknife trees (mean ± 95% CI), showing stability to integration dropout.

**Figure S8.**
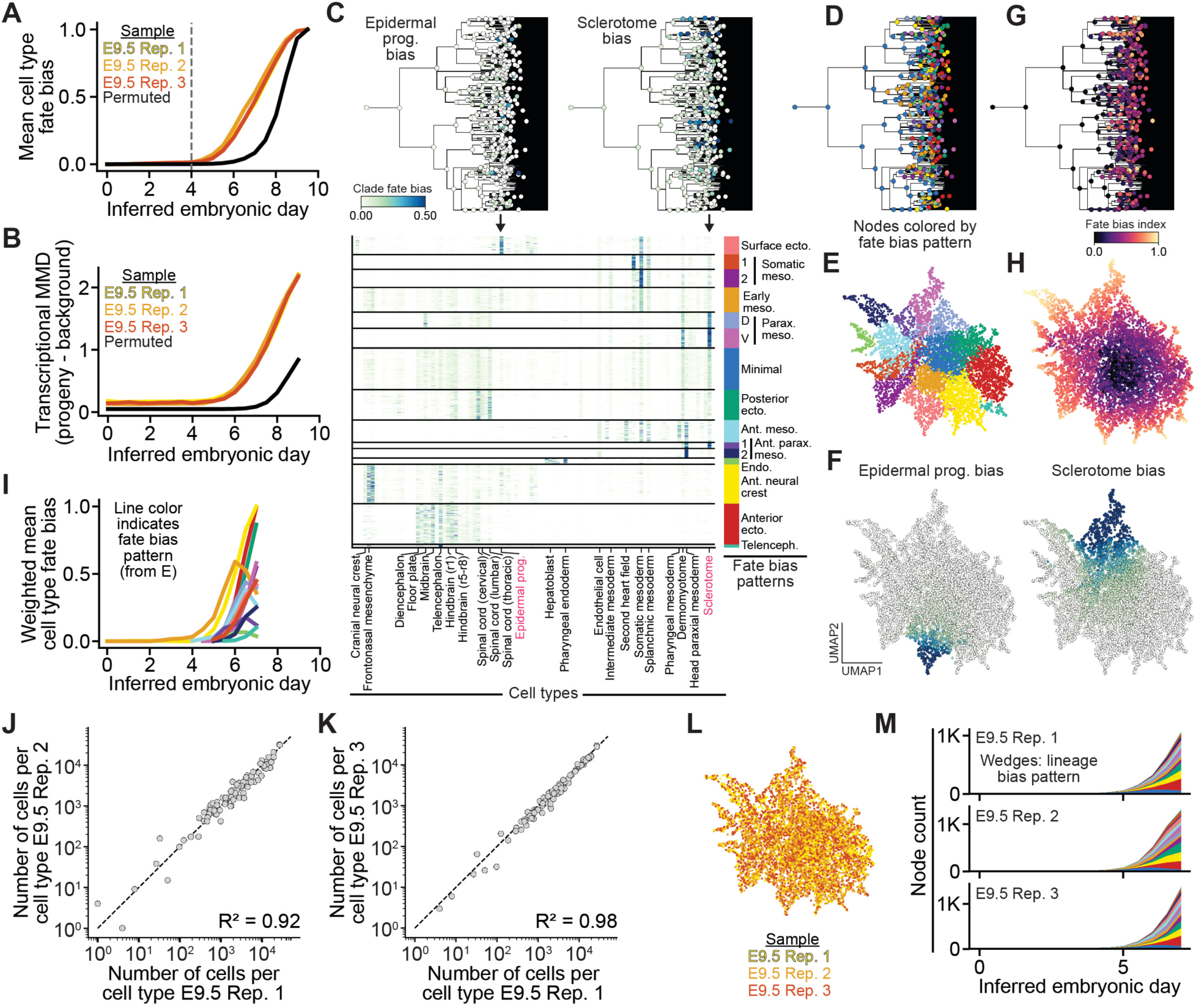
Quantification of fate bias and reproducibility across cell types, related to Figure 3. (**A**) Mean cell type-level fate bias over inferred embryonic day for three E9.5 replicates and a permuted control (black). The dashed line marks the earliest time at which fate bias is statistically significant (permutation test) across all three replicates. (**B**) Mean transcriptional MMD (maximum mean discrepancy) between descendants of internal nodes and the overall distribution, for three E9.5 replicates and a permuted control (black). (**C**) <u>Top:</u> Example lineage tree (E9.5-R1-C2) with early ancestral nodes (inferred day <7 and >50 descendants) from E9.5 replicates colored by fate bias toward epidermal progenitor (left) and sclerotome (right) identities. <u>Bottom:</u> Heatmap representing early ancestral node fate bias (rows) across cell types (columns), grouped by dominant fate bias pattern (colored labels, right). (**D**) Example lineage tree with ancestral nodes colored by assigned fate bias pattern. (**E**) UMAP embedding of ancestral node fate distributions colored by assigned fate bias pattern. (**F**) UMAP embedding from (E) colored by fate bias towards epidermal progenitor (left) and sclerotome (right) identities as in (C). (**G,H**) Example tree and UMAP from (D,E) colored by cell type-level fate bias index. (**I**) Mean cell type-level fate bias for each lineage pattern weighted by abundance, with lines colored by fate bias pattern (from C). (**J, K**) Cell type abundances between E9.5 replicate 1 and replicate 2 (J, R²= 0.92) or replicate 3 (K, R²= 0.98). Each point represents a cell type. (**L**) UMAP embedding of ancestral nodes from (E) colored by E9.5 replicate. (**M**) Number of extant ancestral nodes over inferred embryonic day for each E9.5 replicate colored by fate bias patterns (from C).

**Figure S9.**
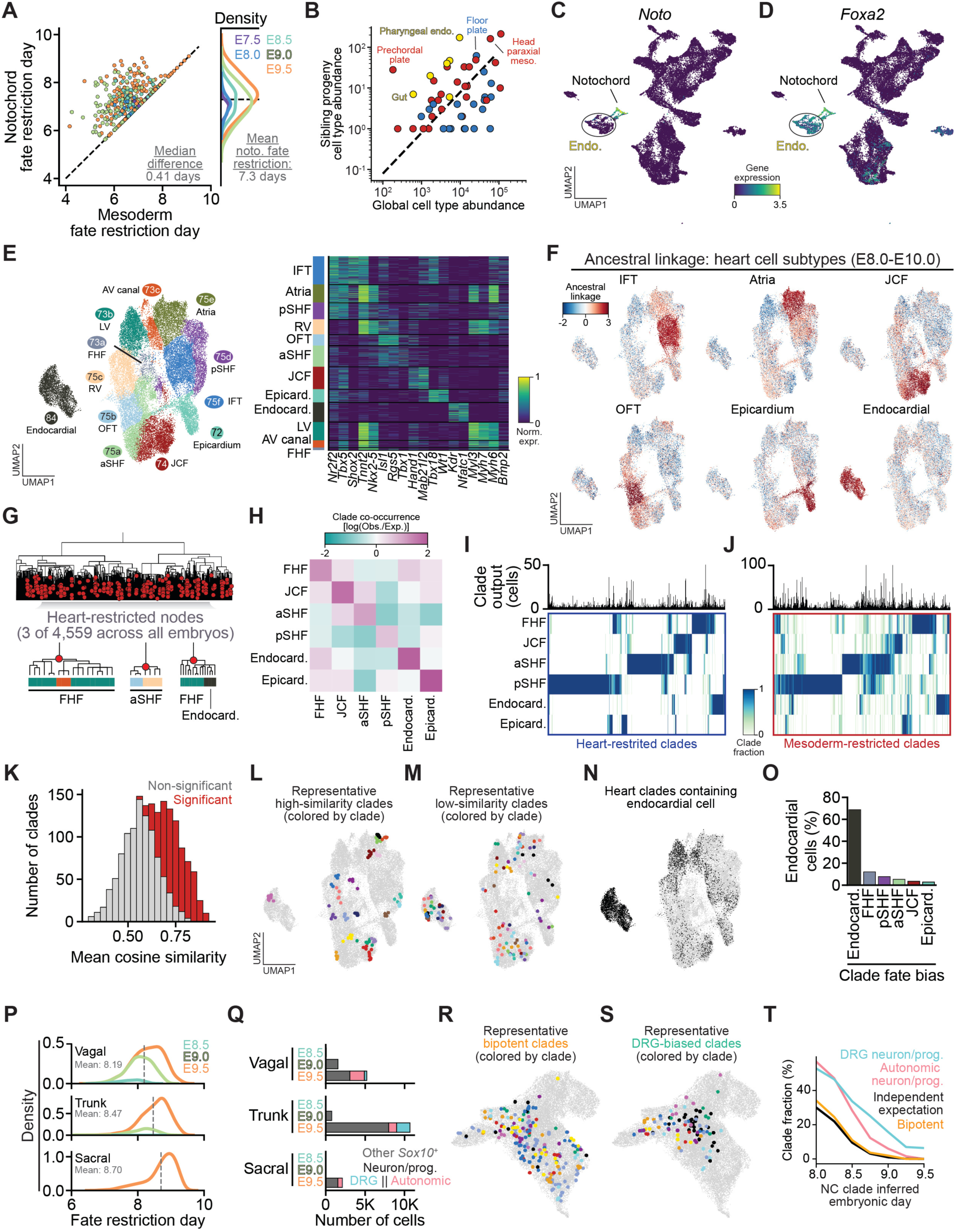
Supporting analyses for notochord and heart field fate restriction dynamics, related to Figure 4. (**A-D**) <u>Notochord:</u> (**A**) <u>Left:</u> Scatter plot of mesoderm fate restriction day versus notochord fate restriction day for each notochord-restricted clade. Points are colored by each clade’s sampled timepoint and points on the diagonal indicate concurrent restriction; the median difference between mesoderm and notochord restriction is 0.41 days. <u>Right:</u> Distribution of inferred notochord fate restriction times across triplicate sampled timepoints (E7.5–E9.5), shown as a kernel density estimate over clades in which the y-axis is probability density (area under curve = 1). Dashed line indicates the mean at inferred embryonic day 7.3. Note that we do not distinguish the Node from Notochord due to their high transcriptional similarity and differential timing. (**B**) Sibling clade cell type abundance versus global cell type abundance. Points above the diagonal are enriched among notochord sibling clades; selected cell types are labeled. Colors indicate germ layer. (**C**) UMAP of E7.5 and E8.0 cells colored by *Noto* expression, and (**D**) *Foxa2* expression. Notochord and endoderm populations are indicated. (**E-O**) <u>Heart fields:</u> (**E**) <u>Left:</u> UMAP of 12 annotated cardiac cell subtypes (as in Figure 4F). <u>Right:</u> heatmap of normalized expression of marker genes used for cardiac cell type annotation. IFT = inflow tract, pSHF = posterior second heart field, RV = right ventricle, OFT = outflow tract, aSHF = anterior SHF, JCF = juxtacardiac field, Epicard. = epicardial cells, Endocard. = endocardium, LV = left ventricle, AV canal = atrioventricular canal, FHF = first heart field. (**F)** Cardiac UMAPs colored by ancestral linkage to each cardiac cell type across E8.0-E10.0 embryos. (**G**) Example lineage tree (E9.5-R2-C1) with heart-restricted nodes (red circles) marked. Inset trees depict 3 of 4,559 heart-restricted nodes across all embryos. Color bars indicate cell type identity of descendant cells. (**H**) Heart field co-occurrence [log(observed/expected)] within heart-restricted clades across sampled timepoints (E8.0-E9.5). (**I**) Clustered heatmap of heart field (rows) representation for individual heart-restricted clades (columns). Clade output (number of cells) shown above. (**J**) Same as (I) for individual mesoderm-restricted clades (columns). (**K**) Histogram of mean cosine similarity of scVI embedding of heart cells within heart-restricted clades across E8.0-E10.0 embryos. Clades with statistically significant within-clade similarity are highlighted in red (permutation test). (**L**) Representative high-similarity and (**M**) low-similarity heart-restricted clades on the UMAP from (E). 10 colors indicate 10 randomly sampled clades. (**N**) Heart-restricted clades containing at least one endocardial cell (black) on the UMAP. (**O**) Distribution of fate bias assignments for endocardial cell-generating heart-restricted clades. (**P**) Distribution of inferred vagal, trunk, and sacral NC fate restriction timing across sampled timepoints (E8.5-E9.5). Dashed line indicates the mean. (**Q**) Cell type composition of vagal, trunk, and sacral NC clades across sampled timepoints, colored by DRG neuron/progenitor (teal), autonomic neuron/progenitor (pink), and other *Sox10*^+^ cells (gray). UMAP of posterior NC cells colored by (**R**) representative bipotent and (**S**) DRG-biased trunk NC clades. 10 colors indicate 10 randomly sampled clades. (**T**) Probability of an E9.5 clade containing DRG, autonomic, and bipotent outputs as a function of NC-restricted clade depth (inferred embryonic day). Black line indicates the expected bipotent frequency under statistical independence between DRG and autonomic fate occurrence.

**Figure S10.**
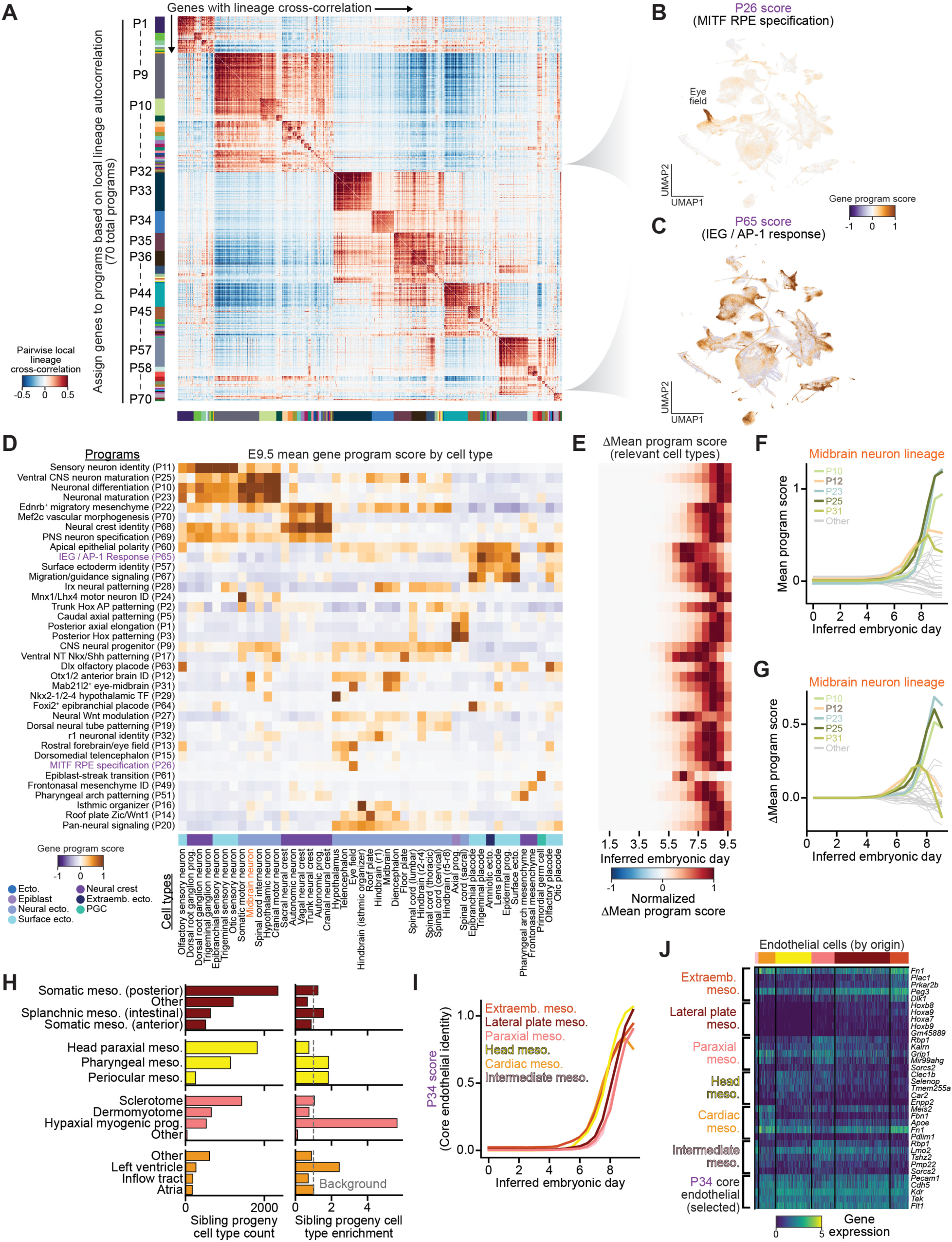
Gene regulatory program nomination and activity across ectodermal cell types, related to Figure 5. (**A**) Pairwise local lineage cross-correlation matrix for 1,322 genes assigned to programs across all embryos. Genes are ordered by their hierarchical clustering assignment via Hotspot, revealing blocks of coordinated structure. Colored bars indicate assignment of genes to individual gene programs (P1-P70). (**B**) UMAP of all sampled cells colored by P26 (MITF RPE specification) or (**C**) P65 (IEG/AP-1 response) program scores. Cell type-specific activation of P26 in the eye field is annotated in (B). For a full list of gene regulatory program names, abbreviations, and descriptions of their assignment see **Table S5**. (**D**) Mean score for each program (rows) across ectodermal cell types (columns) in E9.5 embryos. Cell type developmental domain assignments are indicated by colored bars along the bottom. Highlighted programs in purple: P65 (IEG/AP-1 response) and P26 (MITF RPE specification) from (B,C). Highlighted cell type in orange is midbrain neuron. (**E**) Change in mean program score (ΔMean program score) over inferred embryonic day in E9.5 lineage trees for relevant cell types (mean >0.5 in at least one timepoint), normalized per program. (**F**) Mean program score and (**G**) change in mean program score over inferred embryonic day for gene programs within the midbrain neuron lineage. Programs include P10 (neuronal differentiation), P12 (Otx1/2 anterior brain ID), P23 (neuronal maturation), P25 (ventral CNS neuron maturation), and P31 (Mab21l2⁺ eye-midbrain). (**H**) Endothelial-restricted clade sibling progeny counts (left) and fold enrichment relative to background (right) for E9.5 replicates 1-3. (**I**) P34 (core endothelial identity) mean program score over inferred embryonic day in E9.5 lineage trees colored by inferred endothelial origin. (**J**) Heatmap of E9.5 endothelial cells grouped by inferred origin showing the expression of top 5 differentially expressed genes for each origin as well as selected genes from core endothelial program.

**Figure S11.**
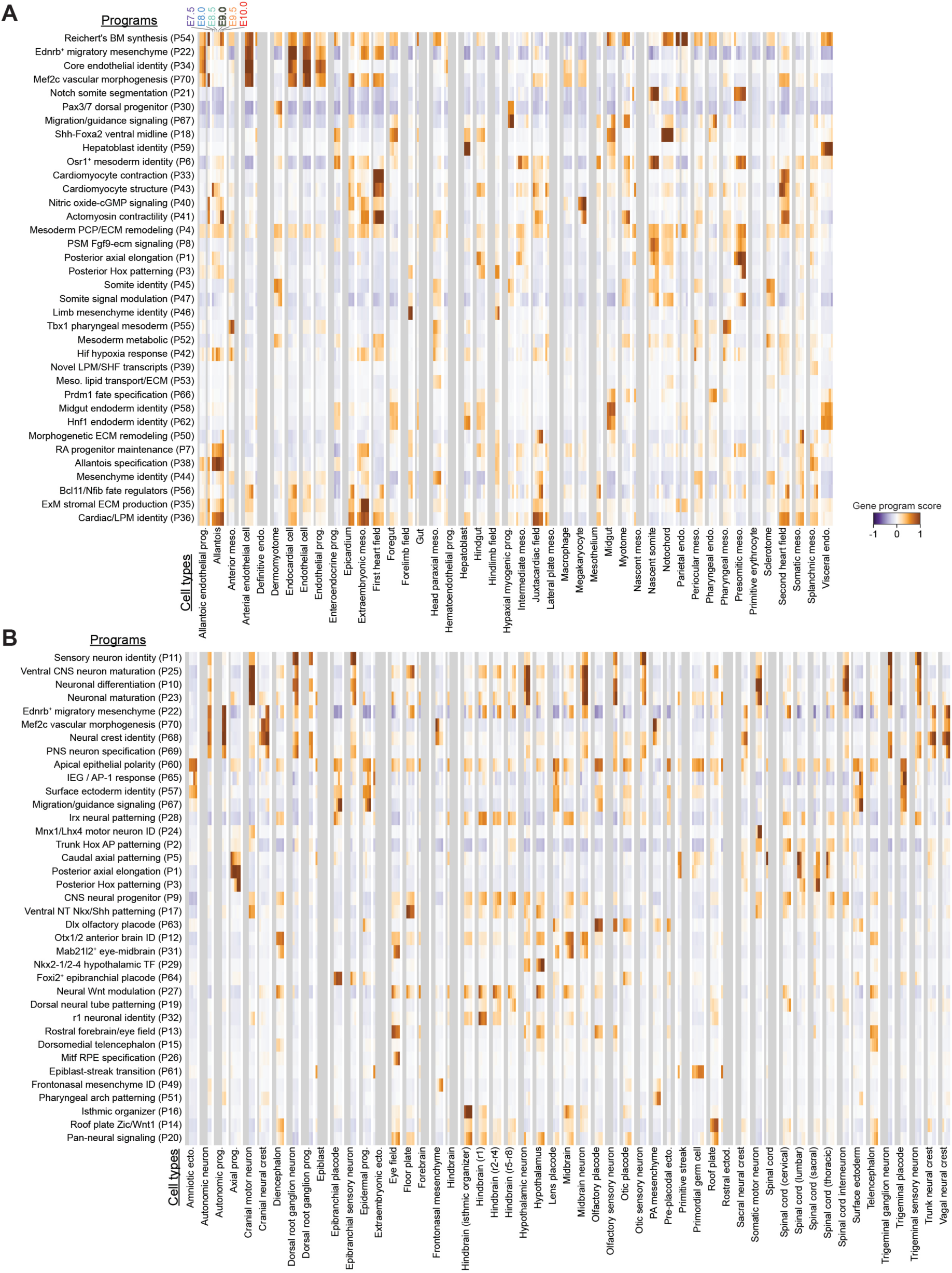
Gene program activity across cell types over time, related to Figure 5. Gene program activity (rows) across sampled timepoints from E7.5 to E10.0 for (**A**) mesodermal and endodermal as well as (**B**) ectodermal cell types. Timepoints where cell types are not present are shown in grey.

**Figure S12.**
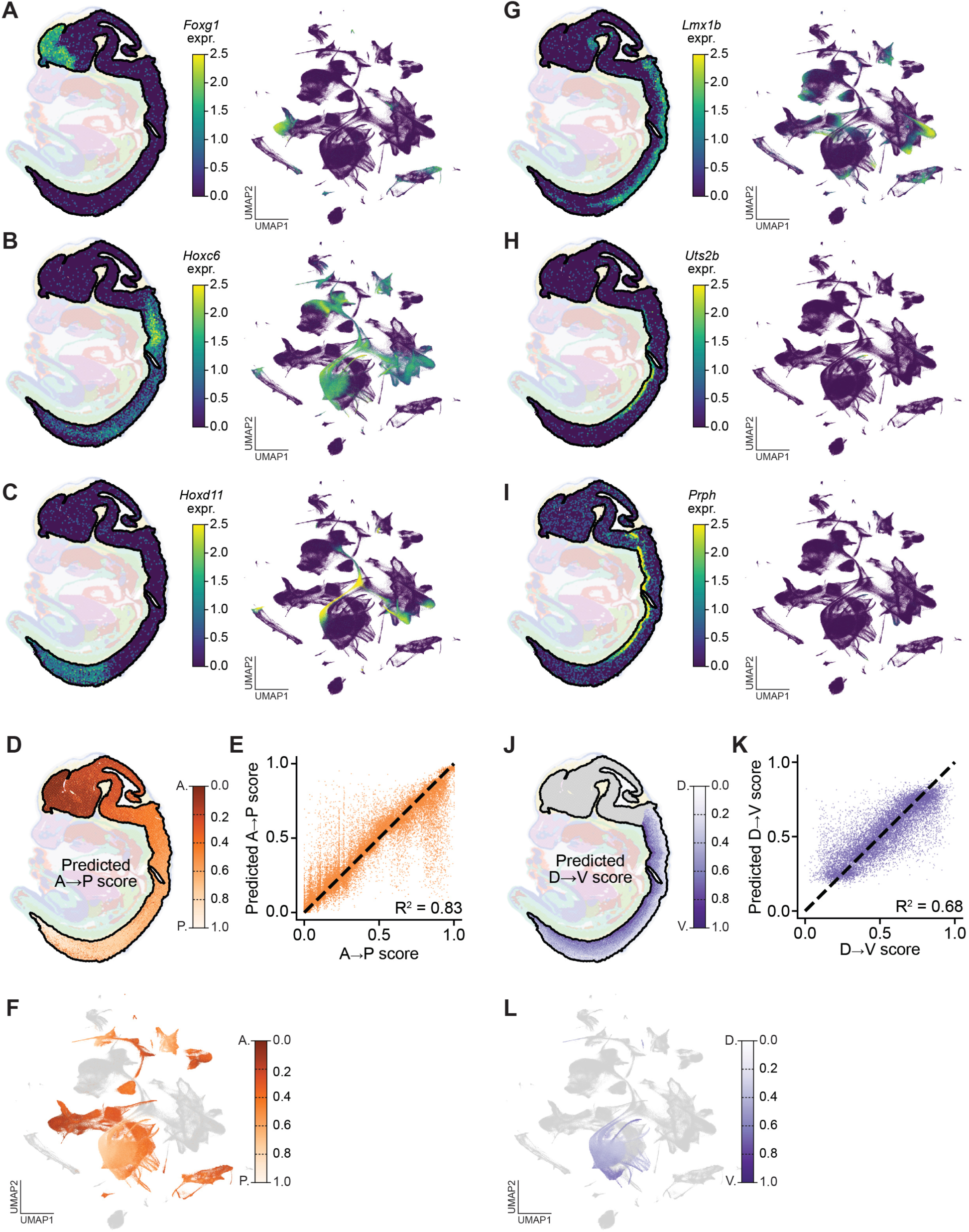
Spatial transcriptomics-based inference of spatial axes, related to Figure 6. (**A-C**) <u>Left:</u> spatial transcriptomics data from a sagittal section of a E13.5 mouse embryo (Stereo-seq; Chen et al., 2022). Neural ectoderm is outlined and colored by expression of example anterior-posterior (A-P) patterning genes, while other tissues are shown in the background for reference. <u>Right:</u> UMAP embedding of all cells in our dataset colored by the expression of the same genes: (**A**) *Foxg1*, (**B**) *Hoxc6*, and (**C**) *Hoxd11*. (**D**) Predicted A-P score of stereo-seq spots, using our model (5-fold cross validation). (**E**) Concordance of predicted and inferred A-P position across the stereo-seq dataset (R²= 0.83). (**F**) UMAP embedding of all cells in our dataset colored by predicted A-P score. (**G-I**) Same as (A-C) for dorsal-ventral (D-V) patterning genes: (**G**) *Lmx1b*, (**H**) *Uts2b*, and (**I**) *Prph*. (**J,K**) Same as (D,E) for D-V score. (**J**) Predicted D-V score of stereo-seq spots, using our model (5-fold cross validation) and (**K**) Concordance of predicted and inferred D-V position across the stereo-seq dataset (R²= 0.68). (**L**) UMAP embedding of all cells in our dataset colored by predicted D-V score.

**Figure S13.**
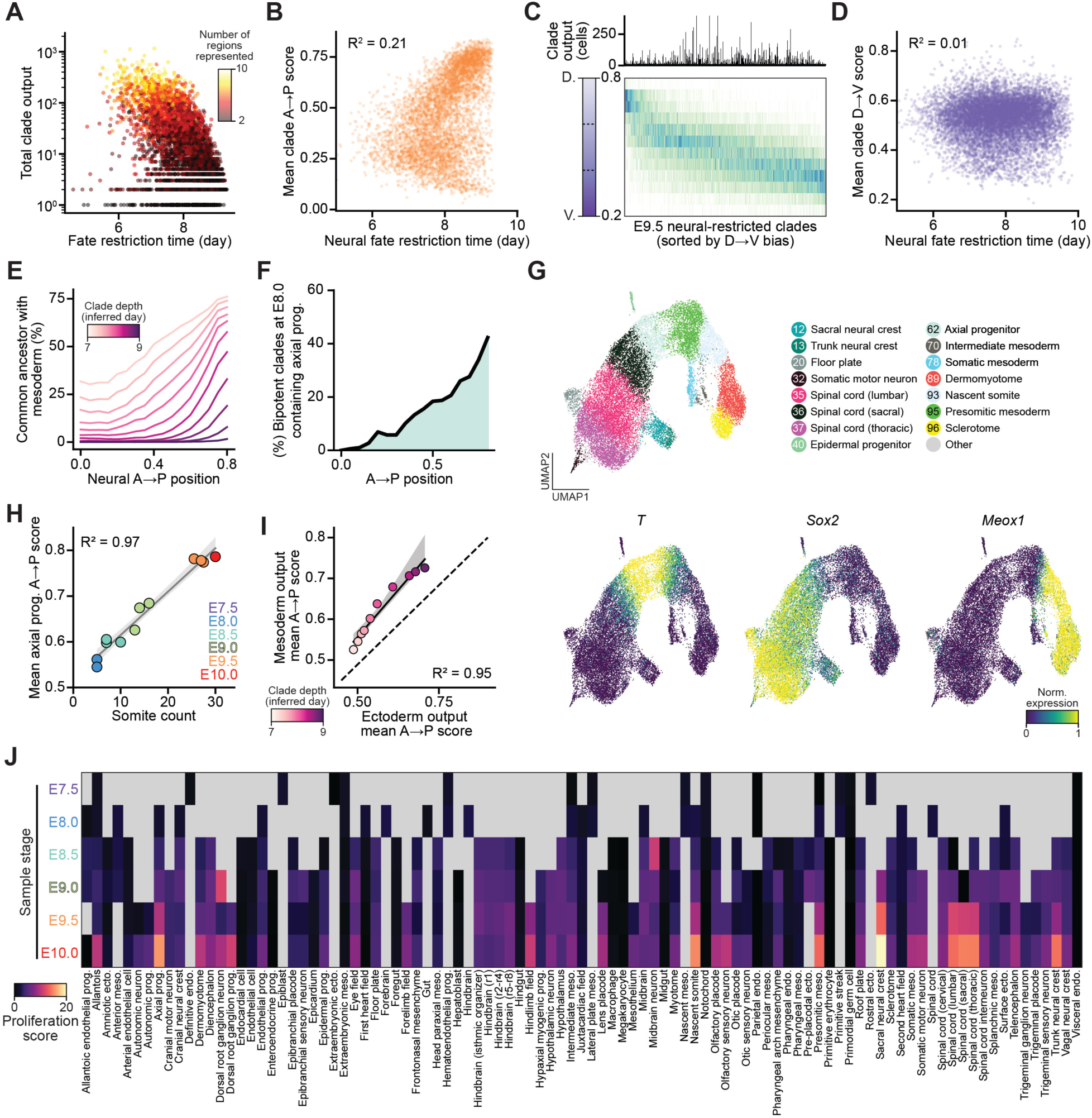
Supporting analysis for spatial axes, regionalization, and axis elongation, related to Figure 7. (**A**) Total clade output (number of descendant cells) versus neural fate restriction time. Points are colored by the number of neural regions represented in each clade. (**B**) Mean A-P score of neural-restricted clades versus fate restriction time. (**C**) Heatmap showing the distribution of D-V positions (rows) for spinal cord clades in E9.5 embryos with >10 descendants (columns), ordered by mean D-V score. Clade output (number of cells) shown above. (**D**) Same as (B) but for mean D-V score of spinal cord clades. (**E**) Fraction of neural cells by A-P position sharing a common ancestor with the mesoderm at clade depths from E7.0-E9.0 (inferred embryonic day). (**F**) Fraction of bipotent clades with common ancestor at E8.0 that contain observed axial progenitors at E9.5, plotted across the A-P axis. (**G**) <u>Top:</u> UMAP of E9.5 posterior neural-restricted clades and their siblings colored by cell type (numbering as in Figure 2) and <u>Bottom:</u> *T*, *Sox2*, and *Meox1* expression. (**H**) Mean axial progenitor A-P score versus somite number across developmental timepoints. Trend line indicates linear regression fit (R²= 0.97). (**I**) Mean A-P scores of mesodermal versus ectodermal outputs from bipotent clades at depths from E7.0-E9.0 (inferred embryonic day). Trend line indicates linear regression fit (R²= 0.95); dashed line indicates y = x. (**J**) Proliferation scores (calculated as mean number of cells sharing a common ancestor within the previous 24 hours) across cell types and developmental timepoints (E7.5-E10.0).

## References

1. Tarkowski, A.K. (1959). Experiments on the development of isolated blastomeres of mouse eggs. Nature 184, 1286–1287. 10.1038/1841286a0.

2. Wagner, D.E., and Klein, A.M. (2020). Lineage tracing meets single-cell omics: opportunities and challenges. Nat. Rev. Genet. 21, 410–427. 10.1038/s41576-020-0223-2.

3. Domcke, S., and Shendure, J. (2023). A reference cell tree will serve science better than a reference cell atlas. Cell 186, 1103–1114. 10.1016/j.cell.2023.02.016.

4. Sulston, J.E., Schierenberg, E., White, J.G., and Thomson, J.N. (1983). The embryonic cell lineage of the nematode Caenorhabditis elegans. Dev. Biol. 100, 64–119. 10.1016/0012-1606(83)90201-4.

5. Kwak, J., Park, O.K., Jung, Y.J., Hwang, B.J., Kwon, S.-H., and Kee, Y. (2013). Live image profiling of neural crest lineages in zebrafish transgenic lines. Mol. Cells 35, 255–260. 10.1007/s10059-013-0001-5.

6. Lange, M., Granados, A., VijayKumar, S., Bragantini, J., Ancheta, S., Kim, Y.-J., Santhosh, S., Borja, M., Kobayashi, H., McGeever, E., et al. (2024). A multimodal zebrafish developmental atlas reveals the state-transition dynamics of late-vertebrate pluripotent axial progenitors. Cell 187, 6742–6759.e17. 10.1016/j.cell.2024.09.047.

7. Olivier, N., Luengo-Oroz, M.A., Duloquin, L., Faure, E., Savy, T., Veilleux, I., Solinas, X., Débarre, D., Bourgine, P., Santos, A., et al. (2010). Cell lineage reconstruction of early zebrafish embryos using label-free nonlinear microscopy. Science 329, 967–971. 10.1126/science.1189428.

8. Wang, M., Zhang, Q., Wang, C., Chi, Y., Zheng, W., Mu, Z., Cao, X., Zhang, W., Yang, B., Schier, A.F., et al. (2026). High-Fidelity Long-term Whole-embryo Lineage and Fate Reconstruction by Iterative Tracking with Error Correction. BioRxiv Prepr. Serv. Biol., 2026.03.12.711203. 10.64898/2026.03.12.711203.

9. Metzger, R.J., Klein, O.D., Martin, G.R., and Krasnow, M.A. (2008). The branching programme of mouse lung development. Nature 453, 745–750. 10.1038/nature07005.

10. Lawson, K.A., Meneses, J.J., and Pedersen, R.A. (1991). Clonal analysis of epiblast fate during germ layer formation in the mouse embryo. Development 113, 891–911. 10.1242/dev.113.3.891.

11. Wilson, V., and Beddington, R.S. (1996). Cell fate and morphogenetic movement in the late mouse primitive streak. Mech. Dev. 55, 79–89. 10.1016/0925-4773(95)00493-9.

12. Jaenisch, R., and Mintz, B. (1974). Simian virus 40 DNA sequences in DNA of healthy adult mice derived from preimplantation blastocysts injected with viral DNA. Proc. Natl. Acad. Sci. U. S. A. 71, 1250–1254. 10.1073/pnas.71.4.1250.

13. Livet, J., Weissman, T.A., Kang, H., Draft, R.W., Lu, J., Bennis, R.A., Sanes, J.R., and Lichtman, J.W. (2007). Transgenic strategies for combinatorial expression of fluorescent proteins in the nervous system. Nature 450, 56–62. 10.1038/nature06293.

14. Snippert, H.J., van der Flier, L.G., Sato, T., van Es, J.H., van den Born, M., Kroon-Veenboer, C., Barker, N., Klein, A.M., van Rheenen, J., Simons, B.D., et al. (2010). Intestinal crypt homeostasis results from neutral competition between symmetrically dividing Lgr5 stem cells. Cell 143, 134–144. 10.1016/j.cell.2010.09.016.

15. Tzouanacou, E., Wegener, A., Wymeersch, F.J., Wilson, V., and Nicolas, J.-F. (2009). Redefining the progression of lineage segregations during mammalian embryogenesis by clonal analysis. Dev. Cell 17, 365–376. 10.1016/j.devcel.2009.08.002.

16. VanHorn, S., and Morris, S.A. (2021). Next-Generation Lineage Tracing and Fate Mapping to Interrogate Development. Dev. Cell 56, 7–21. 10.1016/j.devcel.2020.10.021.

17. Wang, Z., and Jaenisch, R. (2004). At most three ES cells contribute to the somatic lineages of chimeric mice and of mice produced by ES-tetraploid complementation. Dev. Biol. 275, 192–201. 10.1016/j.ydbio.2004.06.026.

18. Briggs, J.A., Weinreb, C., Wagner, D.E., Megason, S., Peshkin, L., Kirschner, M.W., and Klein, A.M. (2018). The dynamics of gene expression in vertebrate embryogenesis at single-cell resolution. Science 360, eaar5780. 10.1126/science.aar5780.

19. Cao, J., Spielmann, M., Qiu, X., Huang, X., Ibrahim, D.M., Hill, A.J., Zhang, F., Mundlos, S., Christiansen, L., Steemers, F.J., et al. (2019). The single-cell transcriptional landscape of mammalian organogenesis. Nature 566, 496–502. 10.1038/s41586-019-0969-x.

20. Farrell, J.A., Wang, Y., Riesenfeld, S.J., Shekhar, K., Regev, A., and Schier, A.F. (2018). Single-cell reconstruction of developmental trajectories during zebrafish embryogenesis. Science 360, eaar3131. 10.1126/science.aar3131.

21. Pijuan-Sala, B., Griffiths, J.A., Guibentif, C., Hiscock, T.W., Jawaid, W., Calero-Nieto, F.J., Mulas, C., Ibarra-Soria, X., Tyser, R.C.V., Ho, D.L.L., et al. (2019). A single-cell molecular map of mouse gastrulation and early organogenesis. Nature 566, 490–495. 10.1038/s41586-019-0933-9.

22. Qiu, C., Martin, B.K., Welsh, I.C., Daza, R.M., Le, T.-M., Huang, X., Nichols, E.K., Taylor, M.L., Fulton, O., O’Day, D.R., et al. (2024). A single-cell time-lapse of mouse prenatal development from gastrula to birth. Nature 626, 1084–1093. 10.1038/s41586-024-07069-w.

23. Qiu, C., Cao, J., Martin, B.K., Li, T., Welsh, I.C., Srivatsan, S., Huang, X., Calderon, D., Noble, W.S., Disteche, C.M., et al. (2022). Systematic reconstruction of cellular trajectories across mouse embryogenesis. Nat. Genet. 54, 328–341. 10.1038/s41588-022-01018-x.

24. Scialdone, A., Tanaka, Y., Jawaid, W., Moignard, V., Wilson, N.K., Macaulay, I.C., Marioni, J.C., and Göttgens, B. (2016). Resolving early mesoderm diversification through single-cell expression profiling. Nature 535, 289–293. 10.1038/nature18633.

25. Wan, Y., El Kholtei, J., Jenie, I., Colomer-Rosell, M., Liu, J., Zhang, Q., Navajas Acedo, J., Du, L.Y., Codina-Tobias, M., Wang, M., et al. (2026). Whole-embryo spatial transcriptomics at subcellular resolution from gastrulation to organogenesis. Science 391, eadt3439. 10.1126/science.adt3439.

26. Weinreb, C., Rodriguez-Fraticelli, A., Camargo, F.D., and Klein, A.M. (2020). Lineage tracing on transcriptional landscapes links state to fate during differentiation. Science 367, eaaw3381. 10.1126/science.aaw3381.

27. Alemany, A., Florescu, M., Baron, C.S., Peterson-Maduro, J., and van Oudenaarden, A. (2018). Whole-organism clone tracing using single-cell sequencing. Nature 556, 108–112. 10.1038/nature25969.

28. Chan, M.M., Smith, Z.D., Grosswendt, S., Kretzmer, H., Norman, T.M., Adamson, B., Jost, M., Quinn, J.J., Yang, D., Jones, M.G., et al. (2019). Molecular recording of mammalian embryogenesis. Nature 570, 77–82. 10.1038/s41586-019-1184-5.

29. Bowling, S., Sritharan, D., Osorio, F.G., Nguyen, M., Cheung, P., Rodriguez-Fraticelli, A., Patel, S., Yuan, W.-C., Fujiwara, Y., Li, B.E., et al. (2020). An Engineered CRISPR-Cas9 Mouse Line for Simultaneous Readout of Lineage Histories and Gene Expression Profiles in Single Cells. Cell 181, 1410–1422.e27. 10.1016/j.cell.2020.04.048.

30. Frieda, K.L., Linton, J.M., Hormoz, S., Choi, J., Chow, K.-H.K., Singer, Z.S., Budde, M.W., Elowitz, M.B., and Cai, L. (2017). Synthetic recording and in situ readout of lineage information in single cells. Nature 541, 107–111. 10.1038/nature20777.

31. He, Z., Maynard, A., Jain, A., Gerber, T., Petri, R., Lin, H.-C., Santel, M., Ly, K., Dupré, J.-S., Sidow, L., et al. (2022). Lineage recording in human cerebral organoids. Nat. Methods 19, 90–99. 10.1038/s41592-021-01344-8.

32. Kalhor, R., Kalhor, K., Mejia, L., Leeper, K., Graveline, A., Mali, P., and Church, G.M. (2018). Developmental barcoding of whole mouse via homing CRISPR. Science 361, eaat9804. 10.1126/science.aat9804.

33. Li, L., Bowling, S., McGeary, S.E., Yu, Q., Lemke, B., Alcedo, K., Jia, Y., Liu, X., Ferreira, M., Klein, A.M., et al. (2023). A mouse model with high clonal barcode diversity for joint lineage, transcriptomic, and epigenomic profiling in single cells. Cell 186, 5183–5199.e22. 10.1016/j.cell.2023.09.019.

34. Loveless, T.B., Grotts, J.H., Schechter, M.W., Forouzmand, E., Carlson, C.K., Agahi, B.S., Liang, G., Ficht, M., Liu, B., Xie, X., et al. (2021). Lineage tracing and analog recording in mammalian cells by single-site DNA writing. Nat. Chem. Biol. 17, 739–747. 10.1038/s41589-021-00769-8.

35. Loveless, T.B., Carlson, C.K., Dentzel Helmy, C.A., Hu, V.J., Ross, S.K., Demelo, M.C., Murtaza, A., Liang, G., Ficht, M., Singhai, A., et al. (2025). Open-ended molecular recording of sequential cellular events into DNA. Nat. Chem. Biol. 21, 512–521. 10.1038/s41589-024-01764-5.

36. McKenna, A., Findlay, G.M., Gagnon, J.A., Horwitz, M.S., Schier, A.F., and Shendure, J. (2016). Whole-organism lineage tracing by combinatorial and cumulative genome editing. Science 353, aaf7907. 10.1126/science.aaf7907.

37. Quinn, J.J., Jones, M.G., Okimoto, R.A., Nanjo, S., Chan, M.M., Yosef, N., Bivona, T.G., and Weissman, J.S. (2021). Single-cell lineages reveal the rates, routes, and drivers of metastasis in cancer xenografts. Science 371, eabc1944. 10.1126/science.abc1944.

38. Raj, B., Wagner, D.E., McKenna, A., Pandey, S., Klein, A.M., Shendure, J., Gagnon, J.A., and Schier, A.F. (2018). Simultaneous single-cell profiling of lineages and cell types in the vertebrate brain. Nat. Biotechnol. 36, 442–450. 10.1038/nbt.4103.

39. Spanjaard, B., Hu, B., Mitic, N., Olivares-Chauvet, P., Janjuha, S., Ninov, N., and Junker, J.P. (2018). Simultaneous lineage tracing and cell-type identification using CRISPR-Cas9-induced genetic scars. Nat. Biotechnol. 36, 469–473. 10.1038/nbt.4124.

40. Yang, D., Jones, M.G., Naranjo, S., Rideout, W.M., Min, K.H.J., Ho, R., Wu, W., Replogle, J.M., Page, J.L., Quinn, J.J., et al. (2022). Lineage tracing reveals the phylodynamics, plasticity, and paths of tumor evolution. Cell 185, 1905–1923.e25. 10.1016/j.cell.2022.04.015.

41. Winter, E., Emiliani, F., Cook, A., Abderrahim, A., and McKenna, A. (2026). BASELINE: a CRISPR base editing platform for mammalian-scale single-cell lineage tracing. Nucleic Acids Res. 54, gkag769. 10.1093/nar/gkag769.

42. Choi, J., Chen, W., Minkina, A., Chardon, F.M., Suiter, C.C., Regalado, S.G., Domcke, S., Hamazaki, N., Lee, C., Martin, B., et al. (2022). A time-resolved, multi-symbol molecular recorder via sequential genome editing. Nature 608, 98–107. 10.1038/s41586-022-04922-8.

43. Koblan, L.W., Yost, K.E., Zheng, P., Colgan, W.N., Jones, M.G., Yang, D., Kumar, A., Sandhu, J., Schnell, A., Sun, D., et al. (2025). High-resolution spatial mapping of cell state and lineage dynamics in vivo with PEtracer. Science 390, eadx3800. 10.1126/science.adx3800.

44. Anzalone, A.V., Koblan, L.W., and Liu, D.R. (2020). Genome editing with CRISPR–Cas nucleases, base editors, transposases and prime editors. Nat. Biotechnol. 38, 824–844. 10.1038/s41587-020-0561-9.

45. Anzalone, A.V., Randolph, P.B., Davis, J.R., Sousa, A.A., Koblan, L.W., Levy, J.M., Chen, P.J., Wilson, C., Newby, G.A., Raguram, A., et al. (2019). Search-and-replace genome editing without double-strand breaks or donor DNA. Nature 576, 149–157. 10.1038/s41586-019-1711-4.

46. Chen, P.J., Hussmann, J.A., Yan, J., Knipping, F., Ravisankar, P., Chen, P.-F., Chen, C., Nelson, J.W., Newby, G.A., Sahin, M., et al. (2021). Enhanced prime editing systems by manipulating cellular determinants of editing outcomes. Cell 184, 5635–5652.e29. 10.1016/j.cell.2021.09.018.

47. Chen, K.H., Boettiger, A.N., Moffitt, J.R., Wang, S., and Zhuang, X. (2015). Spatially resolved, highly multiplexed RNA profiling in single cells. Science 348, aaa6090. 10.1126/science.aaa6090.

48. Poueymirou, W.T., Auerbach, W., Frendewey, D., Hickey, J.F., Escaravage, J.M., Esau, L., Doré, A.T., Stevens, S., Adams, N.C., Dominguez, M.G., et al. (2007). F0 generation mice fully derived from gene-targeted embryonic stem cells allowing immediate phenotypic analyses. Nat. Biotechnol. 25, 91–99. 10.1038/nbt1263.

49. Snow, M.H.L. (1976). Embryo Growth During the Immediate Postimplantation Period. In Novartis Foundation Symposia, K. Elliott and M. O’Connor, eds. (Wiley), pp. 53–70. 10.1002/9780470720226.ch5.

50. Snow, M.H.L. (1977). Gastrulation in the mouse: Growth and regionalization of the epiblast. Development 42, 293–303. 10.1242/dev.42.1.293.

51. Snow, M.H.L., and Bennett, D. (1978). Gastrulation in the mouse: assessment of cell populations in the epiblast of tw18/tw18 embryos. Development 47, 39–52. 10.1242/dev.47.1.39.

52. Handyside, A.H., and Hunter, S. (1986). Cell division and death in the mouse blastocyst before implantation. Rouxs Arch. Dev. Biol. Off. Organ EDBO 195, 519–526. 10.1007/BF00375893.

53. Yu, Q., Kim, H., Seidel, S., Acosta-Clark, J.F., Martin, B.K., O’Connor, K., Daza, R.M., Gasperini, M., Nathans, J.F., Lam, M., et al. (2026). *In vivo* reconstruction of the cell lineage history of a developing mouse with DNA Typewriter, from zygote to late organogenesis. Preprint at Developmental Biology, 10.64898/2026.07.29.741625.

54. Wong, M.D., van Eede, M.C., Spring, S., Jevtic, S., Boughner, J.C., Lerch, J.P., and Henkelman, R.M. (2015). 4D atlas of the mouse embryo for precise morphological staging. Development 142, 3583–3591. 10.1242/dev.125872.

55. Beddington, R.S., and Robertson, E.J. (1989). An assessment of the developmental potential of embryonic stem cells in the midgestation mouse embryo. Development 105, 733–737. 10.1242/dev.105.4.733.

56. Tam, P.P.L., and Rossant, J. (2003). Mouse embryonic chimeras: tools for studying mammalian development. Development 130, 6155–6163. 10.1242/dev.00893.

57. Fang, W., Bell, C.M., Sapirstein, A., Asami, S., Leeper, K., Zack, D.J., Ji, H., and Kalhor, R. (2022). Quantitative fate mapping: A general framework for analyzing progenitor state dynamics via retrospective lineage barcoding. Cell 185, 4604–4620.e32. 10.1016/j.cell.2022.10.028.

58. Henrique, D., Abranches, E., Verrier, L., and Storey, K.G. (2015). Neuromesodermal progenitors and the making of the spinal cord. Development 142, 2864–2875. 10.1242/dev.119768.

59. Wymeersch, F.J., Wilson, V., and Tsakiridis, A. (2021). Understanding axial progenitor biology in vivo and in vitro. Development 148, dev180612. 10.1242/dev.180612.

60. Gammill, L.S., and Bronner-Fraser, M. (2003). Neural crest specification: migrating into genomics. Nat. Rev. Neurosci. 4, 795–805. 10.1038/nrn1219.

61. Simões-Costa, M., and Bronner, M.E. (2015). Establishing neural crest identity: a gene regulatory recipe. Development 142, 242–257. 10.1242/dev.105445.

62. Soldatov, R., Kaucka, M., Kastriti, M.E., Petersen, J., Chontorotzea, T., Englmaier, L., Akkuratova, N., Yang, Y., Häring, M., Dyachuk, V., et al. (2019). Spatiotemporal structure of cell fate decisions in murine neural crest. Science 364, eaas9536. 10.1126/science.aas9536.

63. Tang, W., and Bronner, M.E. (2020). Neural crest lineage analysis: from past to future trajectory. Development 147, dev193193. 10.1242/dev.193193.

64. Stundl, J., Desingu Rajan, A.R., and Bronner, M.E. (2026). Neural crest gene regulatory networks as drivers of development, diversification and disease. Nat. Rev. Mol. Cell Biol. 27, 467–485. 10.1038/s41580-026-00949-1.

65. Schlosser, G. (2010). Making senses: development of vertebrate cranial placodes. Int. Rev. Cell Mol. Biol. 283, 129–234. 10.1016/S1937-6448(10)83004-7.

66. Kalucka, J., de Rooij, L.P.M.H., Goveia, J., Rohlenova, K., Dumas, S.J., Meta, E., Conchinha, N.V., Taverna, F., Teuwen, L.-A., Veys, K., et al. (2020). Single-Cell Transcriptome Atlas of Murine Endothelial Cells. Cell 180, 764–779.e20. 10.1016/j.cell.2020.01.015.

67. Teillet, M.A., Lapointe, F., and Le Douarin, N.M. (1998). The relationships between notochord and floor plate in vertebrate development revisited. Proc. Natl. Acad. Sci. U. S. A. 95, 11733–11738. 10.1073/pnas.95.20.11733.

68. Cleaver, O., and Krieg, P.A. (2001). Notochord Patterning of the Endoderm. Dev. Biol. 234, 1–12. 10.1006/dbio.2001.0214.

69. Lawson, K.A., and Hage, W.J. (1994). Clonal analysis of the origin of primordial germ cells in the mouse. Ciba Found. Symp. 182, 68–84; discussion 84-91. 10.1002/9780470514573.ch5.

70. Saitou, M., and Yamaji, M. (2012). Primordial germ cells in mice. Cold Spring Harb. Perspect. Biol. 4, a008375. 10.1101/cshperspect.a008375.

71. Placzek, M. (1995). The role of the notochord and floor plate in inductive interactions. Curr. Opin. Genet. Dev. 5, 499–506. 10.1016/0959-437x(95)90055-l.

72. Selleck, M.A., and Stern, C.D. (1991). Fate mapping and cell lineage analysis of Hensen’s node in the chick embryo. Development 112, 615–626. 10.1242/dev.112.2.615.

73. Smith, J.L., Gesteland, K.M., and Schoenwolf, G.C. (1994). Prospective fate map of the mouse primitive streak at 7.5 days of gestation. Dev. Dyn. Off. Publ. Am. Assoc. Anat. 201, 279–289. 10.1002/aja.1002010310.

74. Cai, C.-L., Liang, X., Shi, Y., Chu, P.-H., Pfaff, S.L., Chen, J., and Evans, S. (2003). Isl1 identifies a cardiac progenitor population that proliferates prior to differentiation and contributes a majority of cells to the heart. Dev. Cell 5, 877–889. 10.1016/s1534-5807(03)00363-0.

75. Buckingham, M., Meilhac, S., and Zaffran, S. (2005). Building the mammalian heart from two sources of myocardial cells. Nat. Rev. Genet. 6, 826–835. 10.1038/nrg1710.

76. Meilhac, S.M., and Buckingham, M.E. (2018). The deployment of cell lineages that form the mammalian heart. Nat. Rev. Cardiol. 15, 705–724. 10.1038/s41569-018-0086-9.

77. Kelly, R.G., Brown, N.A., and Buckingham, M.E. (2001). The arterial pole of the mouse heart forms from Fgf10-expressing cells in pharyngeal mesoderm. Dev. Cell 1, 435–440. 10.1016/s1534-5807(01)00040-5.

78. Domínguez, J.N., Meilhac, S.M., Bland, Y.S., Buckingham, M.E., and Brown, N.A. (2012). Asymmetric Fate of the Posterior Part of the Second Heart Field Results in Unexpected Left/Right Contributions to Both Poles of the Heart. Circ. Res. 111, 1323–1335. 10.1161/CIRCRESAHA.112.271247.

79. Tyser, R.C.V., Ibarra-Soria, X., McDole, K., Arcot Jayaram, S., Godwin, J., van den Brand, T.A.H., Miranda, A.M.A., Scialdone, A., Keller, P.J., Marioni, J.C., et al. (2021). Characterization of a common progenitor pool of the epicardium and myocardium. Science 371, eabb2986. 10.1126/science.abb2986.

80. Prummel, K.D., Crowell, H.L., Nieuwenhuize, S., Brombacher, E.C., Daetwyler, S., Soneson, C., Kresoja-Rakic, J., Kocere, A., Ronner, M., Ernst, A., et al. (2022). Hand2 delineates mesothelium progenitors and is reactivated in mesothelioma. Nat. Commun. 13, 1677. 10.1038/s41467-022-29311-7.

81. Moran, H.R., Nyarko, O.O., Garfield, A.L., O’Rourke, R., Ching, R.-C.K., Gray, R., Cobb, T.M., Riemslagh, F.W., Lovely, C.B., Peña, B., et al. (2025). The pericardium forms as a distinct structure during heart formation. Nat. Commun. 16, 8566. 10.1038/s41467-025-63599-5.

82. Kelly, R.G., Buckingham, M.E., and Moorman, A.F. (2014). Heart fields and cardiac morphogenesis. Cold Spring Harb. Perspect. Med. 4, a015750. 10.1101/cshperspect.a015750.

83. de Soysa, T.Y., Ranade, S.S., Okawa, S., Ravichandran, S., Huang, Y., Salunga, H.T., Schricker, A., del Sol, A., Gifford, C.A., and Srivastava, D. (2019). Single-cell analysis of cardiogenesis reveals basis for organ-level developmental defects. Nature 572, 120–124. 10.1038/s41586-019-1414-x.

84. Kelly, R.G., and Buckingham, M.E. (2002). The anterior heart-forming field: voyage to the arterial pole of the heart. Trends Genet. TIG 18, 210–216. 10.1016/s0168-9525(02)02642-2.

85. von Both, I., Silvestri, C., Erdemir, T., Lickert, H., Walls, J.R., Henkelman, R.M., Rossant, J., Harvey, R.P., Attisano, L., and Wrana, J.L. (2004). Foxh1 is essential for development of the anterior heart field. Dev. Cell 7, 331–345. 10.1016/j.devcel.2004.07.023.

86. Zaffran, S., Kelly, R.G., Meilhac, S.M., Buckingham, M.E., and Brown, N.A. (2004). Right ventricular myocardium derives from the anterior heart field. Circ. Res. 95, 261–268. 10.1161/01.RES.0000136815.73623.BE.

87. Stefanovic, S., Barnett, P., van Duijvenboden, K., Weber, D., Gessler, M., and Christoffels, V.M. (2014). GATA-dependent regulatory switches establish atrioventricular canal specificity during heart development. Nat. Commun. 5, 3680. 10.1038/ncomms4680.

88. van Wijk, B., van den Berg, G., Abu-Issa, R., Barnett, P., van der Velden, S., Schmidt, M., Ruijter, J.M., Kirby, M.L., Moorman, A.F.M., and van den Hoff, M.J.B. (2009). Epicardium and myocardium separate from a common precursor pool by crosstalk between bone morphogenetic protein- and fibroblast growth factor-signaling pathways. Circ. Res. 105, 431–441. 10.1161/CIRCRESAHA.109.203083.

89. Später, D., Abramczuk, M.K., Buac, K., Zangi, L., Stachel, M.W., Clarke, J., Sahara, M., Ludwig, A., and Chien, K.R. (2013). A HCN4+ cardiomyogenic progenitor derived from the first heart field and human pluripotent stem cells. Nat. Cell Biol. 15, 1098–1106. 10.1038/ncb2824.

90. Ivanovitch, K., Soro-Barrio, P., Chakravarty, P., Jones, R.A., Bell, D.M., Mousavy Gharavy, S.N., Stamataki, D., Delile, J., Smith, J.C., and Briscoe, J. (2021). Ventricular, atrial, and outflow tract heart progenitors arise from spatially and molecularly distinct regions of the primitive streak. PLoS Biol. 19, e3001200. 10.1371/journal.pbio.3001200.

91. Milgrom-Hoffman, M., Harrelson, Z., Ferrara, N., Zelzer, E., Evans, S.M., and Tzahor, E. (2011). The heart endocardium is derived from vascular endothelial progenitors. Development 138, 4777–4787. 10.1242/dev.061192.

92. Newbern, J.M. (2015). Molecular control of the neural crest and peripheral nervous system development. Curr. Top. Dev. Biol. 111, 201–231. 10.1016/bs.ctdb.2014.11.007.

93. Vega-Lopez, G.A., Cerrizuela, S., and Aybar, M.J. (2017). Trunk neural crest cells: formation, migration and beyond. Int. J. Dev. Biol. 61, 5–15. 10.1387/ijdb.160408gv.

94. Artinger, K.B., and Monsoro-Burq, A.H. (2021). Neural crest multipotency and specification: power and limits of single cell transcriptomic approaches. Fac. Rev. 10, 38. 10.12703/r/10-38.

95. Baggiolini, A., Varum, S., Mateos, J.M., Bettosini, D., John, N., Bonalli, M., Ziegler, U., Dimou, L., Clevers, H., Furrer, R., et al. (2015). Premigratory and migratory neural crest cells are multipotent in vivo. Cell Stem Cell 16, 314–322. 10.1016/j.stem.2015.02.017.

96. Bronner-Fraser, M., and Fraser, S.E. (1988). Cell lineage analysis reveals multipotency of some avian neural crest cells. Nature 335, 161–164. 10.1038/335161a0.

97. Vong, K.I., Alvarez, Y.D., Zhang, Q., Weng, J., Noel, G., Barton, S.T., Chung, C., Howarth, R., Meave, N., Jiwani, F., et al. (2026). Developmental organization of sensory and sympathetic ganglia. Nature 654, 734–743. 10.1038/s41586-026-10313-0.

98. DeTomaso, D., and Yosef, N. (2021). Hotspot identifies informative gene modules across modalities of single-cell genomics. Cell Syst. 12, 446–456.e9. 10.1016/j.cels.2021.04.005.

99. Trimm, E., and Red-Horse, K. (2023). Vascular endothelial cell development and diversity. Nat. Rev. Cardiol. 20, 197–210. 10.1038/s41569-022-00770-1.

100. Ciau-Uitz, A., Monteiro, R., Kirmizitas, A., and Patient, R. (2014). Developmental hematopoiesis: ontogeny, genetic programming and conservation. Exp. Hematol. 42, 669–683. 10.1016/j.exphem.2014.06.001.

101. Nguyen, M.-T., Zhu, J., Nakamura, E., Bao, X., and Mackem, S. (2009). Tamoxifen-dependent, inducible Hoxb6CreERT recombinase function in lateral plate and limb mesoderm, CNS isthmic organizer, posterior trunk neural crest, hindgut, and tailbud. Dev. Dyn. Off. Publ. Am. Assoc. Anat. 238, 467–474. 10.1002/dvdy.21846.

102. Christ, B., Huang, R., and Scaal, M. (2007). Amniote somite derivatives. Dev. Dyn. 236, 2382–2396. 10.1002/dvdy.21189.

103. Pardanaud, L., Luton, D., Prigent, M., Bourcheix, L.M., Catala, M., and Dieterlen-Lievre, F. (1996). Two distinct endothelial lineages in ontogeny, one of them related to hemopoiesis. Development 122, 1363–1371. 10.1242/dev.122.5.1363.

104. Sahai-Hernandez, P., Pouget, C., Eyal, S., Svoboda, O., Chacon, J., Grimm, L., Gjøen, T., and Traver, D. (2023). Dermomyotome-derived endothelial cells migrate to the dorsal aorta to support hematopoietic stem cell emergence. eLife 12, e58300. 10.7554/eLife.58300.

105. Stone, O.A., and Stainier, D.Y.R. (2019). Paraxial Mesoderm Is the Major Source of Lymphatic Endothelium. Dev. Cell 50, 247–255.e3. 10.1016/j.devcel.2019.04.034.

106. Mugford, J.W., Sipilä, P., McMahon, J.A., and McMahon, A.P. (2008). Osr1 expression demarcates a multi-potent population of intermediate mesoderm that undergoes progressive restriction to an Osr1-dependent nephron progenitor compartment within the mammalian kidney. Dev. Biol. 324, 88–98. 10.1016/j.ydbio.2008.09.010.

107. Couly, G., Coltey, P., Eichmann, A., and Le Douarin, N.M. (1995). The angiogenic potentials of the cephalic mesoderm and the origin of brain and head blood vessels. Mech. Dev. 53, 97–112. 10.1016/0925-4773(95)00428-9.

108. Lagha, M., Brunelli, S., Messina, G., Cumano, A., Kume, T., Relaix, F., and Buckingham, M.E. (2009). Pax3:Foxc2 Reciprocal Repression in the Somite Modulates Muscular versus Vascular Cell Fate Choice in Multipotent Progenitors. Dev. Cell 17, 892–899. 10.1016/j.devcel.2009.10.021.

109. Cao, X., Mircea, M., Cascione, S., Amel, A., Tsikari, T., van den Hil, F.E., Mei, H., Neumann, K., Alemany, A., Anastassiadis, K., et al. (2025). Tissue microenvironment dictates the state of human iPSC-derived endothelial cells of distinct developmental origin in 3D cardiac microtissues. iScience 28, 113611. 10.1016/j.isci.2025.113611.

110. Zirra, A., Wiethoff, S., and Patani, R. (2016). Neural Conversion and Patterning of Human Pluripotent Stem Cells: A Developmental Perspective. Stem Cells Int. 2016, 8291260. 10.1155/2016/8291260.

111. Brooks, E.R., Moorman, A.R., Bhattacharya, B., Prudhomme, I.S., Land, M., Alcorn, H.L., Sharma, R., Pe’er, D., and Zallen, J.A. (2025). A single-cell atlas of spatial and temporal gene expression in the mouse cranial neural plate. ELife 13 RP102819. https://doi.org/DOI%2010.7554/eLife.102819.3.

112. Duboule, D. (2007). The rise and fall of Hox gene clusters. Development 134, 2549–2560. 10.1242/dev.001065.

113. Wellik, D.M. (2007). Hox patterning of the vertebrate axial skeleton. Dev. Dyn. Off. Publ. Am. Assoc. Anat. 236, 2454–2463. 10.1002/dvdy.21286.

114. Chen, A., Liao, S., Cheng, M., Ma, K., Wu, L., Lai, Y., Qiu, X., Yang, J., Xu, J., Hao, S., et al. (2022). Spatiotemporal transcriptomic atlas of mouse organogenesis using DNA nanoball-patterned arrays. Cell 185, 1777–1792.e21. 10.1016/j.cell.2022.04.003.

115. Cajal, M., Lawson, K.A., Hill, B., Moreau, A., Rao, J., Ross, A., Collignon, J., and Camus, A. (2012). Clonal and molecular analysis of the prospective anterior neural boundary in the mouse embryo. Development 139, 423–436. 10.1242/dev.075499.

116. Metzis, V., Steinhauser, S., Pakanavicius, E., Gouti, M., Stamataki, D., Ivanovitch, K., Watson, T., Rayon, T., Mousavy Gharavy, S.N., Lovell-Badge, R., et al. (2018). Nervous System Regionalization Entails Axial Allocation before Neural Differentiation. Cell 175, 1105–1118.e17. 10.1016/j.cell.2018.09.040.

117. Placzek, M., and Furley, A. (1996). Neural development: Patterning cascades in the neural tube. Curr. Biol. 6, 526–529. 10.1016/s0960-9822(02)00533-x.

118. Zervas, M., Millet, S., Ahn, S., and Joyner, A.L. (2004). Cell Behaviors and Genetic Lineages of the Mesencephalon and Rhombomere 1. Neuron 43, 345–357. 10.1016/j.neuron.2004.07.010.

119. Kim, Y.-I., O’Rourke, R., and Sagerström, C.G. (2023). scMultiome analysis identifies embryonic hindbrain progenitors with mixed rhombomere identities. eLife 12, e87772. 10.7554/eLife.87772.

120. Krumlauf, R., and Wilkinson, D.G. (2021). Segmentation and patterning of the vertebrate hindbrain. Development 148, dev186460. 10.1242/dev.186460.

121. Nonchev, S., Vesque, C., Maconochie, M., Seitanidou, T., Ariza-McNaughton, L., Frain, M., Marshall, H., Sham, M.H., Krumlauf, R., and Charnay, P. (1996). Segmental expression of Hoxa-2 in the hindbrain is directly regulated by Krox-20. Development 122, 543–554. 10.1242/dev.122.2.543.

122. Rhinn, M., and Brand, M. (2001). The midbrain--hindbrain boundary organizer. Curr. Opin. Neurobiol. 11, 34–42. 10.1016/s0959-4388(00)00171-9.

123. Watson, C., Bartholomaeus, C., and Puelles, L. (2019). Time for Radical Changes in Brain Stem Nomenclature-Applying the Lessons From Developmental Gene Patterns. Front. Neuroanat. 13, 10. 10.3389/fnana.2019.00010.

124. Forlani, S., Lawson, K.A., and Deschamps, J. (2003). Acquisition of Hox codes during gastrulation and axial elongation in the mouse embryo. Development 130, 3807–3819. 10.1242/dev.00573.

125. Wymeersch, F.J., Skylaki, S., Huang, Y., Watson, J.A., Economou, C., Marek-Johnston, C., Tomlinson, S.R., and Wilson, V. (2018). Transcriptionally dynamic progenitor populations organised around a stable niche drive axial patterning. Development, dev.168161. 10.1242/dev.168161.

126. Rodrigo Albors, A., Halley, P.A., and Storey, K.G. (2018). Lineage tracing of axial progenitors using Nkx1-2CreERT2 mice defines their trunk and tail contributions. Development 145, dev164319. 10.1242/dev.164319.

127. Wymeersch, F.J., Huang, Y., Blin, G., Cambray, N., Wilkie, R., Wong, F.C.K., and Wilson, V. (2016). Position-dependent plasticity of distinct progenitor types in the primitive streak. eLife 5, e10042. 10.7554/eLife.10042.

128. Attardi, A., Fulton, T., Florescu, M., Shah, G., Muresan, L., Lenz, M.O., Lancaster, C., Huisken, J., van Oudenaarden, A., and Steventon, B. (2018). Neuromesodermal progenitors are a conserved source of spinal cord with divergent growth dynamics. Development 145, dev166728. 10.1242/dev.166728.

129. Guillot, C., Djeffal, Y., Michaut, A., Rabe, B., and Pourquié, O. (2021). Dynamics of primitive streak regression controls the fate of neuromesodermal progenitors in the chicken embryo. eLife 10, e64819. 10.7554/eLife.64819.

130. Burke, A.C., Nelson, C.E., Morgan, B.A., and Tabin, C. (1995). Hox genes and the evolution of vertebrate axial morphology. Development 121, 333–346. 10.1242/dev.121.2.333.

131. Kretzschmar, K., and Watt, F.M. (2012). Lineage tracing. Cell 148, 33–45. 10.1016/j.cell.2012.01.002.

132. Stadler, T., Pybus, O.G., and Stumpf, M.P.H. (2021). Phylodynamics for cell biologists. Science 371, eaah6266. 10.1126/science.aah6266.

133. Sankaran, V.G., Weissman, J.S., and Zon, L.I. (2022). Cellular barcoding to decipher clonal dynamics in disease. Science 378, eabm5874. 10.1126/science.abm5874.

134. Mintz, B. (1967). Gene control of mammalian pigmentary differentiation. I. Clonal origin of melanocytes. Proc. Natl. Acad. Sci. U. S. A. 58, 344–351. 10.1073/pnas.58.1.344.

135. Tam, P.P., and Zhou, S.X. (1996). The allocation of epiblast cells to ectodermal and germ-line lineages is influenced by the position of the cells in the gastrulating mouse embryo. Dev. Biol. 178, 124–132. 10.1006/dbio.1996.0203.

136. Kinder, S.J., Tsang, T.E., Quinlan, G.A., Hadjantonakis, A.-K., Nagy, A., and Tam, P.P.L. (1999). The orderly allocation of mesodermal cells to the extraembryonic structures and the anteroposterior axis during gastrulation of the mouse embryo. Development 126, 4691–4701. 10.1242/dev.126.21.4691.

137. Lu, C.C., and Robertson, E.J. (2004). Multiple roles for Nodal in the epiblast of the mouse embryo in the establishment of anterior-posterior patterning. Dev. Biol. 273, 149–159. 10.1016/j.ydbio.2004.06.004.

138. Gardner, R.L., and Cockroft, D.L. (1998). Complete dissipation of coherent clonal growth occurs before gastrulation in mouse epiblast. Development 125, 2397–2402. 10.1242/dev.125.13.2397.

139. Cambray, N., and Wilson, V. (2002). Axial progenitors with extensive potency are localised to the mouse chordoneural hinge. Development 129, 4855–4866. 10.1242/dev.129.20.4855.

140. Cambray, N., and Wilson, V. (2007). Two distinct sources for a population of maturing axial progenitors. Development 134, 2829–2840. 10.1242/dev.02877.

141. Lin, L., Zhong, J., Jiang, F., Wang, Y., Ma, L., Yang, J., Li, Y., Gao, R.-R., Feng, H., Cai, B., et al. (2026). A single-cell time-series atlas of endothelial cell embryonic development. Cell 189, 1573–1590.e24. 10.1016/j.cell.2026.01.002.

142. Weinreb, C., Wolock, S., Tusi, B.K., Socolovsky, M., and Klein, A.M. (2018). Fundamental limits on dynamic inference from single-cell snapshots. Proc. Natl. Acad. Sci. U. S. A. 115, E2467–E2476. 10.1073/pnas.1714723115.

143. Schiebinger, G., Shu, J., Tabaka, M., Cleary, B., Subramanian, V., Solomon, A., Gould, J., Liu, S., Lin, S., Berube, P., et al. (2019). Optimal-Transport Analysis of Single-Cell Gene Expression Identifies Developmental Trajectories in Reprogramming. Cell 176, 928–943.e22. 10.1016/j.cell.2019.01.006.

144. Blanpain, C., and Simons, B.D. (2013). Unravelling stem cell dynamics by lineage tracing. Nat. Rev. Mol. Cell Biol. 14, 489–502. 10.1038/nrm3625.

145. Wells, J.M., and Watt, F.M. (2018). Diverse mechanisms for endogenous regeneration and repair in mammalian organs. Nature 557, 322–328. 10.1038/s41586-018-0073-7.

146. Martinez Arias, A., Nichols, J., and Schröter, C. (2013). A molecular basis for developmental plasticity in early mammalian embryos. Development 140, 3499–3510. 10.1242/dev.091959.

147. Wolf, F.A., Angerer, P., and Theis, F.J. (2018). SCANPY: large-scale single-cell gene expression data analysis. Genome Biol. 19, 15. 10.1186/s13059-017-1382-0.

148. Lopez, R., Regier, J., Cole, M.B., Jordan, M.I., and Yosef, N. (2018). Deep generative modeling for single-cell transcriptomics. Nat. Methods 15, 1053–1058. 10.1038/s41592-018-0229-2.

149. Dicks, S., Heumos, L., May, L., Jimenez, S., Angerer, P., Gold, I., Virshup, I., Fischer, F., Gill, M., Boerries, M., et al. (2026). GPU-accelerated single-cell analysis at scale with rapids-singlecell. Preprint at arXiv, 10.48550/arXiv.2603.02402 https://doi.org/10.48550/arXiv.2603.02402.

150. Pedregosa, F., Varoquaux, G., Gramfort, A., Michel, V., Thirion, B., Grisel, O., Blondel, M., Müller, A., Nothman, J., Louppe, G., et al. (2012). Scikit-learn: Machine Learning in Python. 10.48550/ARXIV.1201.0490.

151. Jones, M.G., Khodaverdian, A., Quinn, J.J., Chan, M.M., Hussmann, J.A., Wang, R., Xu, C., Weissman, J.S., and Yosef, N. (2020). Inference of single-cell phylogenies from lineage tracing data using Cassiopeia. Genome Biol. 21, 92. 10.1186/s13059-020-02000-8.

152. Price, M.N., Dehal, P.S., and Arkin, A.P. (2010). FastTree 2--approximately maximum-likelihood trees for large alignments. PLoS One 5, e9490. 10.1371/journal.pone.0009490.

153. Prillo, S., Ravoor, A., Yosef, N., and Song, Y.S. (2026). ConvexML: Fast and accurate branch length estimation under irreversible mutation models, illustrated through applications to CRISPR/Cas9-based lineage tracing. Syst. Biol. 75, 115–134. 10.1093/sysbio/syaf054.

154. Chu, G., Schmidt, H., and Raphael, B.J. (2026). LAML-Pro: Joint Maximum Likelihood Inference of Cell Genotypes and Cell Lineage Trees. 10.1093/bioinformatics/btag219.

155. Harris, C.R., Millman, K.J., Van Der Walt, S.J., Gommers, R., Virtanen, P., Cournapeau, D., Wieser, E., Taylor, J., Berg, S., Smith, N.J., et al. (2020). Array programming with NumPy. Nature 585, 357–362. 10.1038/s41586-020-2649-2.

156. McKinney, W. (2010). Data Structures for Statistical Computing in Python. In, pp. 56–61. 10.25080/Majora-92bf1922-00a.

157. Hunter, J.D. (2007). Matplotlib: A 2D Graphics Environment. Comput. Sci. Eng. 9, 90–95. 10.1109/MCSE.2007.55.

158. Waskom, M. (2021). seaborn: statistical data visualization. J. Open Source Softw. 6, 3021. 10.21105/joss.03021.

159. Paszke, A., Gross, S., Massa, F., Lerer, A., Bradbury, J., Chanan, G., Killeen, T., Lin, Z., Gimelshein, N., Antiga, L., et al. (2019). PyTorch: An Imperative Style, High-Performance Deep Learning Library. Preprint at arXiv, 10.48550/ARXIV.1912.01703 https://doi.org/10.48550/ARXIV.1912.01703.

160. Platt, R.J., Chen, S., Zhou, Y., Yim, M.J., Swiech, L., Kempton, H.R., Dahlman, J.E., Parnas, O., Eisenhaure, T.M., Jovanovic, M., et al. (2014). CRISPR-Cas9 knockin mice for genome editing and cancer modeling. Cell 159, 440–455. 10.1016/j.cell.2014.09.014.

161. Uenaka, T., Napole, A., Saha, A.D., Sun, D., Singavarapu, A., Calzada, E., Chen, J., Erlebach, L., McQuade, A., Ramos, D.M., et al. (2026). Prevention of transgene silencing during human pluripotent stem cell differentiation. Cell Stem Cell 33, 517–530.e8. 10.1016/j.stem.2026.01.007.

162. Markoulaki, S., Meissner, A., and Jaenisch, R. (2008). Somatic cell nuclear transfer and derivation of embryonic stem cells in the mouse. Methods 45, 101–114. 10.1016/j.ymeth.2008.04.002.

163. Eggan, K., Akutsu, H., Loring, J., Jackson-Grusby, L., Klemm, M., Rideout, W.M., Yanagimachi, R., and Jaenisch, R. (2001). Hybrid vigor, fetal overgrowth, and viability of mice derived by nuclear cloning and tetraploid embryo complementation. Proc. Natl. Acad. Sci. U. S. A. 98, 6209–6214. 10.1073/pnas.101118898.

164. Hanna, J., Markoulaki, S., Mitalipova, M., Cheng, A.W., Cassady, J.P., Staerk, J., Carey, B.W., Lengner, C.J., Foreman, R., Love, J., et al. (2009). Metastable pluripotent states in NOD-mouse-derived ESCs. Cell Stem Cell 4, 513–524. 10.1016/j.stem.2009.04.015.

165. Matsuda, T., and Cepko, C.L. (2007). Controlled expression of transgenes introduced by in vivo electroporation. Proc. Natl. Acad. Sci. U. S. A. 104, 1027–1032. 10.1073/pnas.0610155104.

166. Bolondi, A., Law, B.K., Kretzmer, H., Gassaloglu, S.I., Buschow, R., Riemenschneider, C., Yang, D., Walther, M., Veenvliet, J.V., Meissner, A., et al. (2024). Reconstructing axial progenitor field dynamics in mouse stem cell-derived embryoids. Dev. Cell 59, 1489–1505.e14. 10.1016/j.devcel.2024.03.024.

167. Eakin, G.S., and Hadjantonakis, A.-K. (2006). Production of chimeras by aggregation of embryonic stem cells with diploid or tetraploid mouse embryos. Nat. Protoc. 1, 1145–1153. 10.1038/nprot.2006.173.

168. Humięcka, M., Krupa, M., Guzewska, M.M., Maleszewski, M., and Suwińska, A. (2016). ESCs injected into the 8-cell stage mouse embryo modify pattern of cleavage and cell lineage specification. Mech. Dev. 141, 40–50. 10.1016/j.mod.2016.06.002.

169. Strawbridge, S.E., Schrattel, A.K., Humphreys, P., Jones, K.A., Artus, J., Hadjantonakis, A.-K., Fletcher, A.G., and Nichols, J. (2025). Donor embryonic stem cells displace host cells of 8-cell-stage chimeras to the extra-embryonic lineages by spatial crowding and FGF4 signalling. Development 152, dev204518. 10.1242/dev.204518.

170. Gong, W., Granados, A.A., Hu, J., Jones, M.G., Raz, O., Salvador-Martínez, I., Zhang, H., Chow, K.-H.K., Kwak, I.-Y., Retkute, R., et al. (2021). Benchmarked approaches for reconstruction of in vitro cell lineages and in silico models of C. elegans and M. musculus developmental trees. Cell Syst. 12, 810–826.e4. 10.1016/j.cels.2021.05.008.

171. Feydy, J., Roussillon, P., Trouvé, A., and Gori, P. (2021). Fast and Scalable Optimal Transport for Brain Tractograms. Preprint at arXiv, 10.48550/arXiv.2107.02010 https://doi.org/10.48550/arXiv.2107.02010.

172. Tirosh, I., Izar, B., Prakadan, S.M., Wadsworth, M.H., Treacy, D., Trombetta, J.J., Rotem, A., Rodman, C., Lian, C., Murphy, G., et al. (2016). Dissecting the multicellular ecosystem of metastatic melanoma by single-cell RNA-seq. Science 352, 189–196. 10.1126/science.aad0501.

173. Lein, E.S., Hawrylycz, M.J., Ao, N., Ayres, M., Bensinger, A., Bernard, A., Boe, A.F., Boguski, M.S., Brockway, K.S., Byrnes, E.J., et al. (2007). Genome-wide atlas of gene expression in the adult mouse brain. Nature 445, 168–176. 10.1038/nature05453.

174. Harris, J.A., Mihalas, S., Hirokawa, K.E., Whitesell, J.D., Choi, H., Bernard, A., Bohn, P., Caldejon, S., Casal, L., Cho, A., et al. (2019). Hierarchical organization of cortical and thalamic connectivity. Nature 575, 195–202. 10.1038/s41586-019-1716-z.

## Supplemental References

1. Price, M.N., Dehal, P.S., and Arkin, A.P. (2010). FastTree 2--approximately maximum-likelihood trees for large alignments. PLoS One 5, e9490. 10.1371/journal.pone.0009490.

2. Chu, G., Schmidt, H., and Raphael, B.J. (2026). LAML-Pro: Joint Maximum Likelihood Inference of Cell Genotypes and Cell Lineage Trees. 10.1093/bioinformatics/btag219.

3. Jones, M.G., Khodaverdian, A., Quinn, J.J., Chan, M.M., Hussmann, J.A., Wang, R., Xu, C., Weissman, J.S., and Yosef, N. (2020). Inference of single-cell phylogenies from lineage tracing data using Cassiopeia. Genome Biol. 21, 92. 10.1186/s13059-020-02000-8.

4. Prillo, S., Ravoor, A., Yosef, N., and Song, Y.S. (2026). ConvexML: Fast and accurate branch length estimation under irreversible mutation models, illustrated through applications to CRISPR/Cas9-based lineage tracing. Syst. Biol. 75, 115–134. 10.1093/sysbio/syaf054.

5. Bondarenko, V., Nikolaev, M., Kromm, D., Belousov, R., Wolny, A., Blotenburg, M., Zeller, P., Rezakhani, S., Hugger, J., Uhlmann, V., et al. (2023). Embryo-uterine interaction coordinates mouse embryogenesis during implantation. EMBO J. 42, e113280. 10.15252/embj.2022113280.

6. Pijuan-Sala, B., Griffiths, J.A., Guibentif, C., Hiscock, T.W., Jawaid, W., Calero-Nieto, F.J., Mulas, C., Ibarra-Soria, X., Tyser, R.C.V., Ho, D.L.L., et al. (2019). A single-cell molecular map of mouse gastrulation and early organogenesis. Nature 566, 490–495. 10.1038/s41586-019-0933-9.

7. Qiu, C., Martin, B.K., Welsh, I.C., Daza, R.M., Le, T.-M., Huang, X., Nichols, E.K., Taylor, M.L., Fulton, O., O’Day, D.R., et al. (2024). A single-cell time-lapse of mouse prenatal development from gastrula to birth. Nature 626, 1084–1093. 10.1038/s41586-024-07069-w.

